# Comparison of individualized behavioral predictions across anatomical, diffusion and functional connectivity MRI

**DOI:** 10.1101/2022.03.08.483564

**Authors:** Leon Qi Rong Ooi, Jianzhong Chen, Zhang Shaoshi, Ru Kong, Angela Tam, Jingwei Li, Elvisha Dhamala, Juan Helen Zhou, Avram J Holmes, B. T. Thomas Yeo

## Abstract

A fundamental goal across the neurosciences is the characterization of relationships linking brain anatomy, functioning, and behavior. Although various MRI modalities have been developed to probe these relationships, direct comparisons of their ability to predict behavior have been lacking. Here, we compared the ability of anatomical T1, diffusion and functional MRI (fMRI) to predict behavior at an individual level. Cortical thickness, area and volume were extracted from anatomical T1 images. Diffusion Tensor Imaging (DTI) and approximate Neurite Orientation Dispersion and Density Imaging (NODDI) models were fitted to the diffusion images. The resulting metrics were projected to the Tract-Based Spatial Statistics (TBSS) skeleton. We also ran probabilistic tractography for the diffusion images, from which we extracted the stream count, average stream length, and the average of each DTI and NODDI metric across tracts connecting each pair of brain regions. Functional connectivity (FC) was extracted from both task and resting-state fMRI. Individualized prediction of a wide range of behavioral measures were performed using kernel ridge regression, linear ridge regression and elastic net regression. Consistency of the results were investigated with the Human Connectome Project (HCP) and Adolescent Brain Cognitive Development (ABCD) datasets. In both datasets, FC-based models gave the best prediction performance, regardless of regression model or behavioral measure. This was especially true for the cognitive domain. Furthermore, all modalities were able to predict cognition better than other behavioral domains. Combining all modalities improved prediction of cognition, but not other behavioral domains. Finally, across all behaviors, combining resting and task FC yielded prediction performance similar to combining all modalities. Overall, our study suggests that in the case of healthy children and young adults, behaviorally-relevant information in T1 and diffusion features might reflect a subset of the variance captured by FC.

**Highlights:**

- FC predicts behavior better than anatomical and diffusion features
- Cognition is predicted better than other behavioral domains regardless of modality
- Combining resting & task FC improves prediction as much as combining all modalities
- Findings were replicated over 3 regression models and 2 datasets

## 1. Introduction

A fundamental aim of neuroscience is to answer how brain characteristics are linked to behavior (Zatorre et al., 2012; Rosenberg et al., 2018). Previous studies have established that inter-individual variation in functional and structural patterns covary with behavioral and demographical traits (Finn et al., 2015; Smith et al., 2015; Llera et al., 2019; Alnæs et al., 2020). In recent years, there is an increasing interest in utilizing machine learning algorithms to predict behavioral traits at an individual level (Bzdok & Meyer-Lindenberg, 2018; Calhoun, 2018). Here, we compare the ability of anatomical, diffusion and functional characteristics of the brain in making individualized predictions of behavioral traits.

Diffusion and anatomical MRI have been used to make individualized predictions in a large variety of neurological and psychiatric disorders (Sabuncu & Konukoglu, 2015; Arbabshirani et al., 2017; Bajaj et al., 2017; Cohen et al., 2021; Elad et al., 2021). However, their utility for behavioral predictions in healthy participants has been less explored. Given that psychiatric symptoms and associated shifts in brain function likely exist on a spectrum from healthy participants to patient populations (Xia et al., 2018; Kebets et al., 2019; Peter et al., 2021), predicting behavioral traits in the former group is an important endeavour (Lui et al., 2016). Functional connectivity has already been widely used to predict individual behavioral traits in healthy participants (Kong et al., 2019; Li et al., 2019; Cai et al., 2020; Chen et al., 2020; He et al., 2020; Sripada et al., 2020). However, similar work utilizing anatomical (Lu et al., 2014; Avinun et al., 2020; Liu et al., 2021) and diffusion MRI (Lewis et al., 2016; Mansour et al., 2021) has been a lot more sparse. Furthermore, most of these studies have performed predictions using a single modality, so the comparative value of each modality in making individualized predictions is unclear.

Several recent studies have tackled the topic of comparing MRI modalities for behavioral prediction (Dhamala et al., 2021; Mansour et al., 2021; Rasero et al., 2021). However, their analyses were performed in the Human Connectome Project (HCP), which is perhaps the most widely used dataset for studies investigating individualized predictions in healthy participants (Finn et al., 2015; Greene et al., 2018; Gao et al., 2019). Repeated use of the HCP for investigating behavior prediction leads to the issue of dataset decay (Thompson et al., 2020). The over-reliance on the dataset results in increased possibility of type I errors as the number of sequential tests on the dataset increases (Thompson et al., 2020). Furthermore, repeated use of the training and test sets from the same dataset leads to overly optimistic prediction results with models less able to generalize to new datasets (Recht et al., 2019; Beyer et al., 2020). These considerations highlight the need for additional analyses of independent data and/or less utilized datasets to replicate the conclusions. Therefore, in the current study, in addition to the widely used HCP dataset, we utilized the adolescent brain cognitive development (ABCD) dataset.

Given that different MRI modalities measure different aspects of the brain biology, one question is whether combining multiple MRI modalities might improve behavioral prediction. Rasero and colleagues found that integrating diffusion MRI and resting FC led to improvement in predicting cognition (Rasero et al., 2021). However, two other studies did not find any benefit from integrating diffusion MRI, resting-state fMRI and/or anatomical features to predict cognition (Dhamala et al., 2021; Xiao et al., 2021). Overall, the literature is inconsistent about the value of integrating multiple modalities. Furthermore, despite the wide range of possible diffusion features, most studies only focused on one particular type of diffusion feature. Most studies have also focused on predicting a small number of behavioral measures (e.g. cognition), which reduces their generalizability to other behavior.

In this study, we compared the utility of different MRI modalities for behavioral prediction across a wide range of behavioral measures in two large datasets (HCP and ABCD) using three different regression models. Unlike previous studies, we considered a wide range of diffusion features, including fractional anisotropy (FA), mean diffusivity (MD), axial diffusivity (AD) and radial diffusivity (RD). An approximate Neurite Orientation Dispersion and Density Imaging (AMICO-NODDI) model was also used to derive orientation dispersion (OD), intracellular volume fraction (ICVF), and isotropic volume fraction (ISOVF) features. Probabilistic tractography was performed to extract structural connectivity (SC) features. Furthermore, unlike most previous studies on multi-modal prediction, we considered both resting and task FC. In the case of anatomical T1, we considered cortical thickness, volume and surface area. We also combined features within and across modalities to investigate whether integrating modalities resulted in improved prediction.

## 2. Methods and materials

### 2.1. Datasets and participants

We considered participants from the HCP WU-Minn S1200 release. After strict pre-processing quality control of imaging data, we filtered participants from Li’s set of 953 participants (Li et al., 2019) based on the availability of a complete set of structural, diffusion and functional (resting and task) scans, as well as all behavioral scores of interest (Table S1). Our main analysis comprised 753 participants, who fulfilled all selection criteria.

We also considered participants from the ABCD 2.0.1 release. After strict pre-processing quality control of imaging data, participants from Chen’s set of 2262 subjects who underwent motion-filtering to remove pseudo-respiratory motion (Chen et al., 2020) were filtered based on the availability of a complete set of structural, diffusion and functional (resting and task) scans, and all behavioral scores of interest (Table S2). We also excluded participants from sites which used Phillips scanners due to incorrect processing, as recommended by the ABCD consortium. Our main analysis comprised 1823 participants, who fulfilled all selection criteria.

### 2.2. Imaging acquisition and processing

Minimally processed T1 and multi-shell diffusion from each dataset were utilized. Details about the acquisition protocol and minimal processing for the HCP data can be found elsewhere (Glasser et al., 2013; Van Essen et al., 2013). Likewise, acquisition protocol and minimal processing pipelines for the ABCD can be found elsewhere (Casey et al., 2018; Hagler et al., 2019).

FMRI data in the HCP included working memory, gambling, motor, language and social cognition tasks, as well as the resting-state scans. We excluded the relational processing and emotional processing tasks in the HCP as the run duration for these tasks were below 3 minutes. The MSMAll ICA-FIX data was used for the resting state scans, and the MSMAll data was used for task fMRI (Glasser et al., 2013). Global signal regression has been shown to improve behavioral prediction (Li et al., 2019), so we further applied global signal regression (GSR) and censoring, consistent with our previous studies (Li et al., 2019; He et al., 2020; Kong et al., 2021). More details of the processing can be found elsewhere (Li et al., 2019).

For the ABCD study, fMRI data included the N-back, monetary incentive delay (MID), stop signal task (SST), as well as resting-state scans. The minimally processed functional data were utilized (Hagler et al., 2019). We additionally aligned the functional images to the T1 images using boundary-based registration, and performed motion filtering, nuisance regression, GSR, censoring and bandpass filtering. The data was then projected onto FreeSurfer fsaverage6 surface space and smoothed using a 6 mm full-width half maximum kernel. More details of the processing can be found elsewhere (Chen et al., 2020).

### 2.3. Imaging features for behavioral prediction

#### 2.3.1. Anatomical feature processing

The 400-region Schaefer parcellation was projected to each participant’s native surface space (Schaefer et al., 2018). Using each participant’s T1 image, cortical volume, cortical thickness and cortical area were extracted from each of the 400 regions of interest (ROIs) using Freesurfer 5.3.0 (Dale et al., 1999). Cortical volumes were divided by intra-cranial volume (ICV), while cortical area was divided by ICV^2/3^. This resulted in three 400 x participants feature matrices for each dataset.

#### 2.3.2. Diffusion feature processing

A diffusion tensor model (DTI) was fitted to each participant’s diffusion images using FSL’s DTIFIT (Basser et al., 1994). The fractional anisotropy (FA), mean diffusivity (MD), axial diffusivity (AD) and radial diffusivity (RD) images were generated for each participant. Additionally, a relaxed Neurite Orientation Dispersion and Density Imaging (AMICO-NODDI) model was also fitted to the diffusion images (Daducci et al., 2015). Orientation dispersion (OD), intracellular volume fraction (ICVF), and isotropic volume fraction (ISOVF) images were generated for each participant.

The diffusion features were further processed in two ways. First, a TBSS skeleton was generated for each set of participants (one for HCP and one for ABCD), and the seven diffusion metric images (FA, MD, AD, RD, OD, ICVF, ISOVF) were projected to the skeleton (Smith et al., 2006). The voxels of each diffusion metric skeleton were vectorized for each participant, yielding seven feature matrices for each dataset. Each matrix is of size number of TBSS voxels x number of participants.

Secondly, probabilistic tractography was run for each participant using MRtrix (Tournier et al., 2019). The 400-region Schaefer parcellation was projected to each participant’s native surface space (Schaefer et al., 2018). Nine 400 x 400 structural connectivity (SC) matrices were generated. The first matrix was a symmetric matrix containing the log transformation of stream count connecting each ROI pair. The second matrix comprised the average length of streams. The final seven matrices corresponded to the seven diffusion metrics averaged along and across streams connecting each ROI pair. The lower triangle of each matrix was vectorized for each participant, yielding nine 79,800 x number of participants feature matrices for each dataset.

#### 2.3.3. Functional feature processing

A functional connectivity (FC) matrix was generated for each task fMRI and resting-state fMRI scan using the 400-region Schaefer parcellation. The FC matrix was constructed by computing the Pearson’s correlation between the fMRI signals of each ROI pair. The lower triangle of each matrix was vectorized for each participant, yielding six feature matrices for the HCP and four feature matrices for the ABCD study.

### 2.4. Behavioral data

We analyzed 58 behavioral scores from the HCP, consistent with our previous studies (Kong et al., 2019; Li et al., 2019). In the case of ABCD, we considered 36 behavioral scores, consistent with our previous study (Chen et al., 2020). A complete list of scores used for the HCP and ABCD can be found in Tables S1 and S2 respectively.

Because many behavioral scores were correlated, we performed a factor analysis within each dataset to derive components explaining differing aspects of behavior. The scores from all participants with a full set of scores (even if they were missing imaging data) from each dataset underwent a principal component analysis. The top three components explaining the most variance were retained and entered into varimax rotation (Kaiser, 1958). In the HCP, based on the behavioral loadings (Table S3), we interpreted the three components to be related to (1) cognition, (2) life dissatisfaction and (3) emotional recognition. In the ABCD, based on the behavioral loadings (Table S4), we interpreted the three components to the 3 components to be related to (1) cognition, (2) personality and (3) mental health.

Some studies have suggested that in the context of phenotypic prediction, the behavioral component scores should be estimated from the training data and then applied to the test data. However, such a procedure would result in a significantly more complex workflow. To ensure our conclusions were not biased by the estimation of behavioral components from the full dataset, we also considered prediction results from individual behavioral scores (58 measures in HCP and 36 measures in ABCD).

### 2.5. Single-feature-type prediction models

We utilized different regression models to predict the 3 behavioral components and each behavioral measure in each dataset. Our main analysis utilized kernel ridge regression (KRR), which has shown strong behavioral prediction performance (Kong et al., 2019; Chen et al., 2020; He et al., 2020). Briefly, KRR performs predictions based on the similarity between imaging features. A L2-regularization term was used in the model to reduce overfitting.

A separate predictive model was built for each feature type within each MRI modality. In the case of anatomical features, three KRR models were evaluated for each behavioral measure, corresponding to cortical volume, thickness and area. In the case of TBSS, seven KRR models were evaluated for each behavioral measure, corresponding to FA, MD, AD, RD, OD, ISOVF and ICVF. In the case of structural connectivity, nine KRR models were evaluated for each behavioral measure, corresponding to the log transformation of stream counts, stream length, FA, MD, AD, RD, OD, ISOVF and ICVF. In the case of FC in HCP, six KRR models were evaluated for each behavioral measure, corresponding to resting FC and five different tasks. In the case of FC in ABCD, four KRR models were evaluated for each behavioral measure, corresponding to resting FC and three different tasks.

Each regression model was trained using a nested cross-validation procedure. In the HCP, we performed 60 random replications of 10-fold nested cross-validation. The family structure was taken into account when performing the cross-validation – participants from the same family were placed into either the test fold or training folds, but not split across training and test folds.

In the case of ABCD, similar to our previous study (Chen et al., 2020), we combined participants across the 22 imaging sites, yielding 10 “site-clusters”. Each site-cluster comprised at least 140 individuals (see Table S5). We then performed a leave-3-site-clusters out nested cross-validation – 7 random site-clusters were used for training while the remaining 3 site-clusters were used for testing. The prediction was performed for every possible split of the site clusters, resulting in 120 replications.

Age and sex were regressed from the behavioral measures. Regression was performed on the training folds and the regression coefficients were applied to the test fold. Accuracy of each model was defined as the correlation between the predicted scores of the test participants and their actual scores within each test fold, and then averaged across test folds and replications. We additionally computed accuracy using the coefficient of determination (COD).

To ensure our conclusions are across different regression approaches, we also considered linear ridge regression (LRR) and elastic net regression (Friedman et al., 2010).

### 2.6. Multiple-feature-type prediction models

To combine across features, we applied a stacking procedure. For each participant, predictions from the single-feature-type KRR models (first-level predictions) were concatenated into a vector and used as prediction features in a 2nd level linear regression with no regularization. We also considered the use of multi-kernel ridge regression (multi-KRR), which we have previously utilized to predict behavioral measures using task and resting FC.

Overall, we trained three models: a multi-KRR model combining all FC features, a stacking model combining all FC-based models, and a stacking model combining all single-feature-type models from all modalities. We note that we did not consider a multi-KRR model combining all features from all modalities because that was too computationally expensive.

In the case of the stacking, to prevent data leakage between the training and test folds, cross-validation splits were fixed from the single-feature-type models. The training data consisted of first-level predictions made by the “inner-loops” of the first level models so that none of the first-level predictions would have been made from participants of the test folds. Similar to the single-modality models, prediction performance was again evaluated using Pearson’s correlation and COD.

### 2.7. Statistical tests

To test whether a model performed better than chance, we performed a permutation test by shuffling behavioral measures across participants and repeating the prediction procedure. Care was taken to avoid shuffling between families or sites.

To compare models, we used the corrected resampled t-test (Nadeau & Benigo, 2003; Bouckaert & Frank, 2004) because a permutation test would not be valid. To control for multiple comparisons, we performed a false discovery rate (FDR) correction with q < 0.05.

### 2.8. Data and code availability

The lists of participants, features, and behavior scores utilized are released for both datasets. Data for the HCP are available in this Github repository (https://github.com/ThomasYeoLab/Ooi2022_MMP_HCP). Data for the ABCD are available on the NIMH Data Archive (NDA) website^1^ (https://dx.doi.org/10.15154/1523482). The folder structure for ABCD is similar to that of the HCP. Any additional data can be accessed directly from the HCP (https://www.humanconnectome.org/) and ABCD (https://abcdstudy.org/) websites, as they are both publicly available.

Code for this study is publicly available in the Github repository maintained by the Computational Brain Imaging Group (https://github.com/ThomasYeoLab/CBIG). Code specific to the regression models and analyses in this study can be found here (https://github.com/ThomasYeoLab/Standalone_Ooi2022_MMP).

a. To replicate the results in this study, first download the features and training-test splits provided for each dataset, and train the regression algorithms with the regression code from the CBIG repository.
b. To compare a new set of features against the benchmarks in this study. Download the participant list and training-test split for each dataset. Using the participant list provided in each dataset repository, extract a #features x #participants matrix for each participant in the list and perform the predictions using the regression codes from the CBIG repository using the same training-test splits.
c. To compare a new predictive model against the benchmarks in this study, download the features and training-test splits for each dataset. Using the same features and training-test splits, predictive performance of the new model can be compared to the results in this study.

Processing pipelines for diffusion data (https://github.com/ThomasYeoLab/CBIG/tree/master/stable_projects/preprocessing/CBIG2022_DiffProc), and functional data (https://github.com/ThomasYeoLab/CBIG/tree/master/stable_projects/preprocessing/CBIG_fMRI_Preproc2016) are provided in their respective links.

## 3. Results

### 3.1. Functional connectivity (FC) outperforms other features for predicting behavior

Across both HCP and ABCD, for each feature-type, a separate KRR model was trained to predict each of three behavioral components and each behavioral measure. Figure 1 shows the KRR predictive performance (Pearson’s correlation) averaged across single-feature-type predictive models for each of the three behavioral components. Figure 1 also shows the predictive performance averaged across single-feature-type predictive models for all behavioral measures, which we refer to as “grand average”. The grand average corresponded to averaging the prediction performance across 58 behavioral measures in the case of HCP and 36 behavioral measures in the case of ABCD.

**Figure 1.**
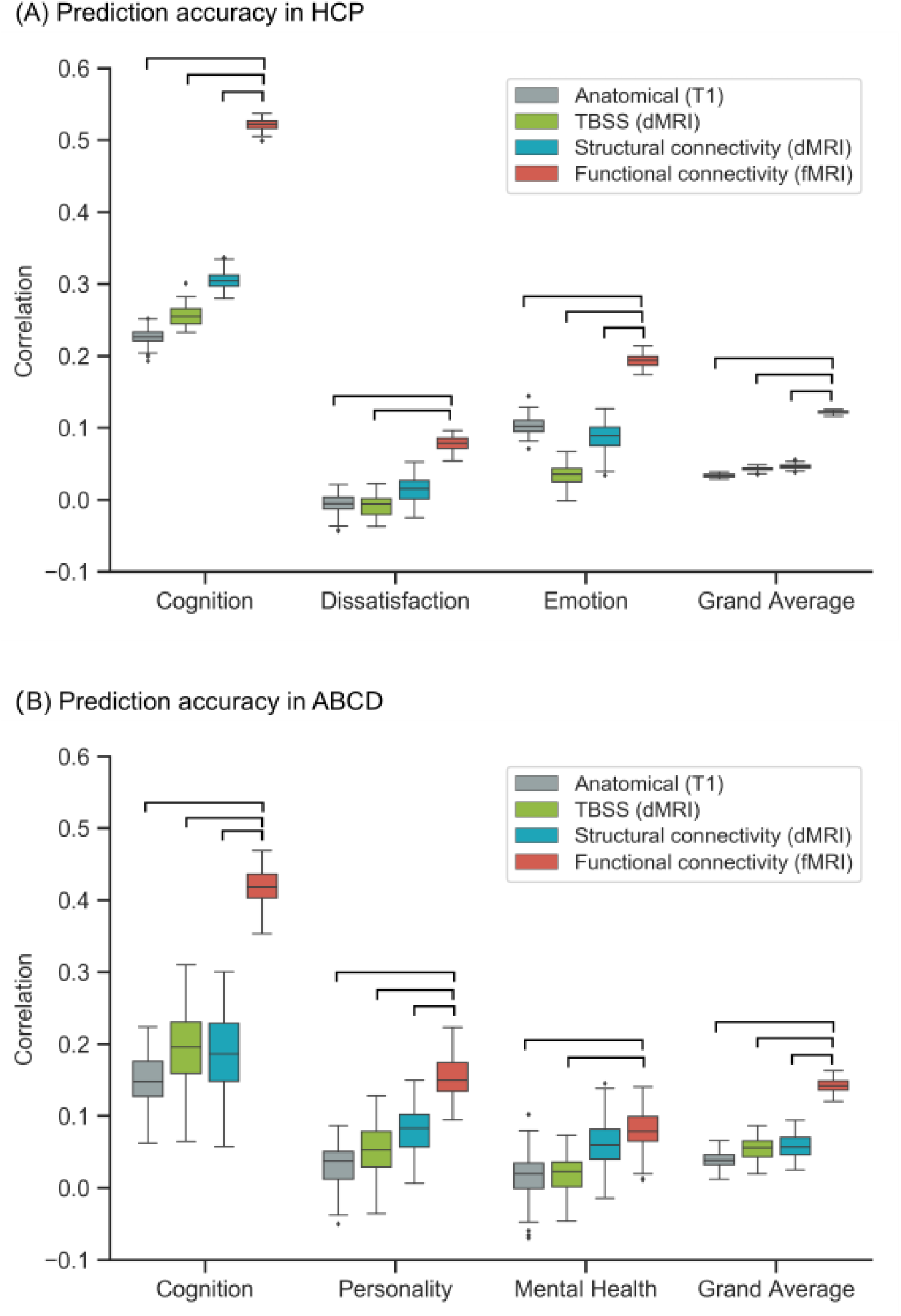
Functional connectivity (FC) outperforms other modalities for kernel ridge regression (KRR). (A) Prediction performance (Pearson’s correlation) of KRR averaged across single-feature-type predictive models within each modality (anatomical, TBSS, structural connectivity, functional connectivity) in the HCP dataset. Results are shown for the three behavioral components and “grand average” obtained by averaging prediction performance across 58 behavioral measures. Each boxplot shows the distribution of performance over 60 repetitions of the nested cross-validation procedure. (B) Prediction performance (Pearson’s correlation) of KRR averaged across single-feature-type predictive models within each modality (anatomical, TBSS, structural connectivity, functional connectivity) in the ABCD dataset. Results are shown for the three behavioral components and “grand average” obtained by averaging prediction performance across 36 behavioral measures. Each boxplot shows the distribution of performance over 120 repetitions of the nested cross-validation procedure. Connecting lines between boxes denote significantly different model performances after correction for multiple comparisons (FDR q < 0.05).

In both datasets, FC-based models performed the best, especially in the case of the cognition component (p < 1e-9) and the grand average (p < 1e-17). Predictions of cognition were also significantly better for FC-based models compared to models trained on anatomical features, TBSS, and SC in both the HCP (p=1.5e-20, p=2.4e-17, p=4.2e-9 respectively) and ABCD (p=1.1e-28, p=2.6e-15, p=1.4e-13 respectively) datasets.

Similar results were obtained with COD (Figure S1). Prediction performance (Pearson’s correlation) for each individual behavioral measure can be found in Figures S2 to S6. LRR and elastic net yielded slightly lower prediction performance, but similar conclusions (Figures 2 and 3).

**Figure 2.**
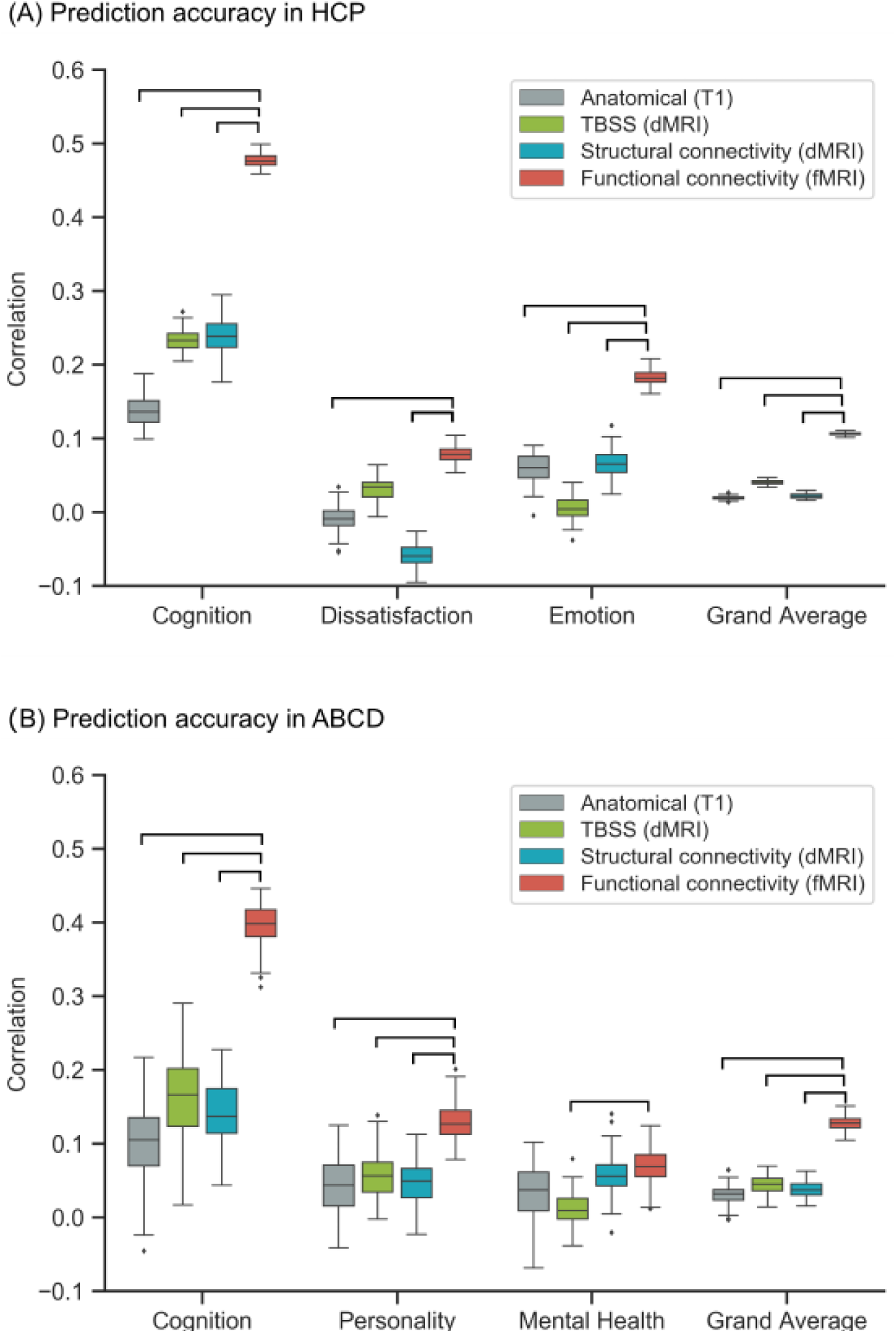
Functional connectivity (FC) outperforms other modalities for linear ridge regression (LRR). Figure is the same as Figure 1 except that LRR was utilized instead of kernel ridge regression. (A) Prediction performance (Pearson’s correlation) of LRR averaged across single-feature-type predictive models within each modality (anatomical, TBSS, structural connectivity, functional connectivity) in the HCP dataset. Results are shown for the three behavioral components and “grand average” obtained by averaging prediction performance across 58 behavioral measures. Each boxplot shows the distribution of performance over 60 repetitions of the nested cross-validation procedure. (B) Prediction performance (Pearson’s correlation) of LRR averaged across single-feature-type predictive models within each modality (anatomical, TBSS, structural connectivity, functional connectivity) in the ABCD dataset. Results are shown for the three behavioral components and “grand average” obtained by averaging prediction performance across 36 behavioral measures. Each boxplot shows the distribution of performance over 120 repetitions of the nested cross-validation procedure. Connecting lines between boxes denote significantly different model performances after correction for multiple comparisons (FDR q < 0.05).

**Figure 3.**
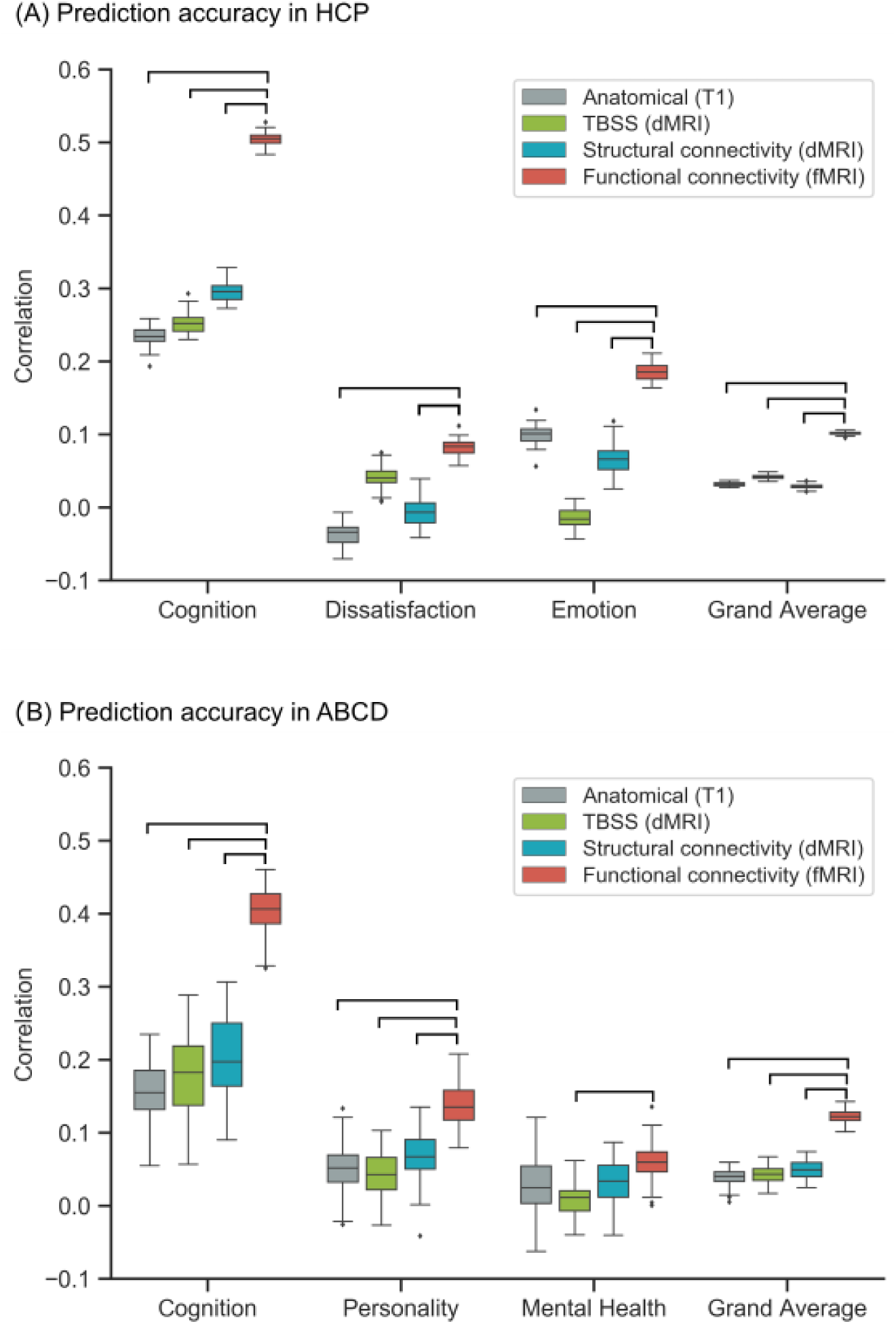
Functional connectivity (FC) outperforms other modalities for elastic net. Figure is the same as Figure 1 except that elastic net was utilized instead of kernel ridge regression. (A) Prediction performance (Pearson’s correlation) of elastic net averaged across single-feature-type predictive models within each modality (anatomical, TBSS, structural connectivity, functional connectivity) in the HCP dataset. Results are shown for the three behavioral components and “grand average” obtained by averaging prediction performance across 58 behavioral measures. Each boxplot shows the distribution of performance over 60 repetitions of the nested cross-validation procedure. (B) Prediction performance (Pearson’s correlation) of elastic net averaged across single-feature-type predictive models within each modality (anatomical, TBSS, structural connectivity, functional connectivity) in the ABCD dataset. Results are shown for the three behavioral components and “grand average” obtained by averaging prediction performance across 36 behavioral measures. Each boxplot shows the distribution of performance over 120 repetitions of the nested cross-validation procedure. Connecting lines between boxes denote significantly different model performances after correction for multiple comparisons (FDR q < 0.05).

Figure 4 shows the best single-feature-type (based on KRR) from each modality for each behavior component and grand average. In both datasets, FC was better than anatomical features, TBSS and SC. Similar results were obtained with LRR and elastic net (Figures S7 to S8).

**Fig 4.**
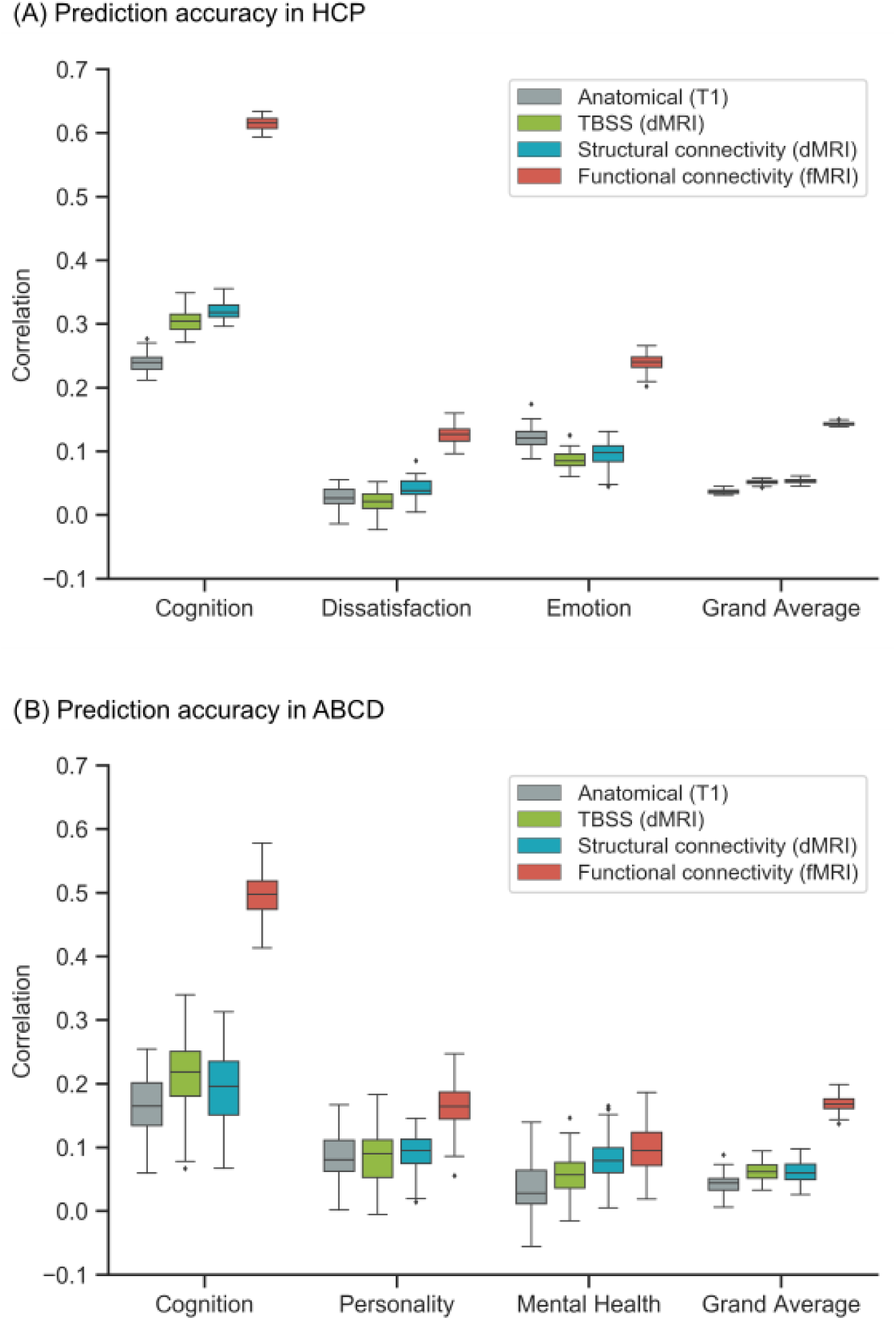
Functional connectivity (FC) outperforms other modalities for kernel ridge regression (KRR). Figure is the same as Figure 1 except that the best-feature-type for each behavioral measure was selected instead of averaging across feature-types. (A) Prediction performance (Pearson’s correlation) of KRR for the best performing feature-type within each modality in the HCP dataset. For the cognition component, the best features were cortical volume, TBSS AD, SC stream length and language FC. For the dissatisfaction component, the best features were cortical thickness, TBSS OD, SC stream count and working memory FC. For the emotion component, the best features were cortical volume, TBSS OD, SC stream length and social cognition FC. For the grand average, the best features were cortical volume, TBSS AD, SC stream count and language FC. (B) Prediction performance (Pearson’s correlation) of KRR for the best performing feature-type within each modality in the ABCD dataset. For the cognition component, the best features were cortical thickness, TBSS ICVF, SC FA and N-back FC. For the personality component, the best features were cortical volume, TBSS AD, SC stream length and N-back FC. For the mental health component, the best features were cortical thickness, TBSS ISOVF, SC RD and SST FC. For the grand average, the best features were cortical thickness, TBSS OD, SC RD and N-back FC. We note that no statistical test was performed here since maximum statistic is prone to outliers.

### 3.2. All modalities predict cognition better than chance

Figures 5 and 6 show the KRR prediction performance (Pearson’s correlation) for each single-feature-type predictive model in the HCP and ABCD datasets respectively. Across all feature types and both datasets, the cognitive component was predicted better than chance. This was not the case for the other two behavioral components in HCP and ABCD. Similar results were obtained with COD (Figures S9 and S10), as well as LRR and elastic net (Figures S11 to S14).

**Figure 5.**
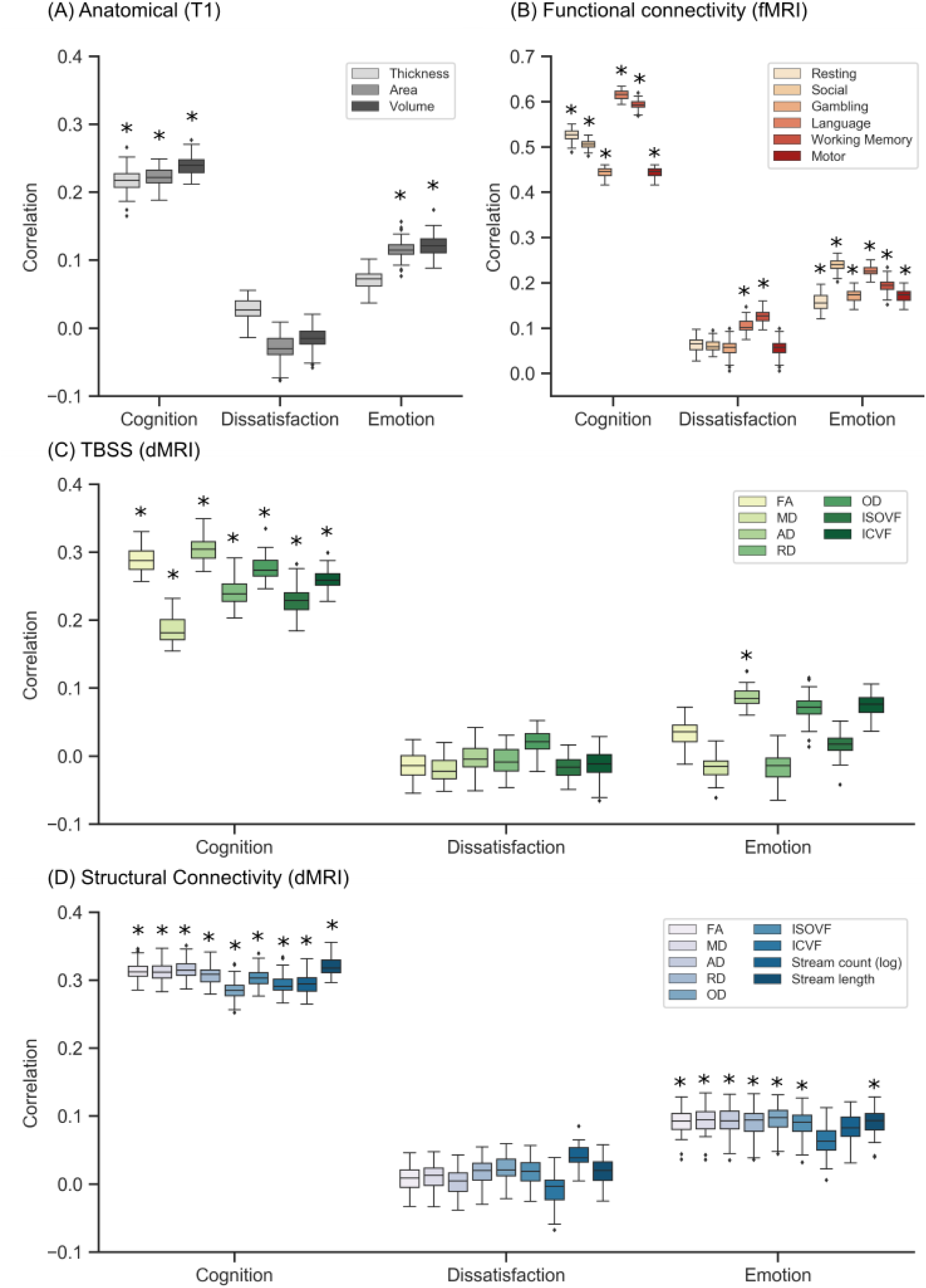
Prediction performance (Pearson’s correlation) of kernel ridge regression (KRR) for each single-feature-type in the HCP dataset. Results are shown separately for (A) anatomical features, (B) FC, (C) TBSS and (D) structural connectivity. * denotes that the model performed better than chance after correction for multiple comparisons (FDR q < 0.05). Across all feature types, the cognitive component was predicted better than chance. This was not the case for the other two behavioral components. We note that no statistical test was performed to compare models; see Figures 1 to 4 for model comparisons.

**Figure 6.**
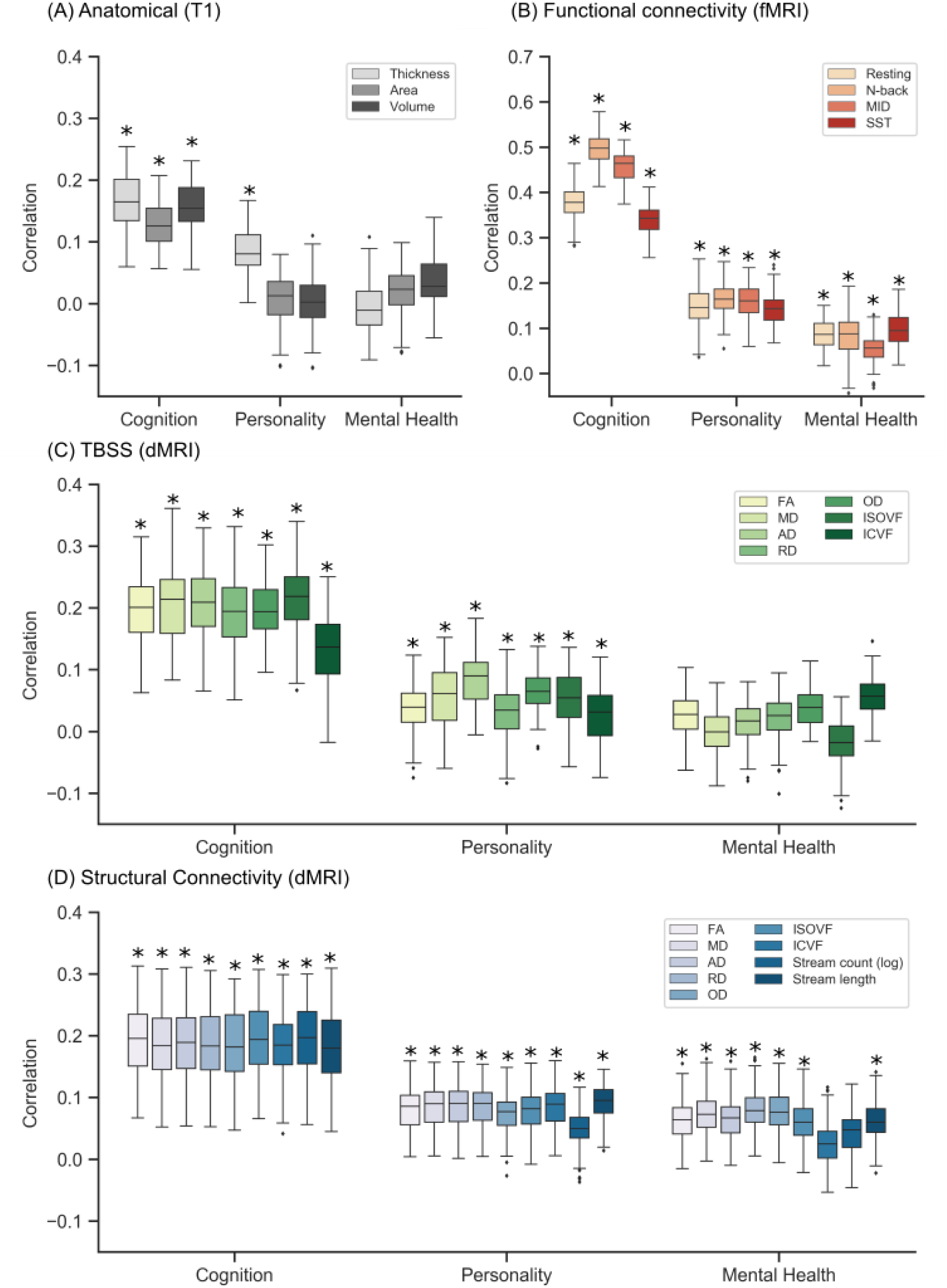
Prediction performance (Pearson’s correlation) of kernel ridge regression (KRR) for each single-feature-type in the ABCD dataset. Figure is the same as Figure 5 except that the results here corresponded to the ABCD dataset (instead of HCP). Results are shown separately for (A) anatomical features, (B) FC, (C) TBSS and (D) structural connectivity. * denotes that the model performed better than chance after correction for multiple comparisons (FDR q < 0.05). Across all feature types, the cognitive component was predicted better than chance. This was not the case for the other two behavioral components. We note that no statistical test was performed to compare models; see Figures 1 to 4 for model comparisons.

Overall, this suggests that in the case of healthy children and young adults, brain characteristics captured by MRI most strongly reflect individual differences in cognition and might reflect the difficulty in capturing subjective aspects of behavior through imaging.

### 3.3. Combining resting and task FC was as good as combining across all modalities

Figure 7 shows the prediction performance (Pearson’s correlation) from combining various MRI features based on stacking or multi-KRR. For comparison, the best single-feature-type from KRR is shown. We note that the best single-feature-type always corresponded to FC (Figure 4).

**Figure 7.**
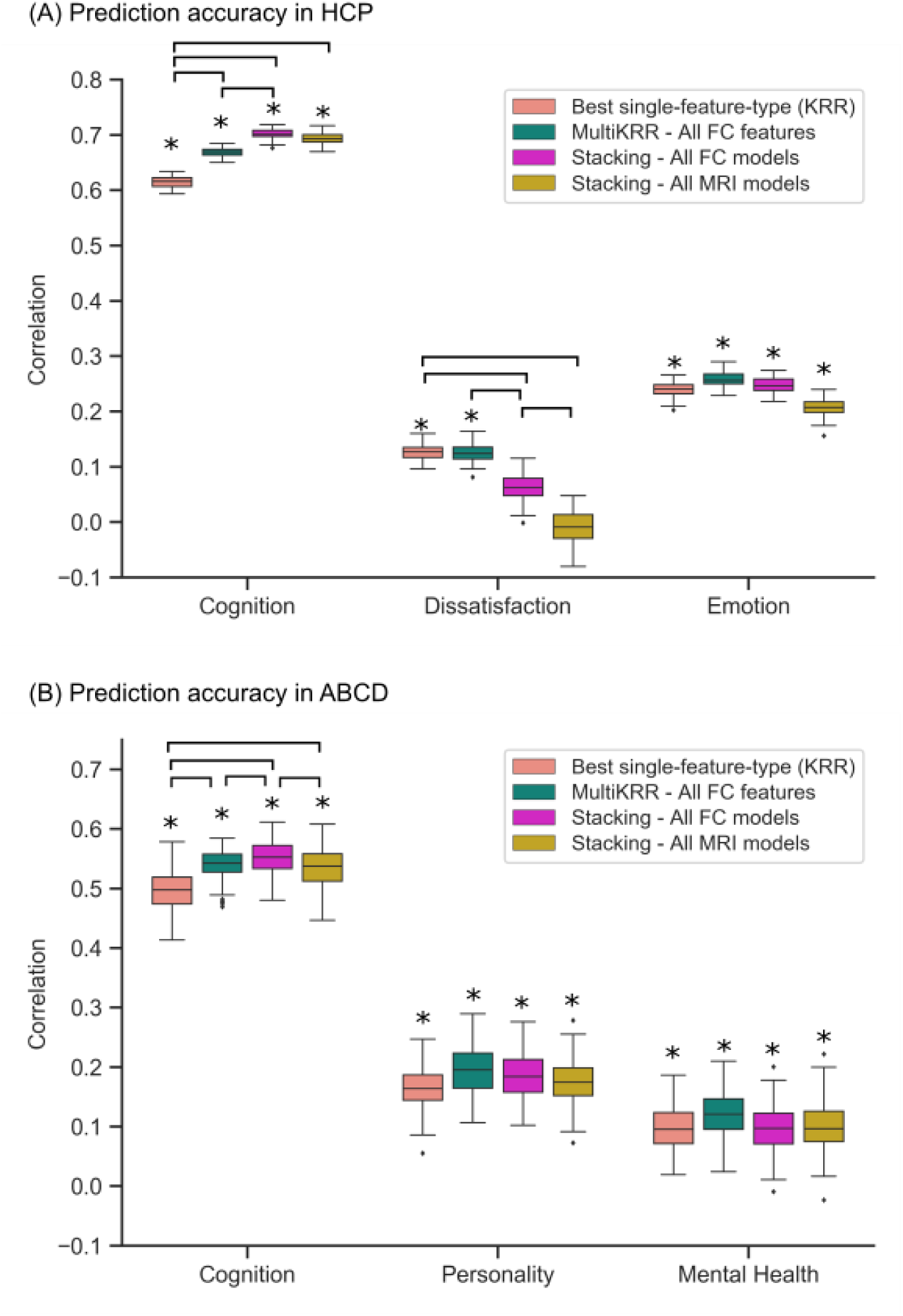
Combining resting and task FC was as good as combining across all modalities. (A) Prediction performance (Pearson’s correlation) from combining various MRI features and modalities in the HCP dataset. We considered multi-KRR of all FC features, stacking of all FC models and stacking of all single-feature-type models across all modalities. For comparison, the best single-feature-type from KRR is shown. Each boxplot shows the distribution over 60 repetitions of the nested cross-validation procedure. (B) Prediction performance (Pearson’s correlation) from combining various MRI features and modalities in the ABCD dataset. We considered multi-KRR of all FC features, stacking of all FC models and stacking of all single-feature-type models across all modalities. For comparison, the best single-feature-type from KRR is shown. Each boxplot shows the distribution over 120 repetitions of the nested cross-validation procedure. * denotes that the model performed better than chance after correction for multiple comparisons (FDR q < 0.05). Connecting lines between boxes denote significantly different model performances after correction for multiple comparisons (FDR q < 0.05). Combining features led to improvements in prediction of the cognition component. Combining all modalities was not better than simply combining resting and task FC.

In the case of the cognitive component, multi-KRR of all FC features, stacking of all FC-based models and stacking of all single-feature-type models of all modalities yielded better prediction performance than the best single-feature-type model in both the HCP (p=7.7e-4, p=1.3e-7, p=8.7e-6 respectively) and ABCD (p=1.1e-4, p=5.5e-8, p=0.0057 respectively).

Furthermore, stacking all modalities did not provide any significant improvement over stacking FC-based models. In addition, stacking the best single-feature-type models from each modality was not better than stacking FC-based models (Figure S15). Similar results were obtained with COD (Figure S16). Overall, this suggests that the gain from stacking all modalities was largely due to the variance account for in FC.

Finally, combining multiple features did not improve the prediction of the remaining two behavioral components in both datasets. In fact, in the case of life dissatisfaction in the HCP dataset, the best performing single-feature model was statistically better than stacking all FC-based models or stacking all single-feature-type models of all modalities.

## 4. Discussion

In this study, we demonstrated that functional connectivity features led to better predictive performance than features derived from anatomical or diffusion MRI. This finding was replicated across two datasets, and three regression models. We also found that integrating features across modalities through stacking mainly improved predictions for cognition, but not for other behaviors. Finally, we showed that combining all features from all modalities was not better than combining functional connectivity features.

### 4.1. Behavioral prediction using FC versus other modalities

There are relatively few studies comparing FC prediction with other modalities in young healthy participants, all of whom focused on the young adult HCP dataset. Dhamala showed that resting FC outperformed SC in predicting cognitive performance in the young adult HCP dataset (Dhamala et al., 2021). Similarly, Mansour also showed that high resolution resting FC achieved higher accuracy than high resolution SC and anatomical features for cognition in the young adult HCP dataset (Mansour et al., 2021). In this study, we replicated Dhamala and Mansour’s results not just in the young adult HCP dataset, but also in the lesser utilized young children (ABCD) dataset. Similar to Dhamala and Mansour, we observed that FC outperformed the other modalities when predicting cognition using KRR. In the HCP, FC achieved a correlation coefficient of between 0.44 – 0.62 using KRR when predicting cognition, whereas diffusion features were between 0.19 – 0.32, and anatomical features were between 0.22 – 0.24. In the ABCD dataset, we found a similar trend that FC outperformed other modalities in prediction of cognition. Moreover, we extended Dhamala and Mansour’s work in two other ways. First, Dhamala and Mansour only considered stream counts in the SC matrix. Here, we considered additional diffusion features from DTI and NODDI models averaged across tracts connecting each pair of brain regions. We also considered DTI and NODDI features extracted from the TBSS skeleton (Smith et al., 2006), which is a widely used approach. Second, we also considered task FC in addition to resting FC. Second, we additionally show that the better behaviour prediction extends over other behavioural domains, and the “grand average” across behavioral measures.

However, we note discrepancy with Rasero and colleagues (Rasero et al., 2021), who showed that the “local connectome” derived from diffusion features was able to outperform resting FC in the HCP when predicting cognition. Rasero found that the local connectome was able to predict “global cognition” with a COD of 0.049, while FC could only achieve an accuracy of 0.016. Conversely, in our study, we found that resting FC could achieve a COD of 0.25 in the cognition component, and diffusion features from SC ranged between a COD of 0.042-0.074. Therefore, our diffusion prediction performance was comparable to Rasero, but our FC prediction performance was significantly better.

One possible reason for this discrepancy might be due to differences between Rasero’s tabulation of “global cognition” and our cognition component, although this cannot explain that our diffusion prediction performance is similar. Another possible reason might be related to fMRI preprocessing, e.g., Rasero opted not to perform GSR on fMRI data, which might have improved prediction performance (Li et al., 2019). Other reasons might be parcellation choice or the application of PCA on the FC features prior to prediction.

Our results are also inconsistent with Xiao and colleagues (Xiao et al., 2021), who found that anatomical features were able to outperform both functional and diffusion features in predicting visual working memory. One potential discrepancy is the use of the CAM-CAN dataset, which focused on elderly participants. It is possible, that our results hold for young healthy participants, but not older participants, perhaps due to late-life age related changes in brain anatomy.

### 4.2. Prediction of cognition is better than other behavioural domains

Previous studies from our group have shown that it’s easier to predict cognition than other measures when using FC (Kong et al., 2019; Li et al., 2019; Liégeois et al., 2019; Chen et al., 2020; Kong et al., 2021). Mansour and colleagues extended this result by showing that this is also true for anatomical and diffusion MRI in the HCP. Our current study confirmed Mansour’s results and replicated them in a new independent ABCD dataset.

Attaining better prediction for cognitive behaviour might be due to the subjective nature of personality and emotion, which might result in additional difficulty in predicting them. For example, Uher has described a lack of explicit formulation when investigating personality traits (Uher, 2015). This could result in greater difficulty in predicting such scores with a more subjective nature using brain imaging (Dubois et al., 2018). As such, we might expect to see increases in prediction performances of personality and emotion as reliability of behavioural measures increase.

### 4.3. Multimodal integration

Recent studies have suggested that task FC achieves better prediction of cognition over resting FC (Rosenberg et al., 2016; Greene et al., 2018; Jiang et al., 2020). Furthermore, combining task and resting FC in the young adult HCP dataset further boosts the prediction of cognition (Elliott et al., 2019; Gao et al., 2019). Chen and colleagues further expanded on this by showing that combining resting and task FC improved prediction of cognition in ABCD with little or no improvement for other behavioral domains (Chen et al., 2020).

We replicated these previous results in the HCP and ABCD datasets. We also extended these results further by showing that combining resting and task FC was as good as combining all features from all modalities. A possible explanation for the lack of improvement when combining all modalities is that interindividual differences in functional and structural brain characteristics lead to similar behavior changes. This has been observed by Llera and colleagues, who showed that modes of variation linking behaviour to structural variation and behaviour to functional variation significantly overlap (Llera et al., 2019). Together with the superior performance of FC in predicting behavior, this suggests that diffusion and anatomical features might not contain behaviour-related variation outside what FC already contains.

We note that our results were consistent with Dhamala and colleagues (Dhamala et al., 2021), who found that combining resting FC and SC did not increase prediction accuracy of cognition. In our study, stacking models that combined anatomical, diffusion and FC features were not significantly better than stacking models that combined only task and resting FC features. We expanded Dhamala’s work by showing the consistency of this finding in the ABCD dataset and by considering a wider range of features. More specifically, we show that neither the inclusion of anatomical data, nor the wider range of diffusion features could boost prediction performance above what could be achieved from integrating the various FC features.

However, we note that our results were inconsistent with that of Rasero and colleagues (Rasero et al., 2021), who found improvements in prediction of global cognition when stacking anatomical, diffusion and FC features. The discrepancy could be due to the much better prediction performance of FC in our current study, compared with Rasero and colleagues. Given that prediction performance of FC features was much better than diffusion and anatomical features in our current study, there might be limited gain in combining functional with anatomical or diffusion features.

### 4.3. Limitations, methodological considerations and future work

The feature dimensionalities varied greatly across modalities. For example, in the case of cortical thickness, there were 400 features per participant, corresponding to the 400-region Schaefer parcellation. Both the SC and FC matrices comprised 79,800 features (corresponding to the lower portion of the 400 x 400 matrix) per participant. Finally, in the case of TBSS, there were 133k and 109k for HCP and ABCD respectively. Despite the great variation in the number of features, we note that the cross-validation framework obviates the need to control for the number of features. The reason is that more features could lead to overly complex models and poor performance in the out-of-sample data. Indeed, FC outperformed TBSS despite having less features.

In this study, we have mainly focused on prediction using the 400 cortical ROIs from the Schaefer parcellation. In the case of anatomical features, we did not include contributions from subcortical regions to allow for fair comparisons among surface, thickness and volumetric features – there’s no concept of surface and thickness for subcortical structures. Given that we excluded subcortical structures for anatomical features, we also decided to exclude subcortical structures from the functional and structural connectivity analyses in order to be consistent.

Here, we have shown that FC outperforms diffusion and anatomical features in young healthy participants. However, several studies have begun to explore the value of multimodal individualized prediction performance in disease populations (Meng et al., 2017; Sui et al., 2020) and in aging (Engemann et al., 2020; Xiao et al., 2021), showing improved prediction of clinical markers with multimodal imaging (Mill et al., 2021). The benefits of multimodal imaging could be further explored in future work, focusing on the identification of disease and aging markers that can benefit from multimodal imaging, and comparing the utility of each modality in predicting these markers.

## 5. Conclusion

Through applying KRR, LRR, and elastic net regression to anatomical, diffusion and functional connectivity features in the HCP and ABCD datasets, we showed that functional connectivity was able to achieve better prediction of behavioral traits. Combining resting and task FC improved prediction of cognition, but not other behavioral traits. On the other hand, there was no additional benefit from combining all features from all modalities compared with combining resting and task FC, suggesting that FC features might encompass behaviorally relevant information from anatomical and diffusion features.

## Acknowledgements

Our research is currently supported by the Singapore National Research Foundation (NRF) Fellowship (Class of 2017), the NUS Yong Loo Lin School of Medicine (NUHSRO/2020/124/TMR/LOA), and the USA NIH (R01MH120080). Our computational work was partially performed on resources of the National Supercomputing Centre, Singapore (https://www.nscc.sg). Any opinions, findings and conclusions or recommendations expressed in this material are those of the authors and do not reflect the views of the Singapore NRF.

Data were provided [in part] by the Human Connectome Project, WU-Minn Consortium (Principal Investigators: David Van Essen and Kamil Ugurbil; 1U54MH091657) funded by the 16 NIH Institutes and Centers that support the NIH Blueprint for Neuroscience Research; and by the McDonnell Center for Systems Neuroscience at Washington University.

Data used in the preparation of this article were obtained from the Adolescent Brain Cognitive Development^SM^ (ABCD) Study (https://abcdstudy.org), held in the NIMH Data Archive (NDA). This is a multisite, longitudinal study designed to recruit more than 10,000 children age 9-10 and follow them over 10 years into early adulthood. The ABCD Study® is supported by the National Institutes of Health and additional federal partners under award numbers U01DA041048, U01DA050989, U01DA051016, U01DA041022, U01DA051018, U01DA051037, U01DA050987, U01DA041174, U01DA041106, U01DA041117, U01DA041028, U01DA041134, U01DA050988, U01DA051039, U01DA041156, U01DA041025, U01DA041120, U01DA051038, U01DA041148, U01DA041093, U01DA041089, U24DA041123, U24DA041147. A full list of supporters is available at https://abcdstudy.org/federal-partners.html. A listing of participating sites and a complete listing of the study investigators can be found at https://abcdstudy.org/consortium_members/. ABCD consortium investigators designed and implemented the study and/or provided data but did not necessarily participate in the analysis or writing of this report. This manuscript reflects the views of the authors and may not reflect the opinions or views of the NIH or ABCD consortium investigators. The ABCD data repository grows and changes over time. The ABCD data used in this report came from http://dx.doi.org/10.15154/1504041.

## Supplemental material

**Table S1.**
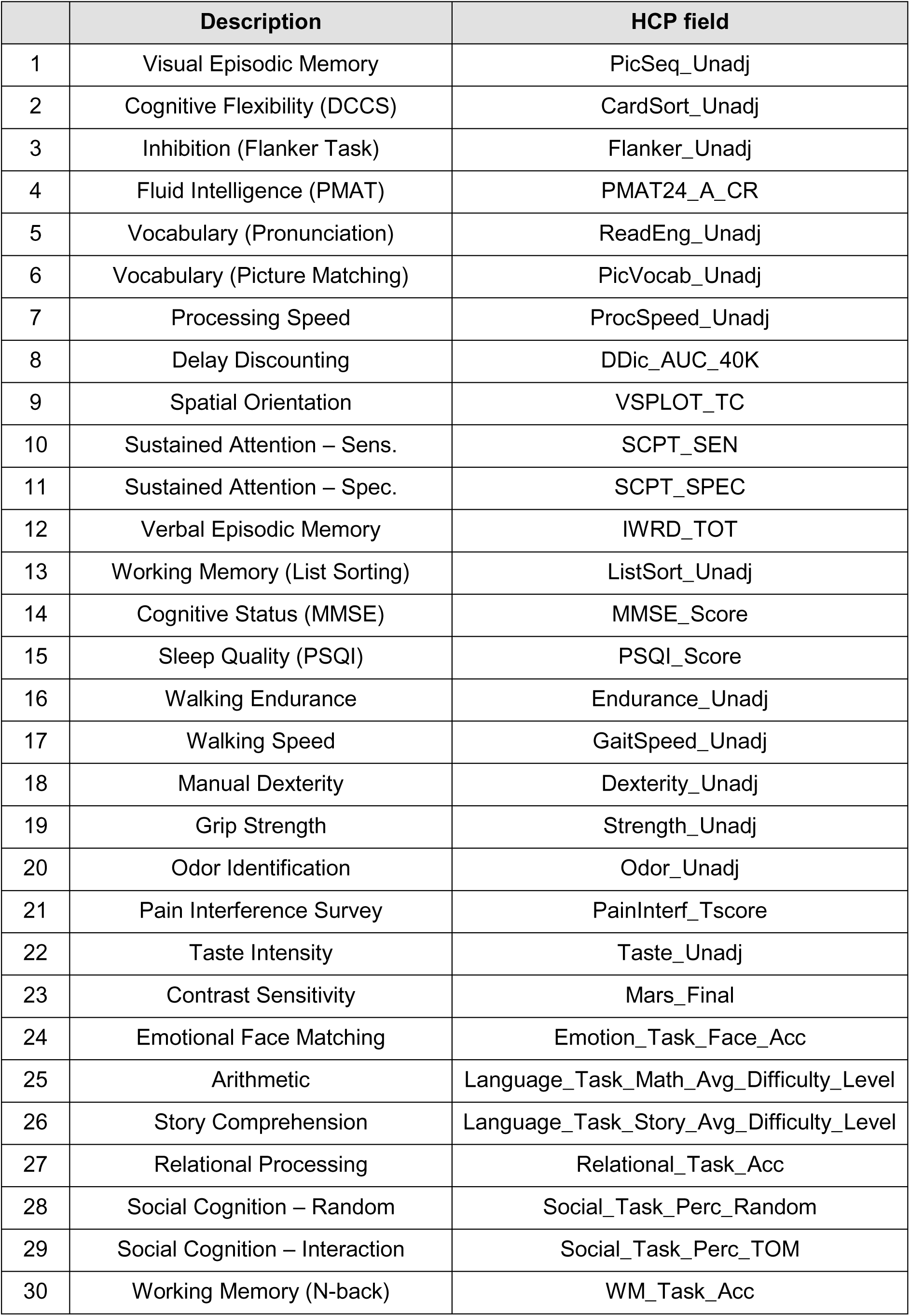
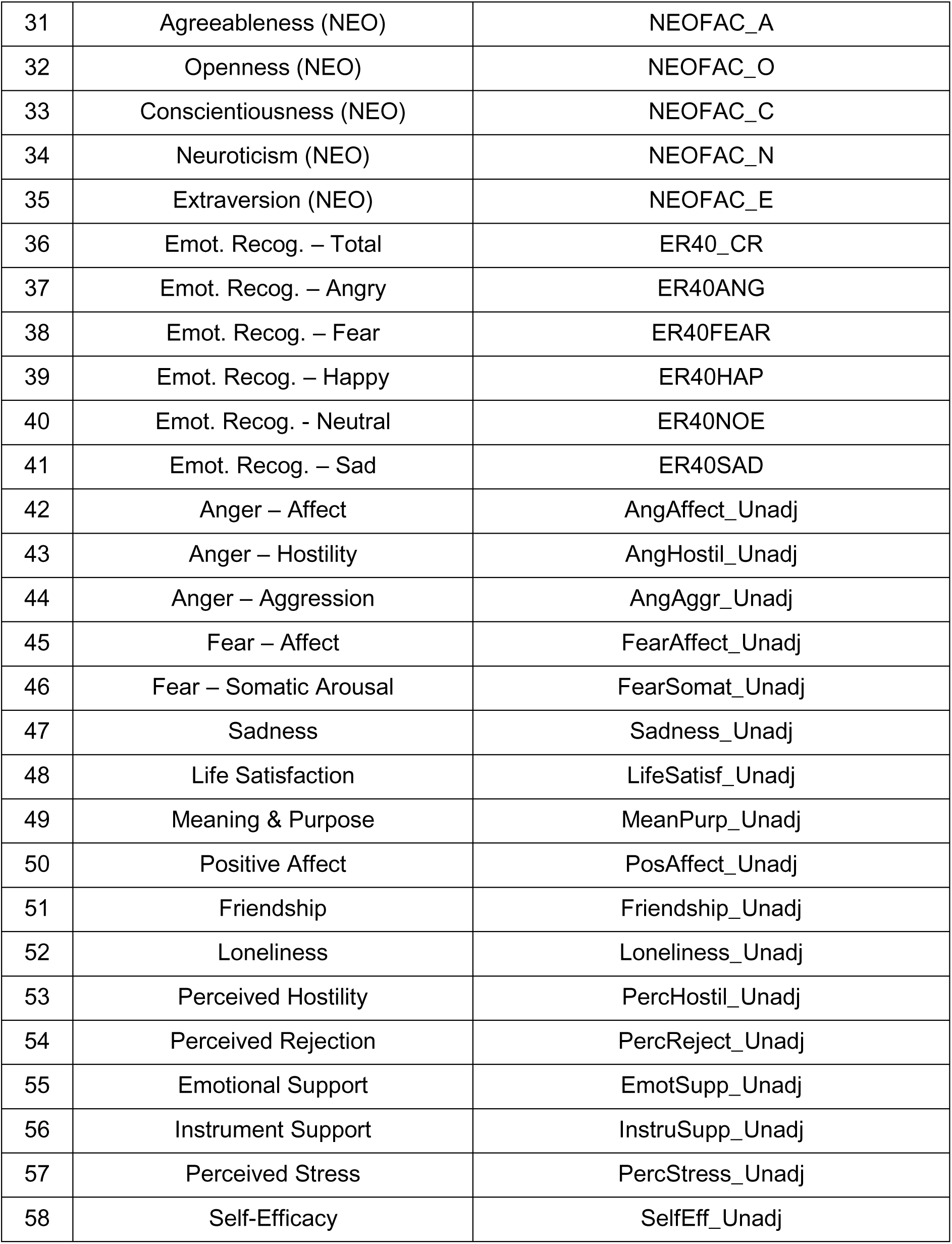
Behavioral measures for HCP

**Table S2.**
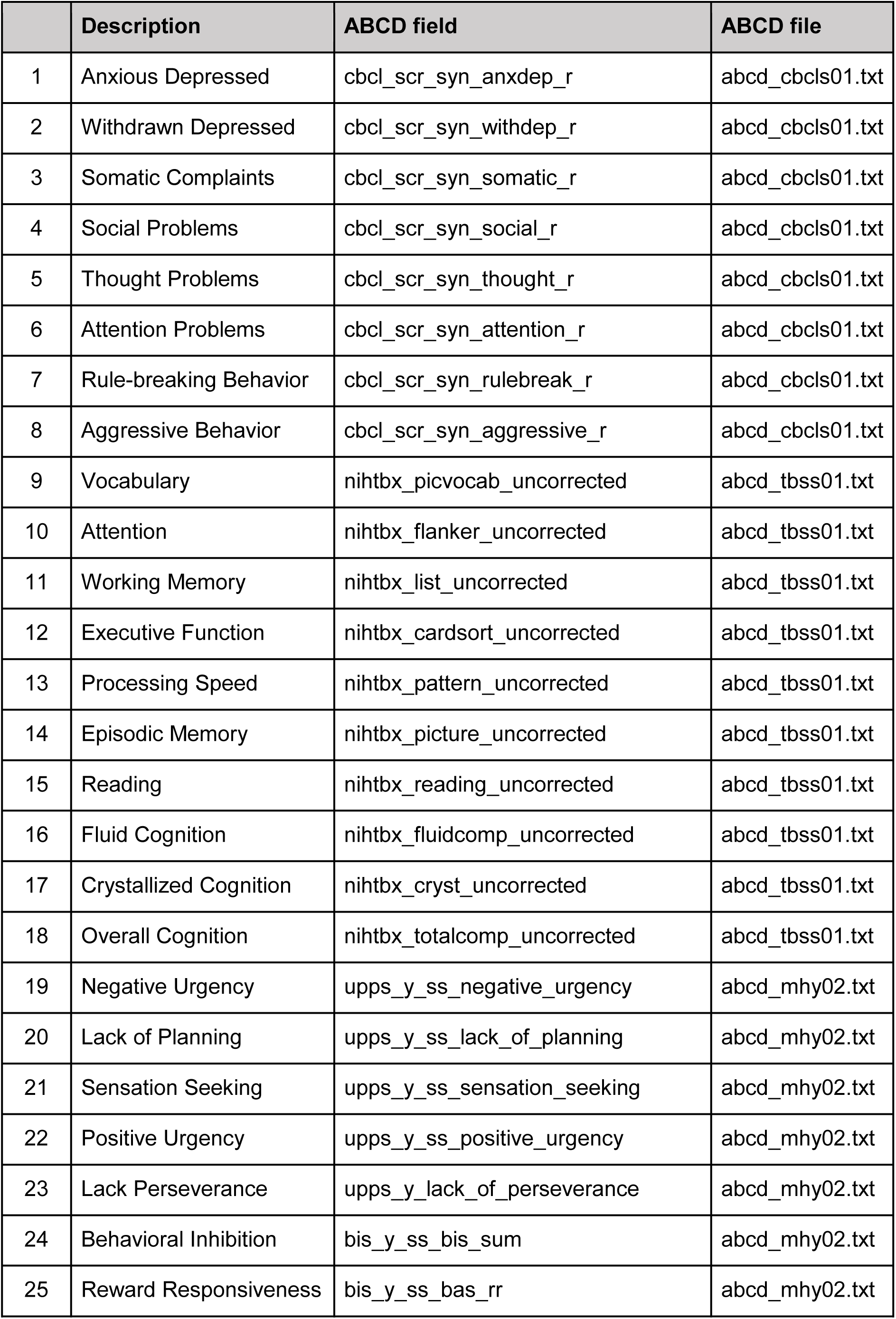
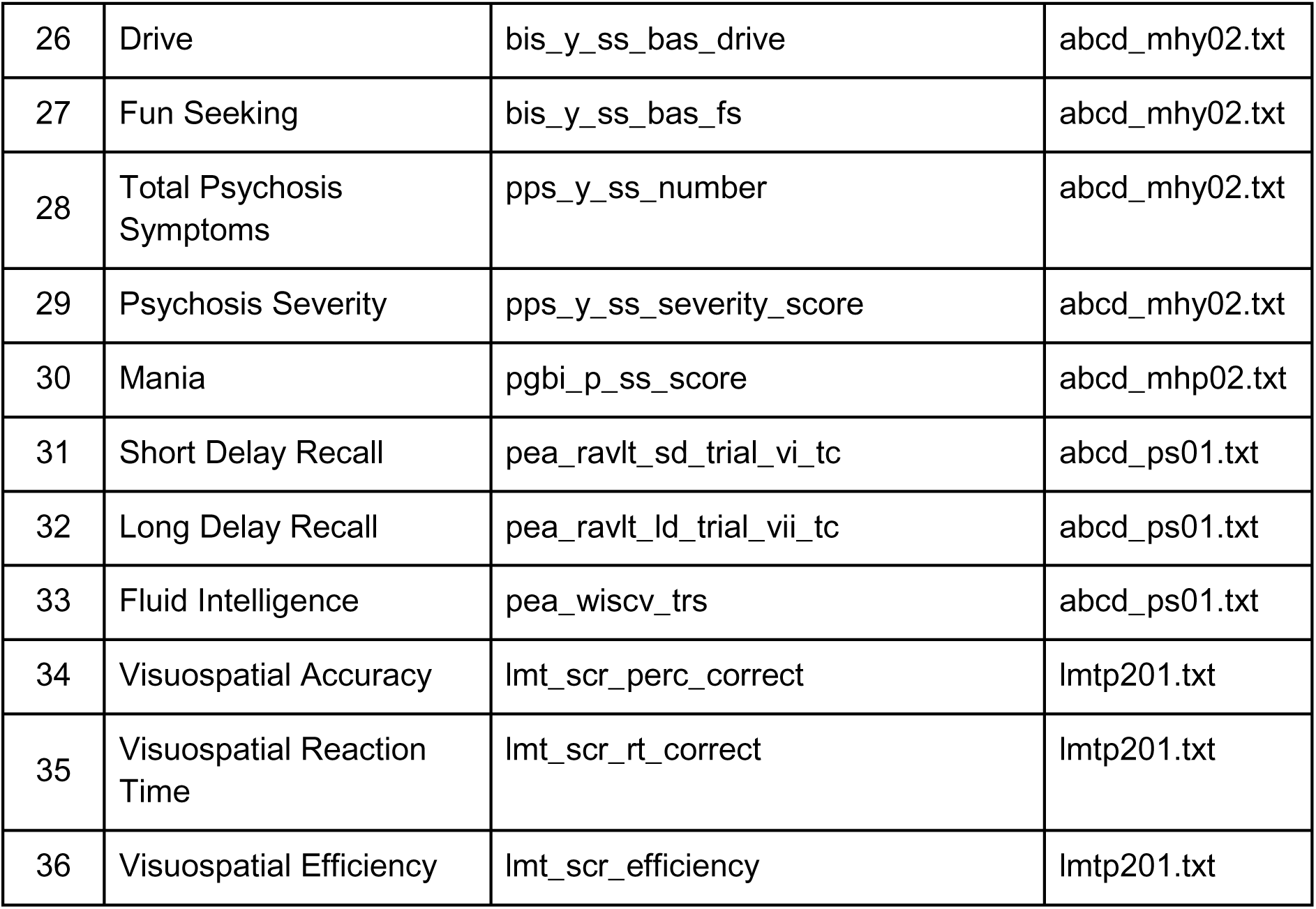
Behavioral measures for ABCD

**Table S3.**
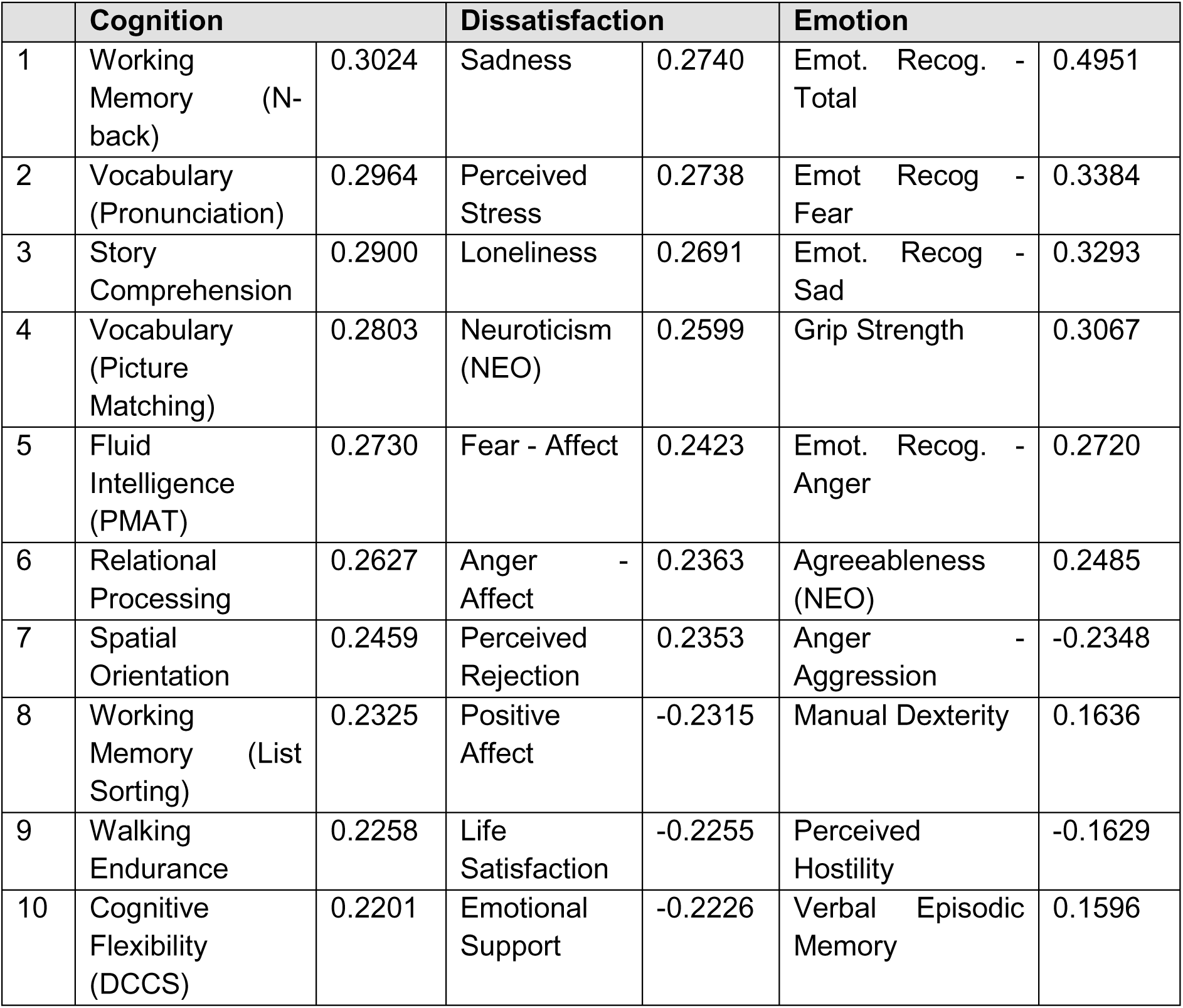
Top 10 loadings (absolute) for HCP behavioral factors

**Table S4.**
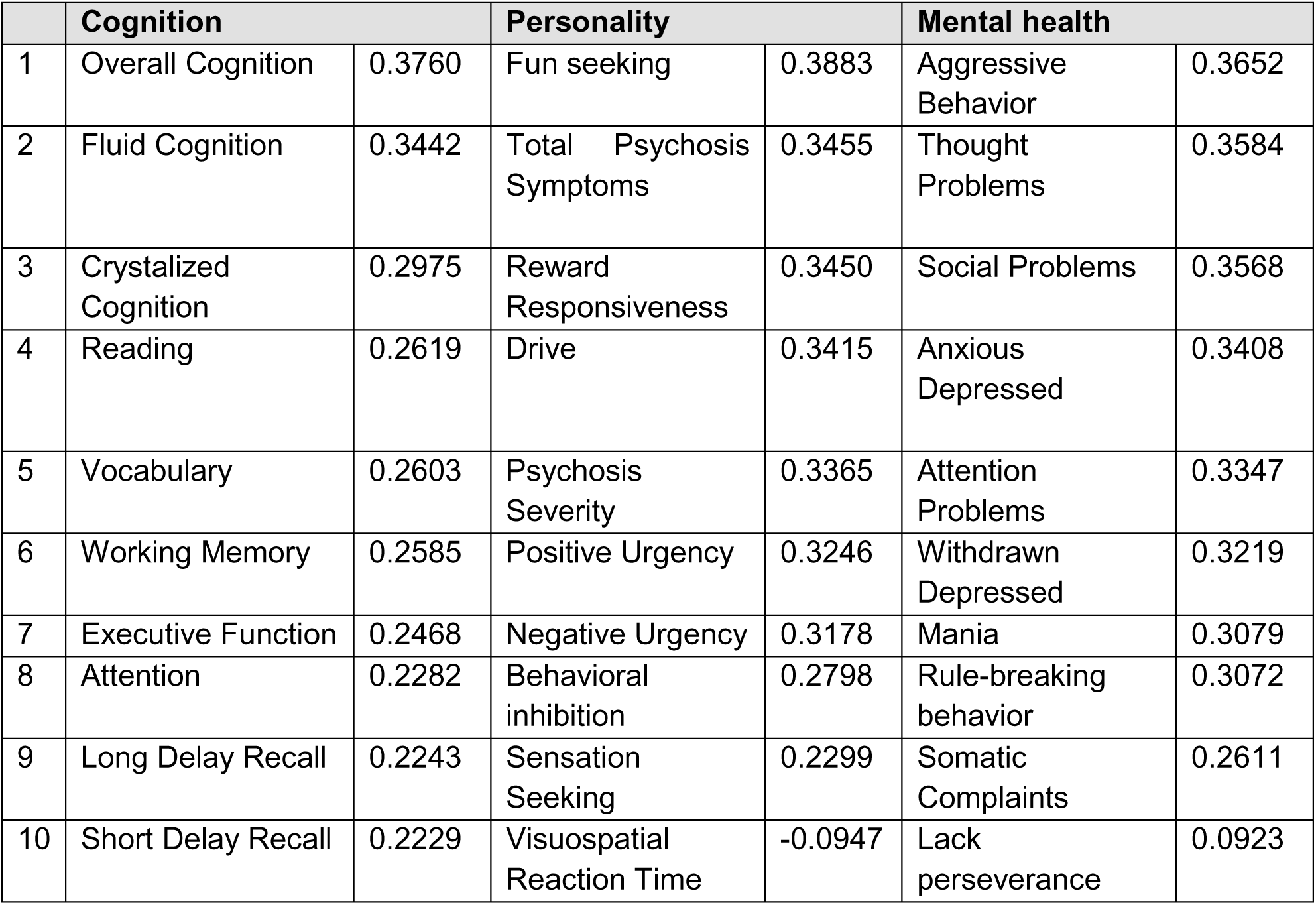
Top 10 loadings (absolute) for ABCD behavioral factors

**Table S5.**
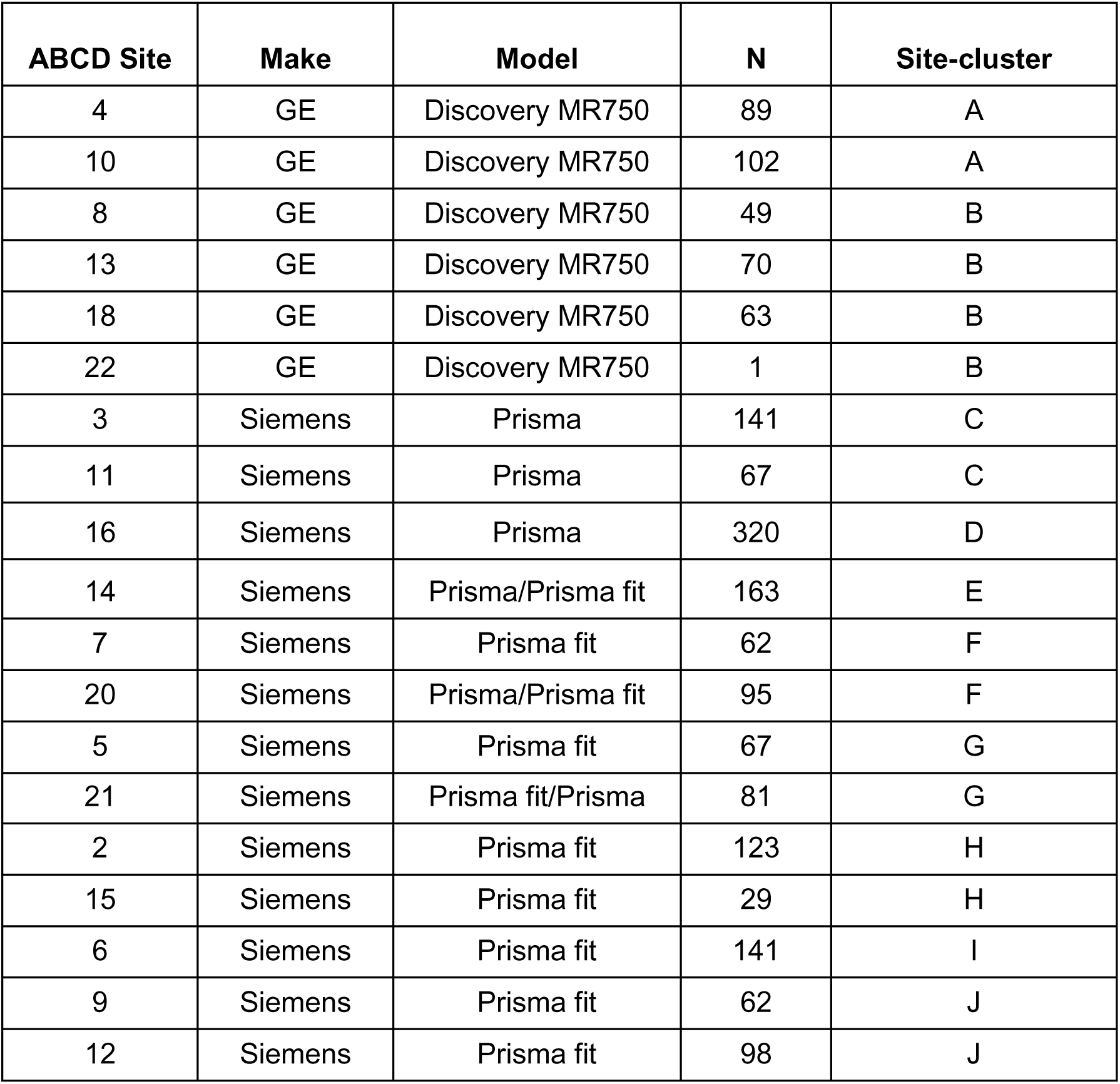
Site clusters for ABCD

**Figure S1.**
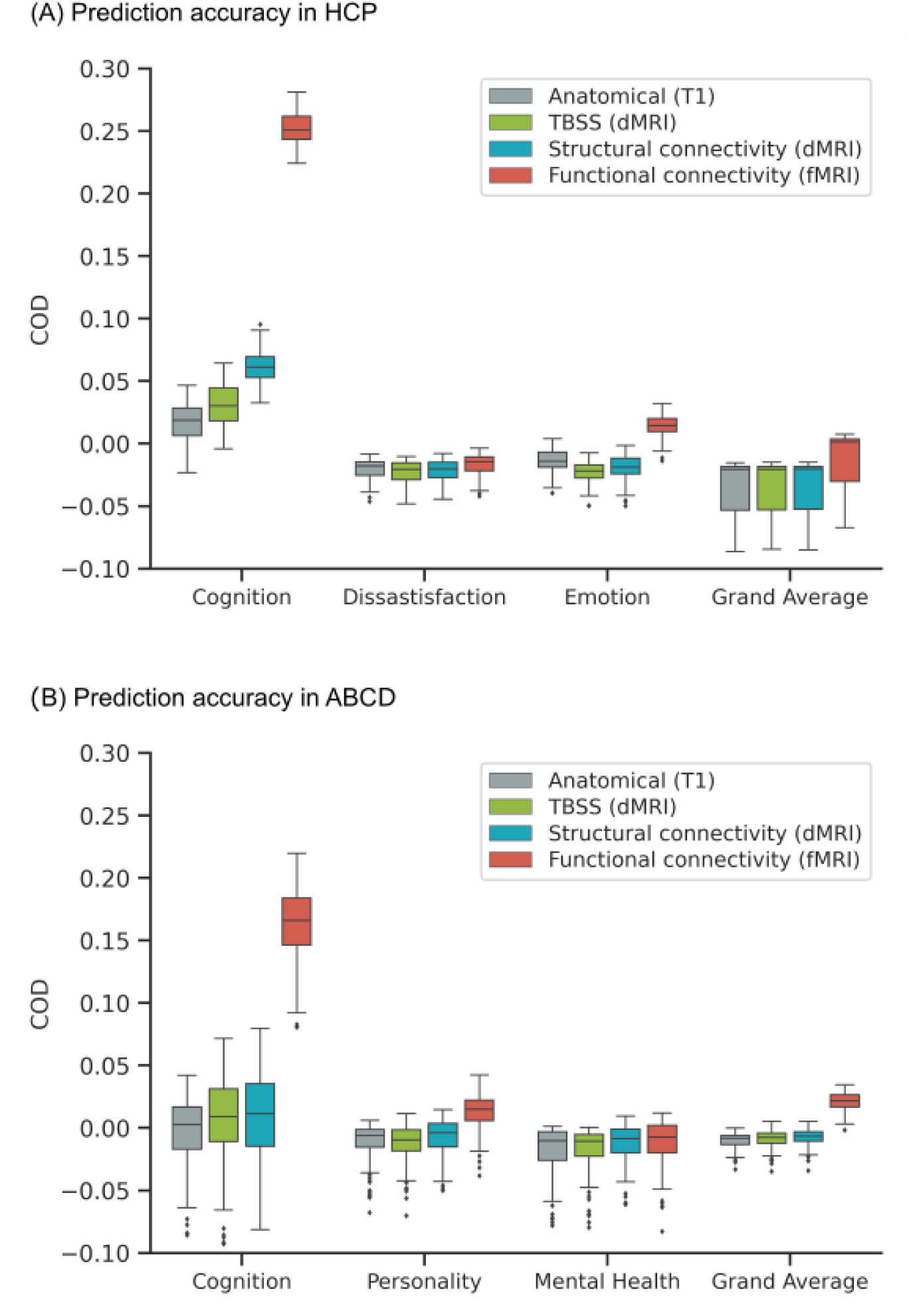
Functional connectivity (FC) outperforms other modalities for kernel ridge regression (KRR). Figure is the same as Figure 1 except that COD is shown instead of Pearson’s correlation. (A) Prediction performance (COD) of KRR averaged across single-feature-type predictive models within each modality (anatomical, TBSS, structural connectivity, functional connectivity) in the HCP dataset. Results are shown for the three behavioral components and “grand average” obtained by averaging prediction performance across 58 behavioral measures. Each boxplot shows the distribution of performance over 60 repetitions of the nested cross-validation procedure. (B) Prediction performance (COD) of KRR averaged across single-feature-type predictive models within each modality (anatomical, TBSS, structural connectivity, functional connectivity) in the ABCD dataset. Results are shown for the three behavioral components and “grand average” obtained by averaging prediction performance across 36 behavioral measures. Each boxplot shows the distribution of performance over 120 repetitions of the nested cross-validation procedure.

**Figure S2.**
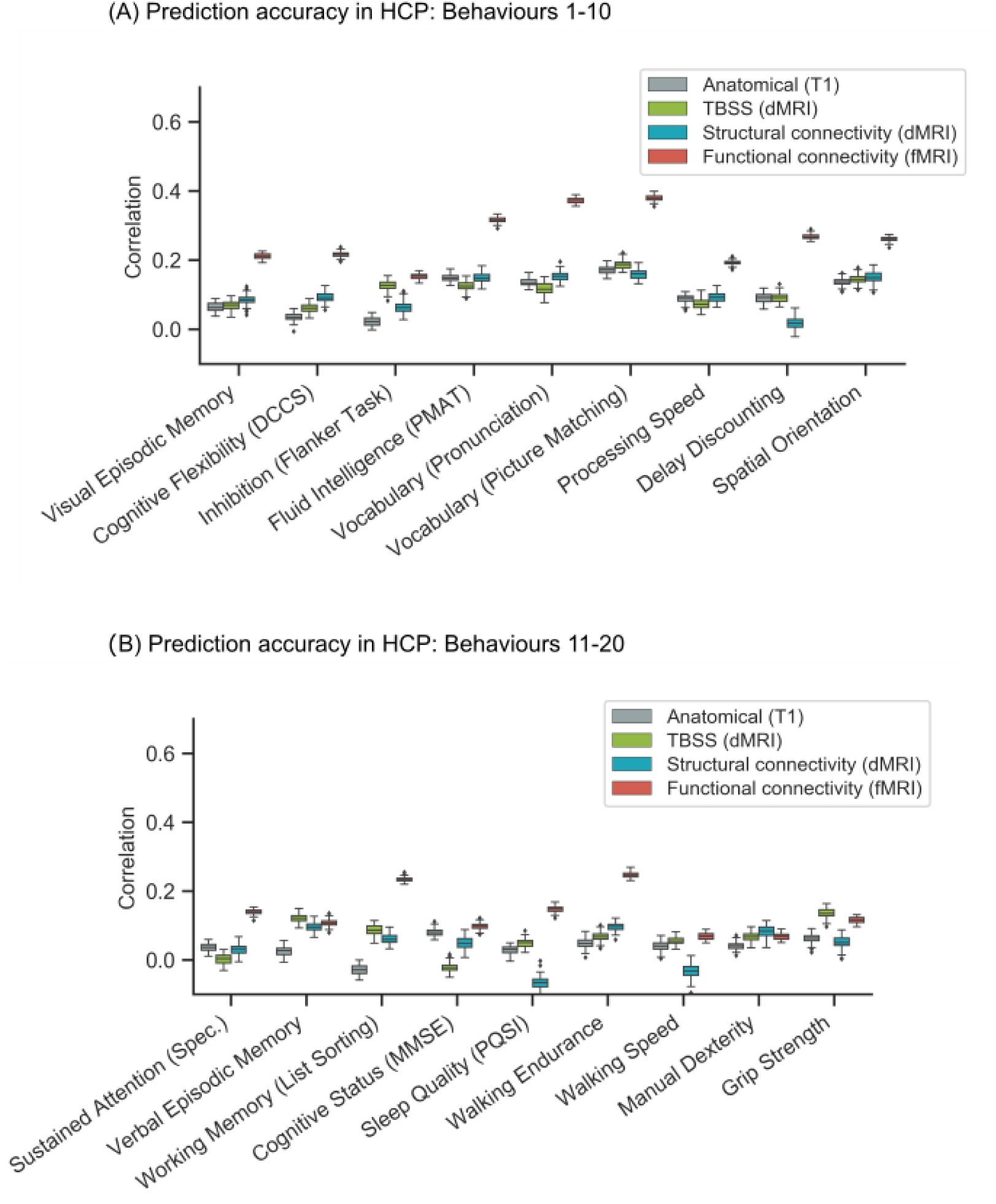
Prediction performance (Pearson’s correlation) of kernel ridge regression (KRR) for individual behavioural measures in the HCP dataset. (A) Prediction performance (Pearson’s correlation) of KRR averaged across single-feature-type predictive models within each modality (anatomical, TBSS, structural connectivity, functional connectivity) in the HCP dataset. Results are shown for behaviors 1-10 of Table S1. Each boxplot shows the distribution of performance over 60 repetitions of the nested cross-validation procedure. (B) Prediction performance (Pearson’s correlation) of KRR averaged across single-feature-type predictive models within each modality (anatomical, TBSS, structural connectivity, functional connectivity) in the HCP dataset. Results are shown for behaviors 11-21 of Table S1. Each boxplot shows the distribution of performance over 60 repetitions of the nested cross-validation procedure.

**Figure S3.**
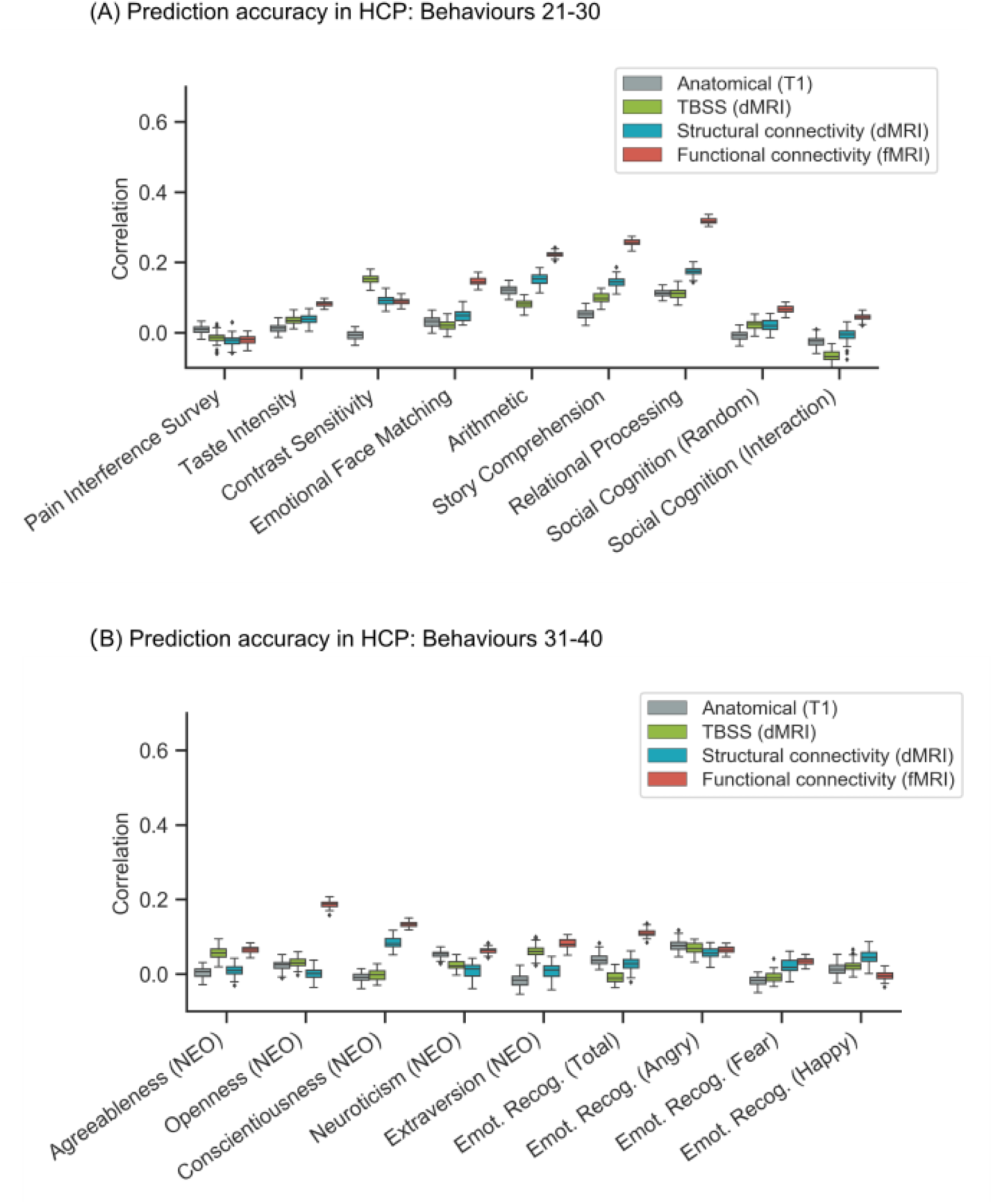
Prediction performance (Pearson’s correlation) of kernel ridge regression (KRR) for individual behavioural measures in the HCP dataset. (A) Prediction performance (Pearson’s correlation) of KRR averaged across single-feature-type predictive models within each modality (anatomical, TBSS, structural connectivity, functional connectivity) in the HCP dataset. Results are shown for behaviors 21-30 of Table S1. Each boxplot shows the distribution of performance over 60 repetitions of the nested cross-validation procedure. (B) Prediction performance (Pearson’s correlation) of KRR averaged across single-feature-type predictive models within each modality (anatomical, TBSS, structural connectivity, functional connectivity) in the HCP dataset. Results are shown for behaviors 31-40 of Table S1. Each boxplot shows the distribution of performance over 60 repetitions of the nested cross-validation procedure.

**Figure S4.**
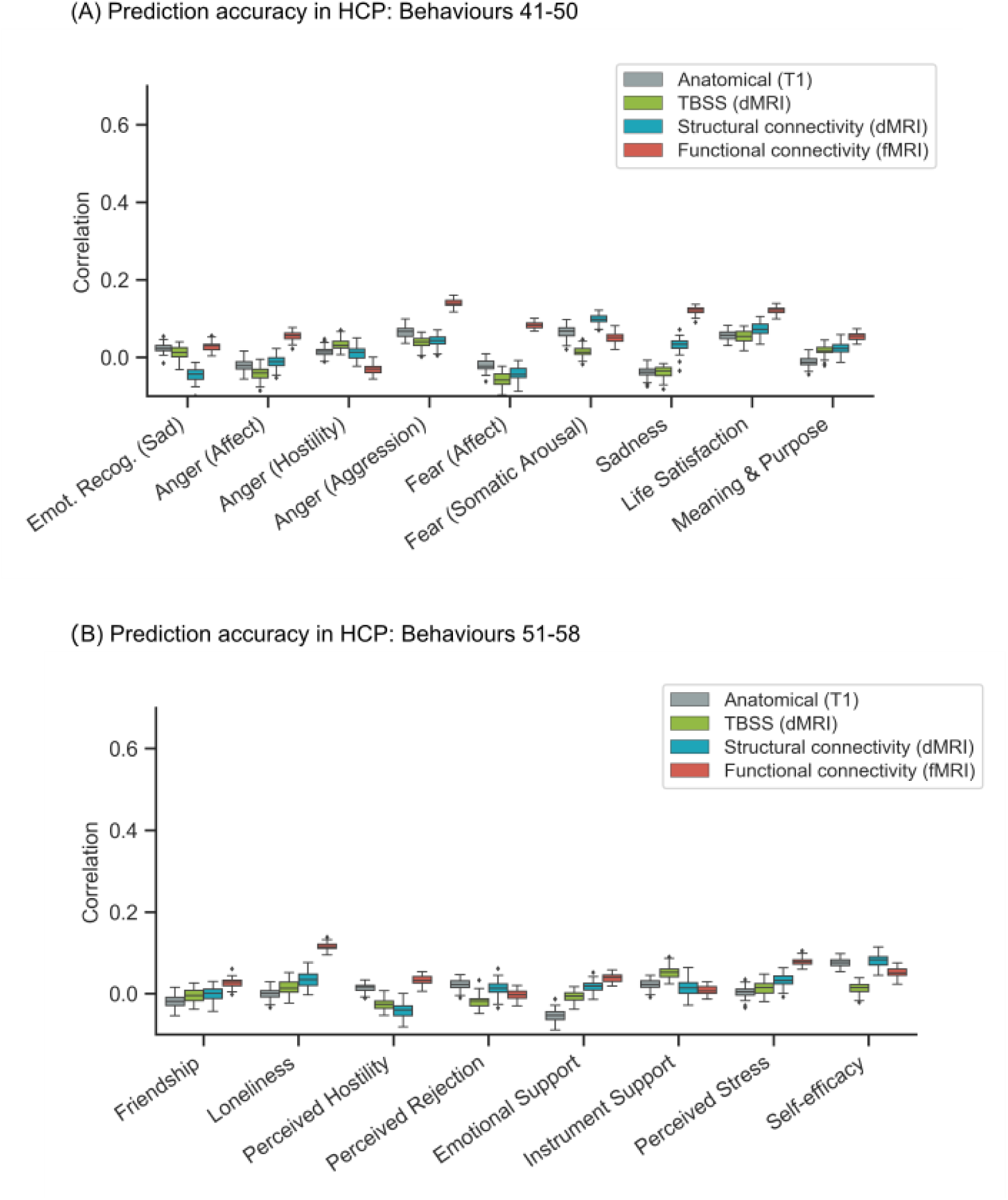
(Prediction performance (Pearson’s correlation) of kernel ridge regression (KRR) for individual behavioural measures in the HCP dataset. (A) Prediction performance (Pearson’s correlation) of KRR averaged across single-feature-type predictive models within each modality (anatomical, TBSS, structural connectivity, functional connectivity) in the HCP dataset. Results are shown for behaviors 41-50 of Table S1. Each boxplot shows the distribution of performance over 60 repetitions of the nested cross-validation procedure. (B) Prediction performance (Pearson’s correlation) of KRR averaged across single-feature-type predictive models within each modality (anatomical, TBSS, structural connectivity, functional connectivity) in the HCP dataset. Results are shown for behaviors 51-58 of Table S1. Each boxplot shows the distribution of performance over 60 repetitions of the nested cross-validation procedure.

**Figure S5.**
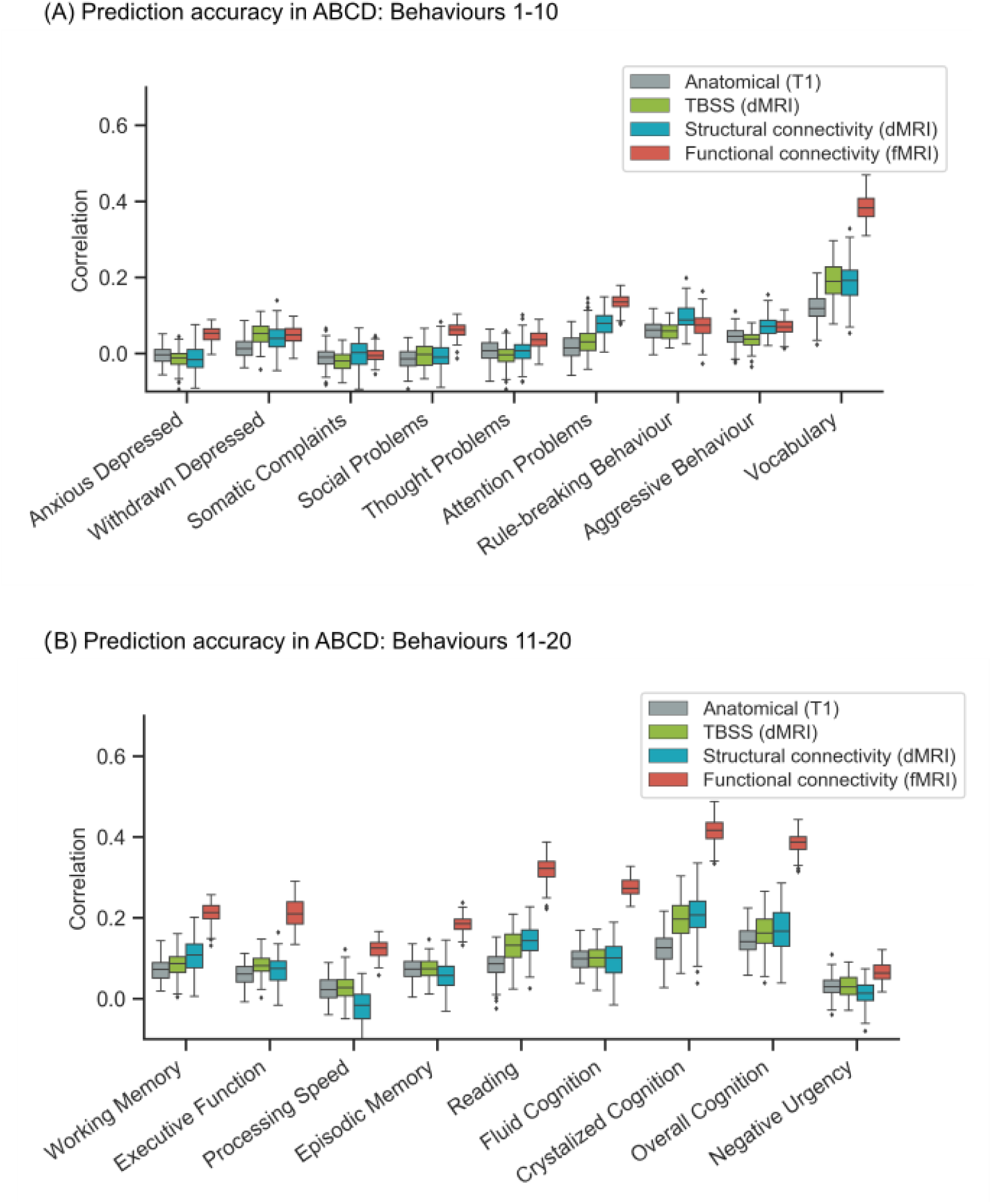
Prediction performance (Pearson’s correlation) of kernel ridge regression (KRR) for individual behavioural measures in the ABCD dataset. (A) Prediction performance (Pearson’s correlation) of KRR averaged across single-feature-type predictive models within each modality (anatomical, TBSS, structural connectivity, functional connectivity) in the ABCD dataset. Results are shown for behaviors 1-10 of Table S2. Each boxplot shows the distribution of performance over 120 repetitions of the nested cross-validation procedure. (B) Prediction performance (Pearson’s correlation) of KRR averaged across single-feature-type predictive models within each modality (anatomical, TBSS, structural connectivity, functional connectivity) in the ABCD dataset. Results are shown for behaviors 11-20 of Table S2. Each boxplot shows the distribution of performance over 120 repetitions of the nested cross-validation procedure.

**Figure S6.**
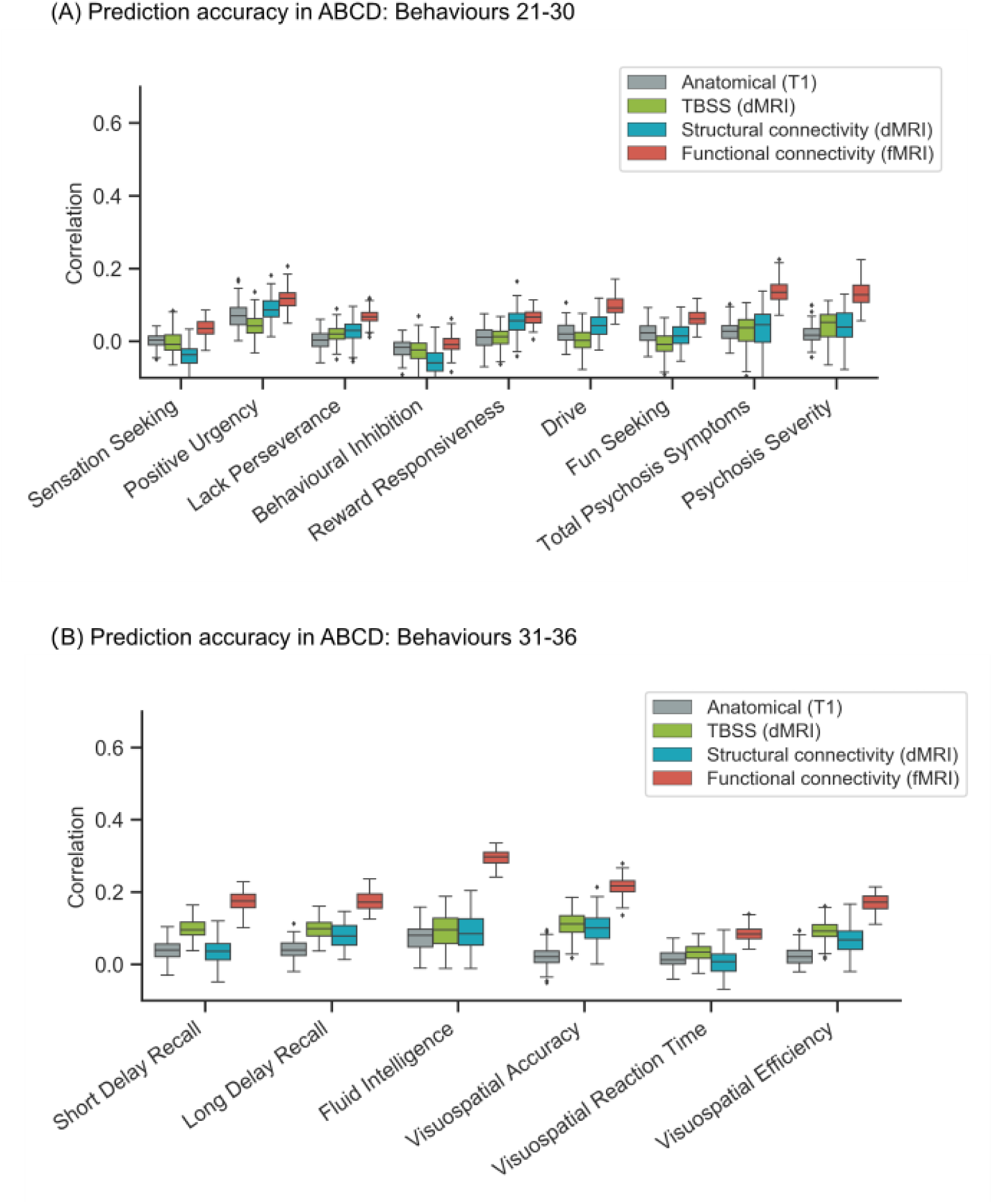
Prediction performance (Pearson’s correlation) of kernel ridge regression (KRR) for individual behavioural measures in the ABCD dataset. (A) Prediction performance (Pearson’s correlation) of KRR averaged across single-feature-type predictive models within each modality (anatomical, TBSS, structural connectivity, functional connectivity) in the ABCD dataset. Results are shown for behaviors 21-30 of Table S2. Each boxplot shows the distribution of performance over 120 repetitions of the nested cross-validation procedure. (B) Prediction performance (Pearson’s correlation) of KRR averaged across single-feature-type predictive models within each modality (anatomical, TBSS, structural connectivity, functional connectivity) in the ABCD dataset. Results are shown for behaviors 31-36 of Table S2. Each boxplot shows the distribution of performance over 120 repetitions of the nested cross-validation procedure.

**Figure S7.**
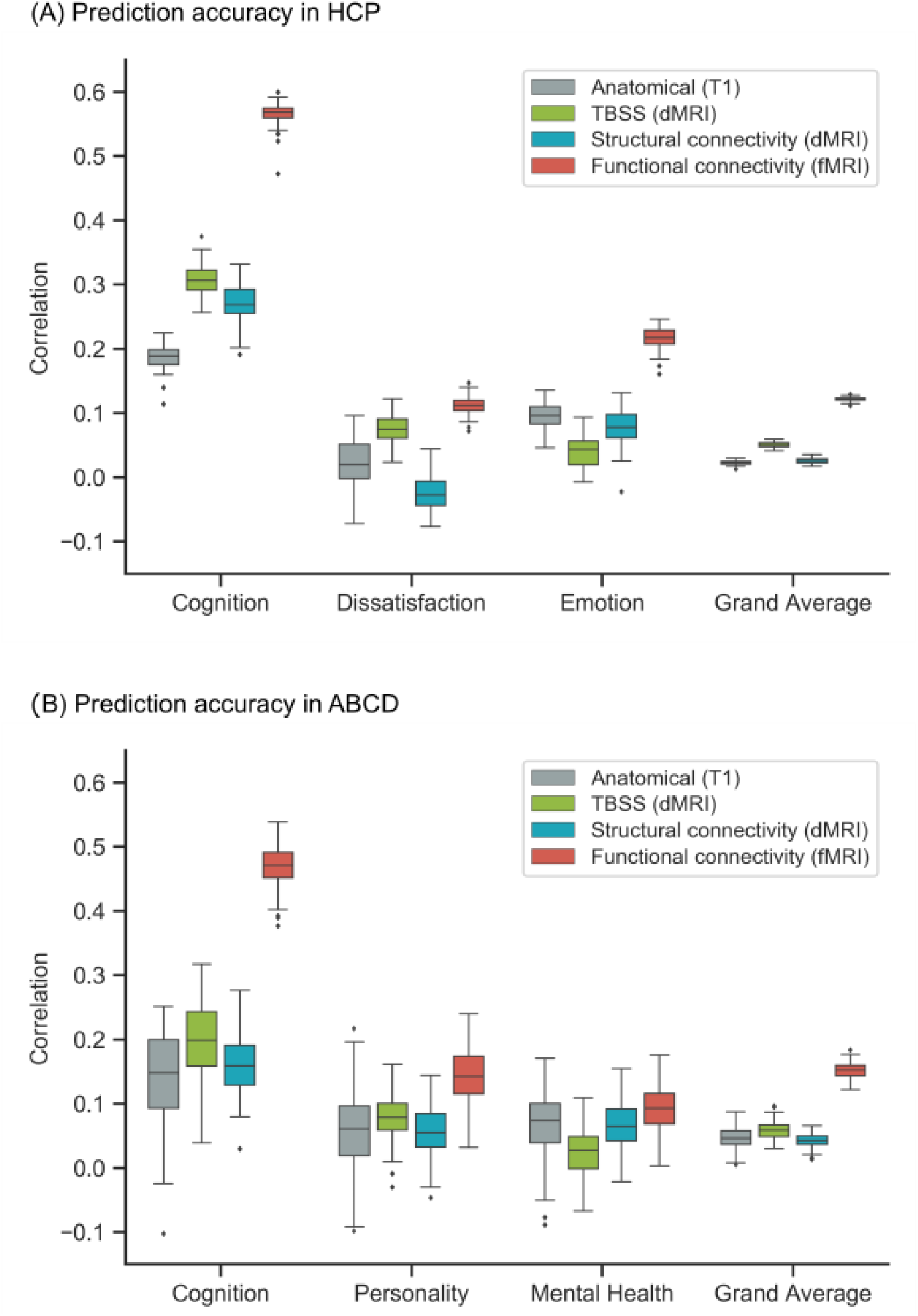
Functional connectivity (FC) outperforms other modalities for linear ridge regression (LRR). Figure is the same as Figure 4 except that LRR was utilized instead of kernel ridge regression. (A) Prediction performance (Pearson’s correlation) of LRR for the best performing feature-type within each modality in the HCP dataset. For the cognition component, the best features were cortical area, TBSS OD, SC FA and language FC. For the dissatisfaction component, the best features were cortical thickness, TBSS FA, SC stream count and working memory FC. For the emotion component, the best features were cortical volume, TBSS AD, SC FA and social cognition FC. For the grand average, the best features were cortical volume, TBSS AD, SC AD and language FC. (B) Prediction performance (Pearson’s correlation) of LRR for the best performing feature-type within each modality in the ABCD dataset. For the cognition component, the best features were cortical area, TBSS OD, SC ICVF and N-back FC. For the personality component, the best features were cortical volume, TBSS OD, SC MD and MID FC. For the mental health component, the best features were cortical area, TBSS OD, SC MD and resting FC. For the grand average, the best features were cortical area, TBSS OD, SC ICVF and N-back FC.

**Fig S8.**
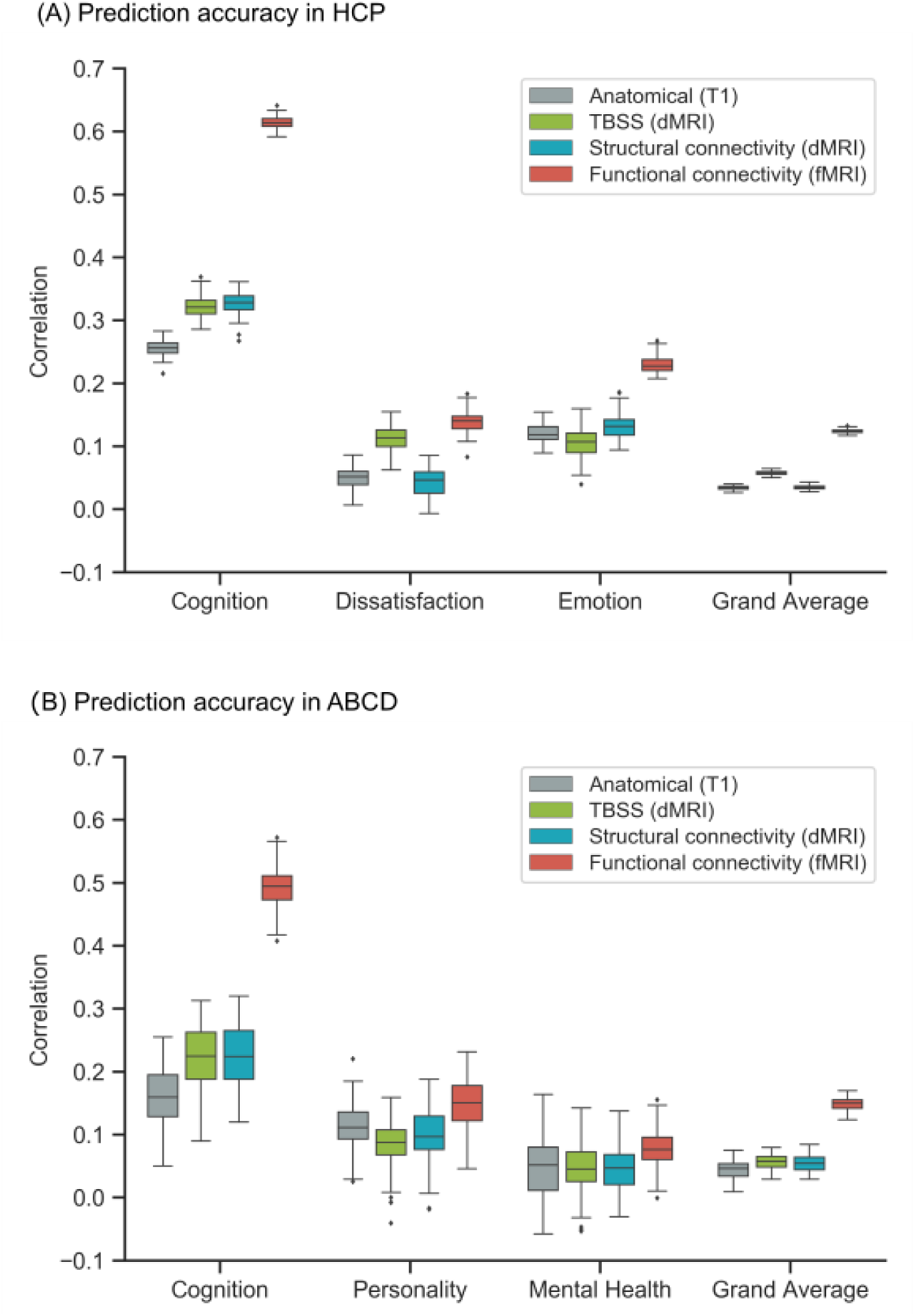
Functional connectivity (FC) outperforms other modalities for elastic net. Figure is the same as Figure 4 except that elastic net was utilized instead of kernel ridge regression. (A) Prediction performance (Pearson’s correlation) of elastic net for the best performing feature-type within each modality in the HCP dataset. For the cognition component, the best features were cortical area, TBSS FA, SC FA and language FC. For the dissatisfaction component, the best features were cortical thickness, TBSS AD, SC stream length and working memory FC. For the emotion component, the best features were cortical volume, TBSS OD, SC FA and language FC. For the grand average, the best features were cortical thickness, TBSS FA, SC AD and language FC. (B) Prediction performance (Pearson’s correlation) of elastic net for the best performing feature-type within each modality in the ABCD dataset. For the cognition component, the best features were cortical thickness, TBSS OD, SC ICVF and N-back FC. For the personality component, the best features were cortical thickness, TBSS OD, SC OD and MID FC. For the mental health component, the best features were cortical area, TBSS OD, SC OD and SST FC. For the grand average, the best features were cortical area, TBSS OD, SC ICVF and N-back FC.

**Figure S9.**
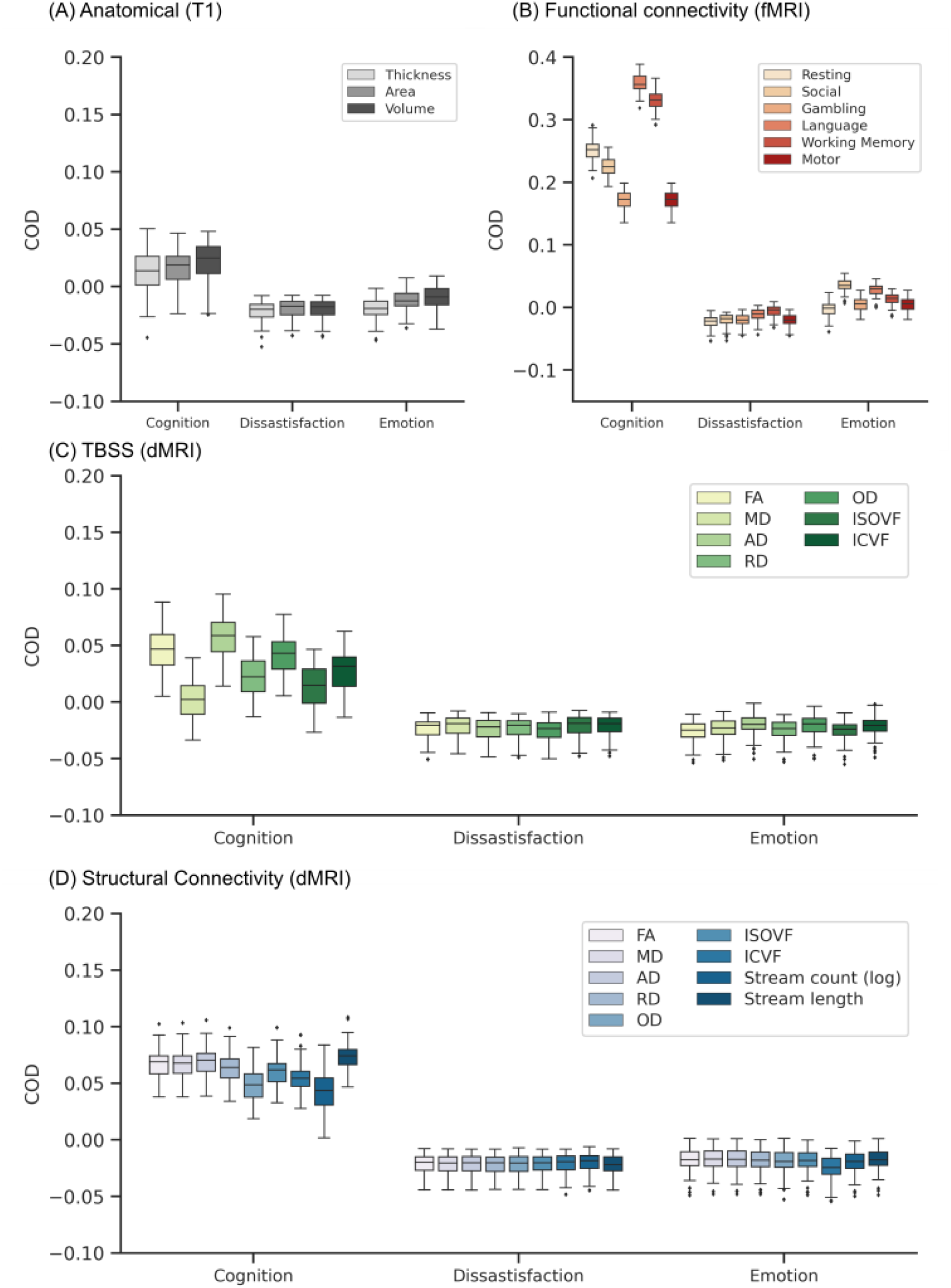
Prediction performance (COD) of kernel ridge regression (KRR) for each single-feature-type in the HCP dataset. Figure is the same as Figure 5, except that COD was shown instead of Pearson’s correlation. Results are shown separately for (A) anatomical features, (B) FC, (C) TBSS and (D) structural connectivity.

**Figure S10.**
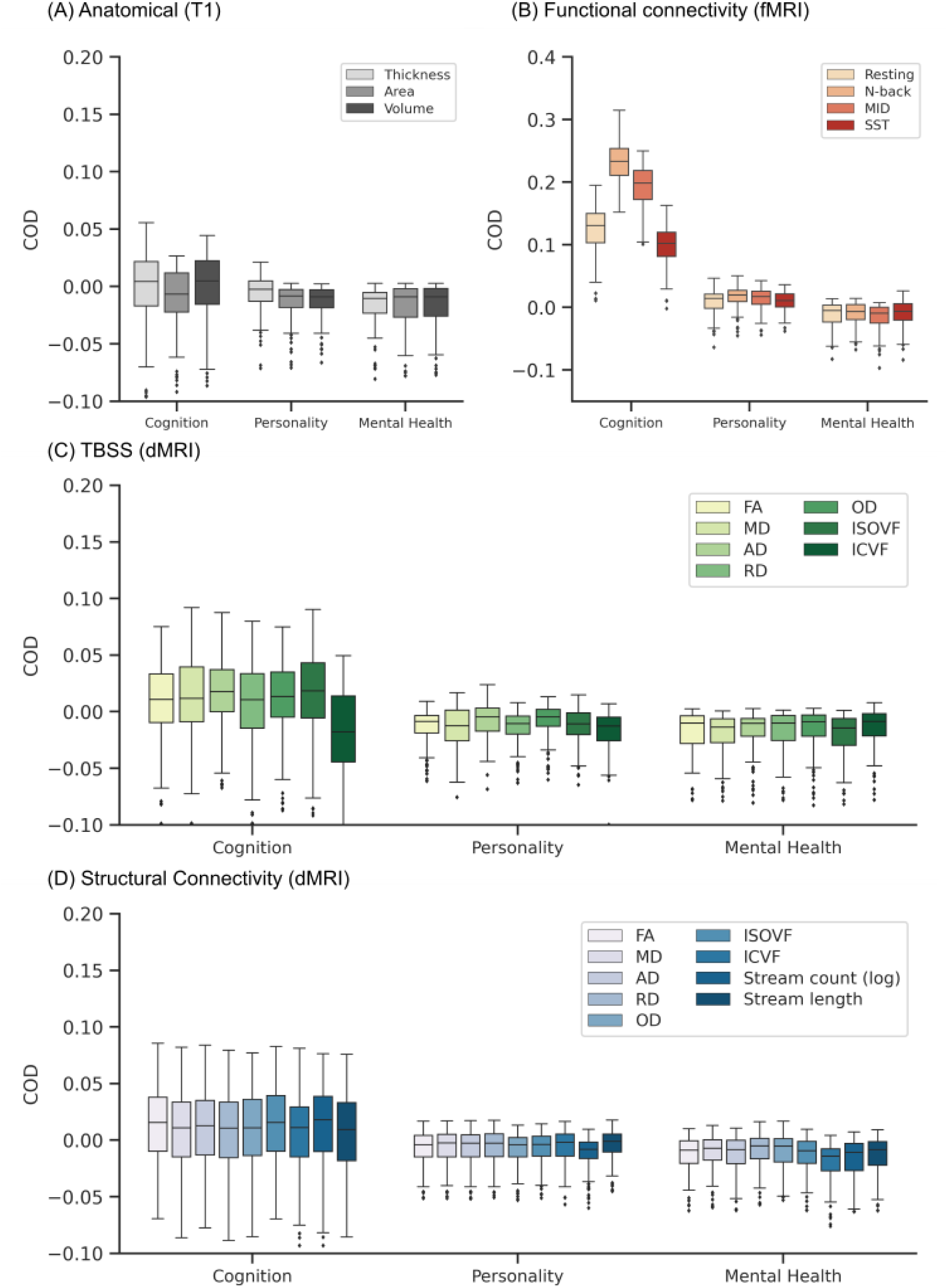
Prediction performance (COD) of kernel ridge regression (KRR) for each single-feature-type in the ABCD dataset. Figure is the same as Figure 6, except that COD was shown instead of Pearson’s correlation. Results are shown separately for (A) anatomical features, (B) FC, (C) TBSS and (D) structural connectivity.

**Figure S11.**
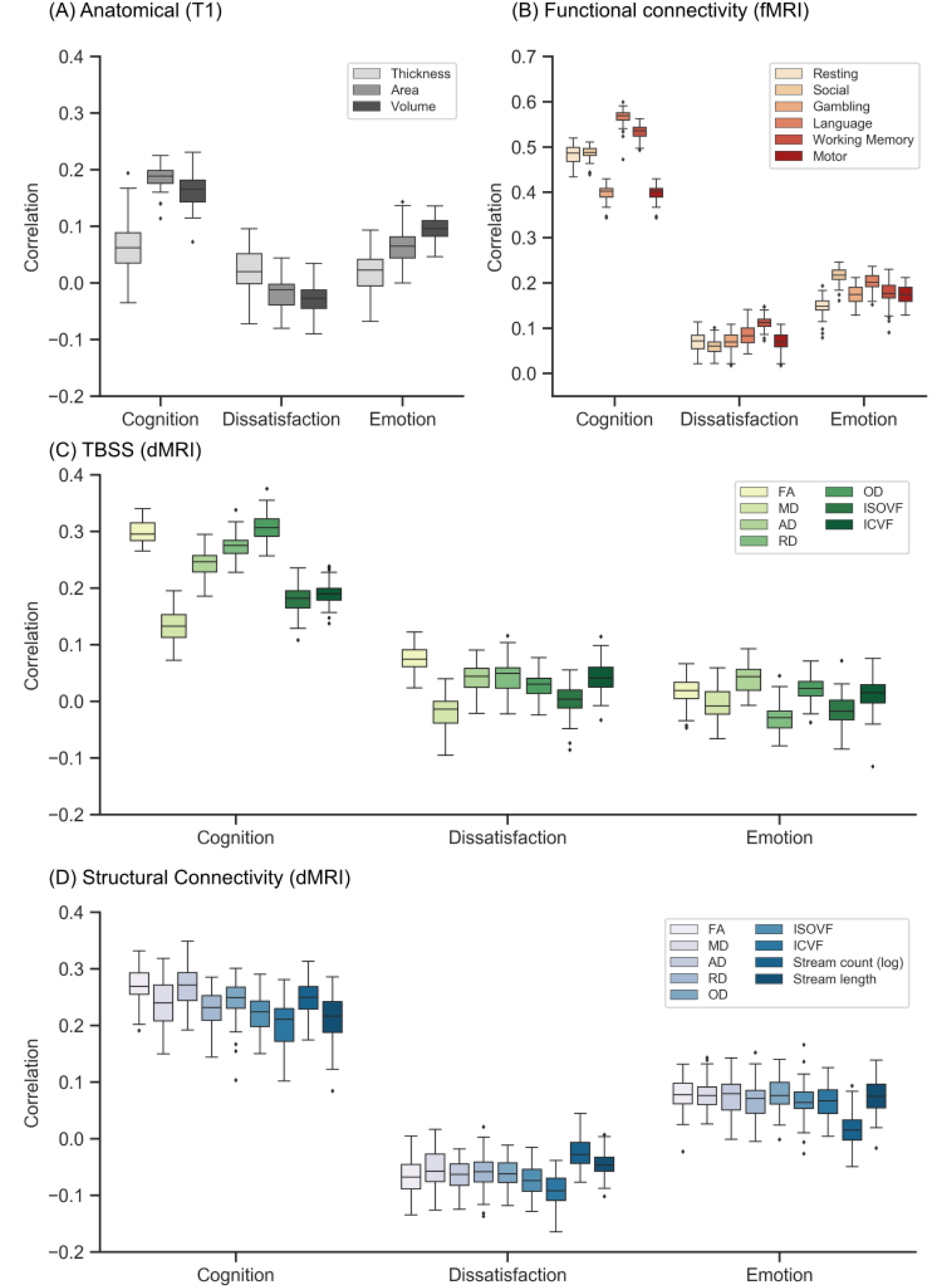
Prediction performance (Pearson’s correlation) of linear ridge regression (LRR) for each single-feature-type in the HCP dataset. Figure is the same as Figure 5 except that LRR was utilized instead of kernel ridge regression. Results are shown separately for (A) anatomical features, (B) FC, (C) TBSS and (D) structural connectivity.

**Figure S12.**
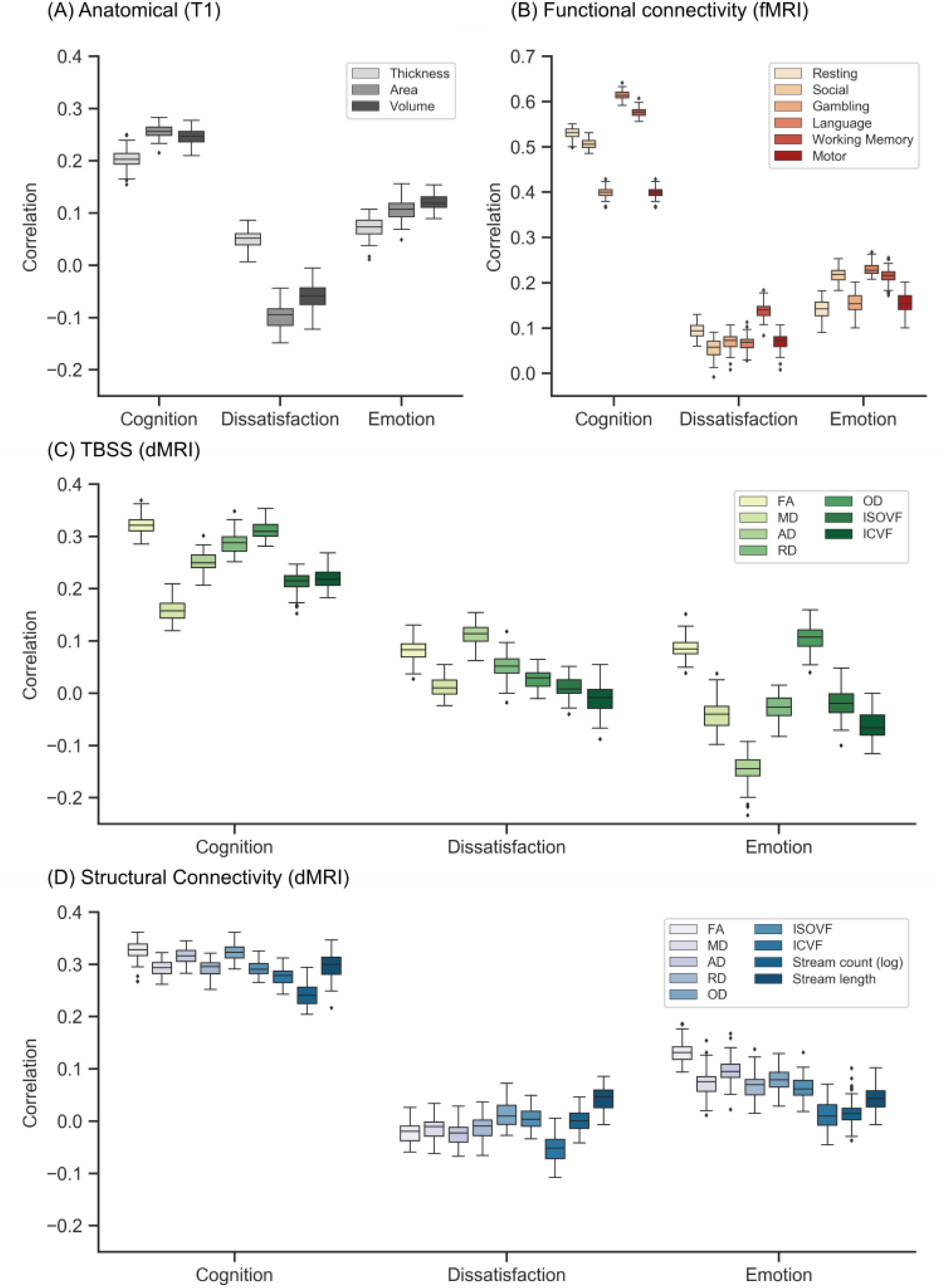
Prediction performance (Pearson’s correlation) of elastic net for each single-feature-type in the HCP dataset. Figure is the same as Figure 5 except that elastic net was utilized instead of kernel ridge regression. Results are shown separately for (A) anatomical features, (B) FC, (C) TBSS and (D) structural connectivity.

**Figure S13.**
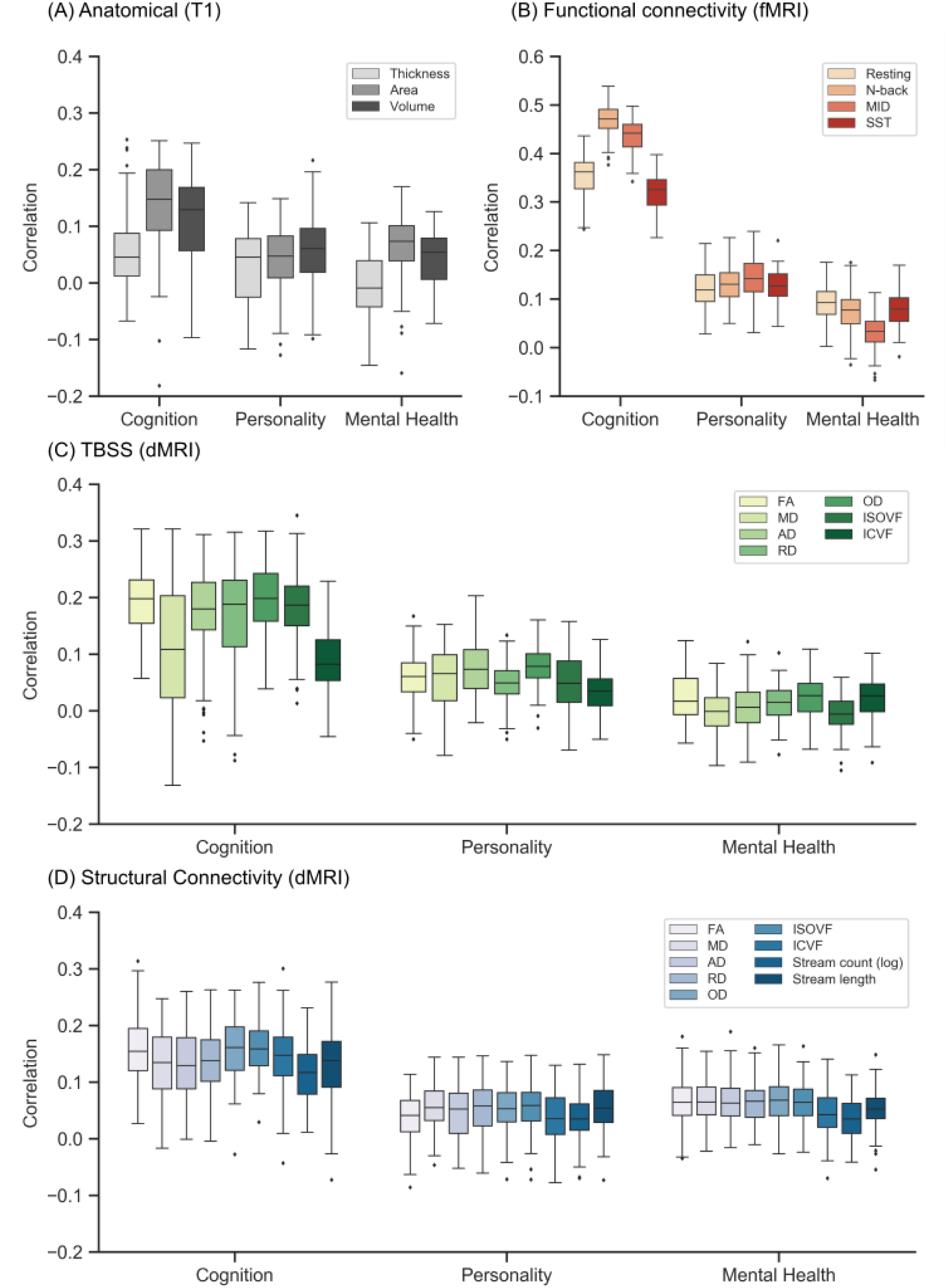
Prediction performance (Pearson’s correlation) of linear ridge regression (LRR) for each single-feature-type in the ABCD dataset. Figure is the same as Figure 6 except that LRR was utilized instead of kernel ridge regression. Results are shown separately for (A) anatomical features, (B) FC, (C) TBSS and (D) structural connectivity.

**Figure S14.**
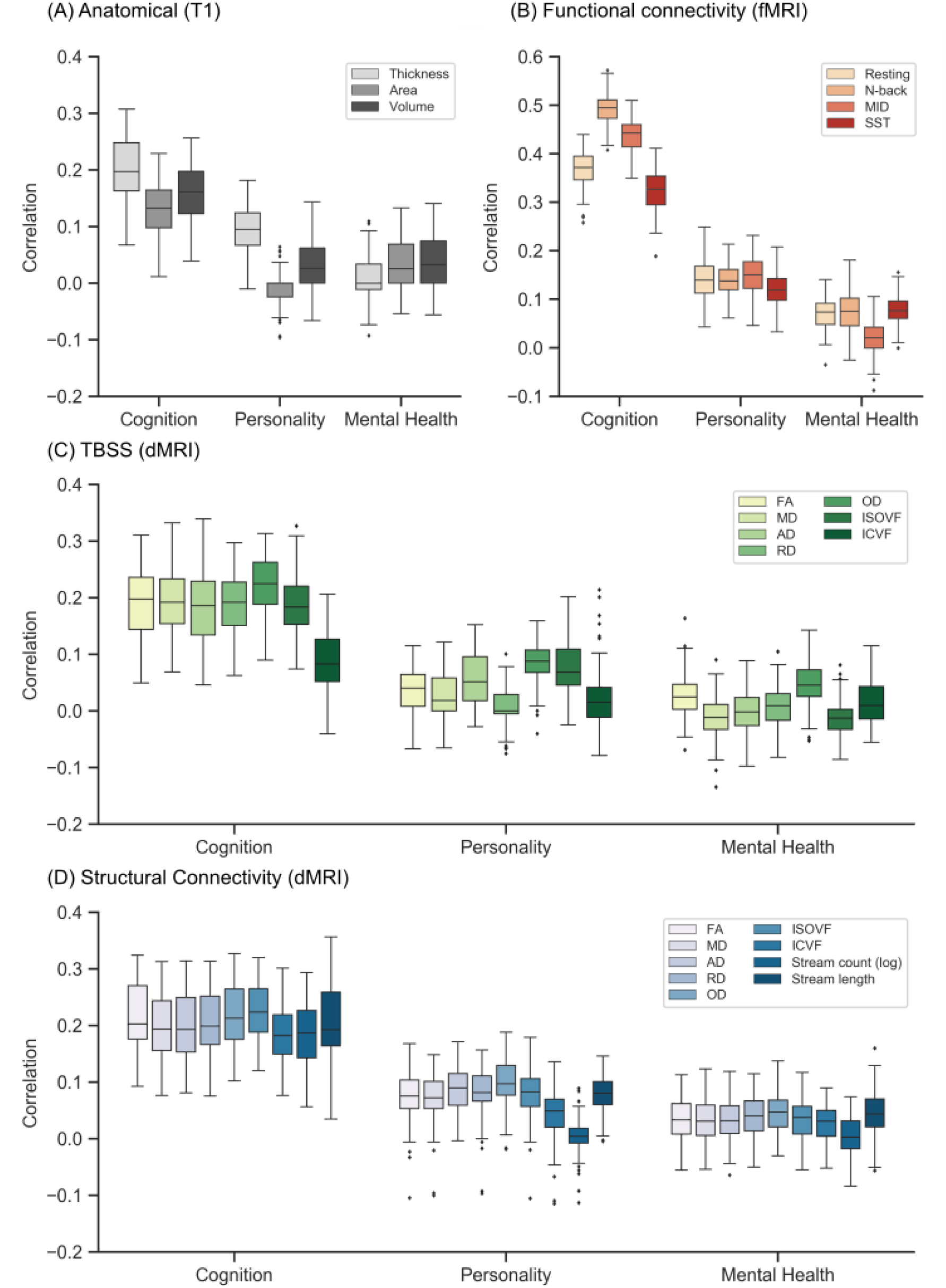
Prediction performance (Pearson’s correlation) of elastic net for each single-feature-type in the ABCD dataset. Figure is the same as Figure 6 except that elastic net was utilized instead of kernel ridge regression. Results are shown separately for (A) anatomical features, (B) FC, (C) TBSS and (D) structural connectivity.

**Figure S15.**
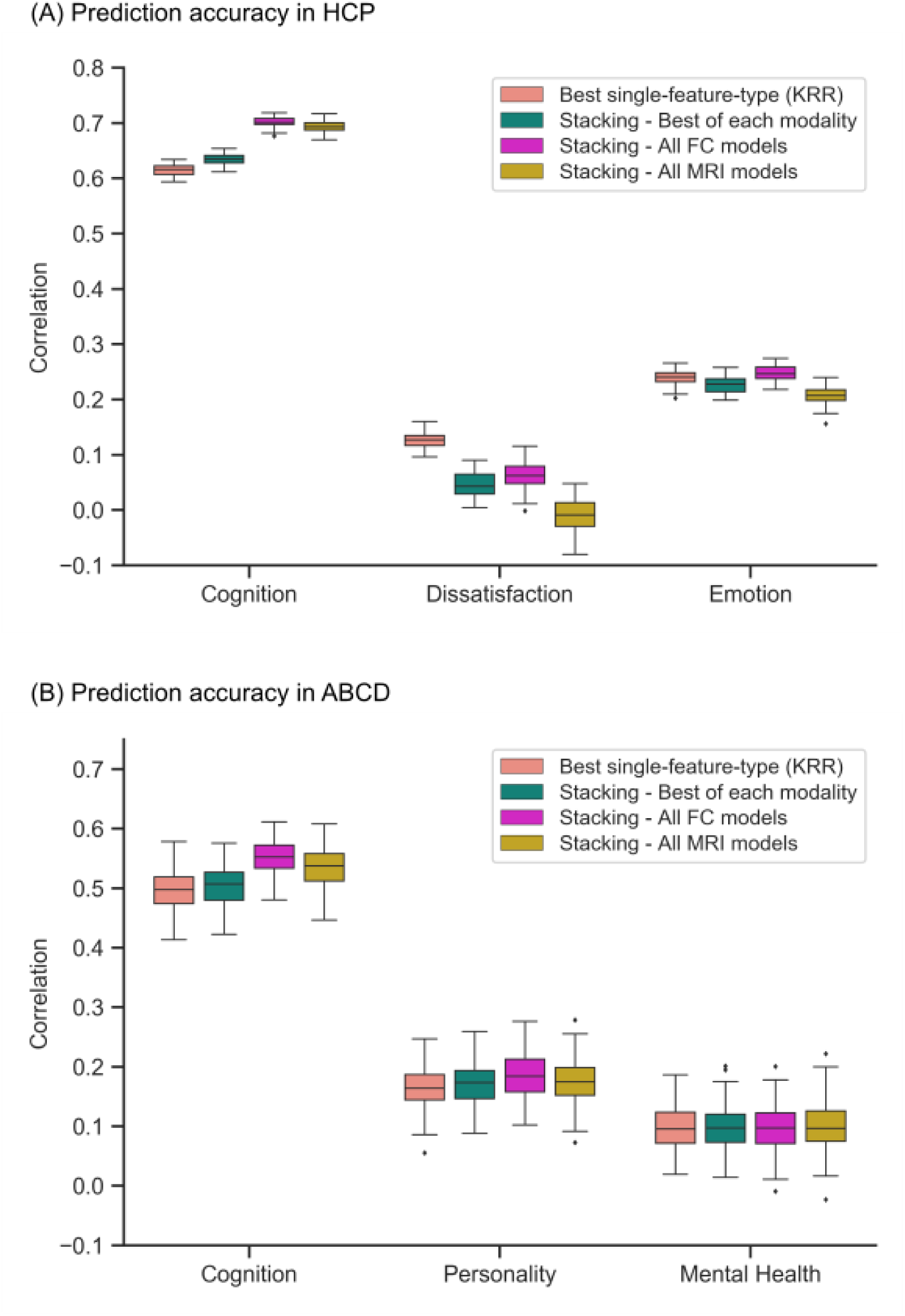
Combining resting and task FC was as good as combining across all modalities, or combining the best single-feature-type models of each modality. Figure is the same as Figure 7, except that the results of multi-KRR are replaced with the results of stacking the best single-feature-type models (which can be found in Figure 4). (A) Prediction performance (Pearson’s correlation) from combining various MRI features and modalities in the HCP dataset. We considered stacking the best single-feature-type model of each modality, all FC models, and all single-feature-type models across all modalities. For comparison, the best single-feature-type from KRR is shown. Each boxplot shows the distribution over 60 repetitions of the nested cross-validation procedure. (B) Prediction performance (Pearson’s correlation) from combining various MRI features and modalities in the ABCD dataset. We considered stacking the best single-feature-type model of each modality, all FC models, and all single-feature-type models across all modalities. For comparison, the best single-feature-type from KRR is shown. Each boxplot shows the distribution over 120 repetitions of the nested cross-validation procedure.

**Figure S16.**
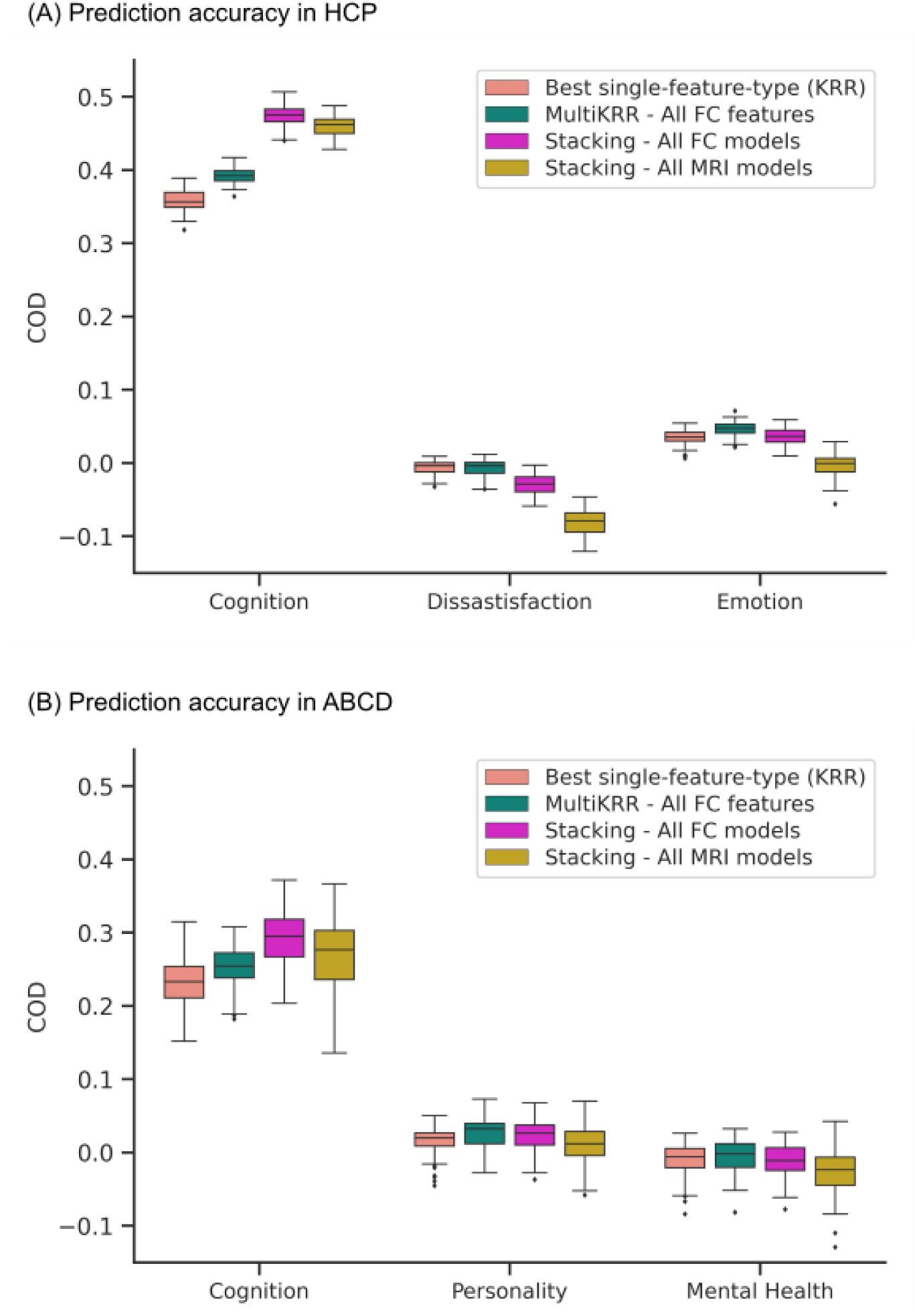
Combining resting and task FC was as good as combining across all modalities. Figure is the same as Figure 7, except that COD was shown instead of Pearson’s correlation. (A) Prediction performance (COD) from combining various MRI features and modalities in the HCP dataset. We considered multi-KRR of all FC features, stacking of all FC models and stacking of all single-feature-type models across all modalities. For comparison, the best single-feature-type from KRR is shown. Each boxplot shows the distribution over 60 repetitions of the nested cross-validation procedure. (B) Prediction performance (COD) from combining various MRI features and modalities in the ABCD dataset. We considered multi-KRR of all FC features, stacking of all FC models and stacking of all single-feature-type models across all modalities. For comparison, the best single-feature-type from KRR is shown. Each boxplot shows the distribution over 120 repetitions of the nested cross-validation procedure.

## Footnotes

1 Note to reviewers: The NDA link will only be public after the manuscript is published, since we will not be able to change the relevant information (e.g. reference to this study) after the link becomes public. However, we have structured the HCP and ABCD data to be as similar as possible.

## References

Alnæs, D., Kaufmann, T., Marquand, A. F., Smith, S. M., & Westlye, L. T. (2020). Patterns of sociocognitive stratification and perinatal risk in the child brain. Proceedings of the National Academy of Sciences, 117(22), 12419. https://doi.org/10.1073/pnas.2001517117

Arbabshirani, M. R., Plis, S., Sui, J., & Calhoun, V. D. (2017). Single subject prediction of brain disorders in neuroimaging: Promises and pitfalls. NeuroImage, 145(Pt B), 137–165. https://doi.org/10.1016/j.neuroimage.2016.02.079

Avinun, R., Israel, S., Knodt, A. R., & Hariri, A. R. (2020). Little evidence for associations between the Big Five personality traits and variability in brain gray or white matter. NeuroImage, 220, 117092. https://doi.org/10.1016/j.neuroimage.2020.117092

Bajaj, S., Krismer, F., Palma, J.-A., Wenning, G. K., Kaufmann, H., Poewe, W., & Seppi, K. (2017). Diffusion-weighted MRI distinguishes Parkinson disease from the parkinsonian variant of multiple system atrophy: A systematic review and meta-analysis. PLoS ONE, 12(12), e0189897. https://doi.org/10.1371/journal.pone.0189897

Basser, P. J., Mattiello, J., & Lebihan, D. (1994). MR diffusion tensor spectroscopy and imaging. Biophysical Journal, 66(1), 259–267. https://doi.org/10.1016/s0006-3495(94)80775-1

Beyer, L., Olivier, Alexander, & A\“aron. (2020). Are we done with ImageNet? arXiv pre-print server. https://doi.org/None arxiv:2006.07159

Bouckaert, R. R., & Frank, E. (2004). Evaluating the Replicability of Significance Tests for Comparing Learning Algorithms. In (pp. 3–12). Springer Berlin Heidelberg. https://doi.org/10.1007/978-3-540-24775-3_3

Bzdok, D., & Meyer-Lindenberg, A. (2018). Machine Learning for Precision Psychiatry: Opportunities and Challenges. Biological Psychiatry: Cognitive Neuroscience and Neuroimaging, 3(3), 223–230. https://doi.org/10.1016/j.bpsc.2017.11.007

Cai, H., Zhu, J., & Yu, Y. (2020). Robust prediction of individual personality from brain functional connectome. Social Cognitive and Affective Neuroscience, 15(3), 359–369. https://doi.org/10.1093/scan/nsaa044

Calhoun, V. (2018). Data-driven approaches for identifying links between brain structure and function in health and disease. Neurocircuitry, 20(2), 87–99. https://doi.org/10.31887/dcns.2018.20.2/vcalhoun

Casey, B. J., Cannonier, T., Conley, M. I., Cohen, A. O., Barch, D. M., Heitzeg, M. M., Soules, M. E., Teslovich, T., Dellarco, D. V., Garavan, H., Orr, C. A., Wager, T. D., Banich, M. T., Speer, N. K., Sutherland, M. T., Riedel, M. C., Dick, A. S., Bjork, J. M., Thomas, K. M., Chaarani, B., Mejia, M. H., Hagler, D. J., Daniela Cornejo, M., Sicat, C. S., Harms, M. P., Dosenbach, N. U. F., Rosenberg, M., Earl, E., Bartsch, H., Watts, R., Polimeni, J. R., Kuperman, J. M., Fair, D. A., & Dale, A. M. (2018). The Adolescent Brain Cognitive Development (ABCD) study: Imaging acquisition across 21 sites. Developmental Cognitive Neuroscience, 32, 43–54. https://doi.org/https://doi.org/10.1016/j.dcn.2018.03.001

Chen, J., Tam, A., Kebets, V., Orban, C., Ooi, L. Q. R., Marek, S., Dosenbach, N., Eickhoff, S., Bzdok, D., Holmes, A. J., & Thomas Yeo, B. T. (2020). Shared and unique brain network features predict cognition, personality and mental health in childhood. Cold Spring Harbor Laboratory. https://dx.doi.org/10.1101/2020.06.24.168724

Cohen, S. E., Zantvoord, J. B., Wezenberg, B. N., Bockting, C. L. H., & Van Wingen, G. A. (2021). Magnetic resonance imaging for individual prediction of treatment response in major depressive disorder: a systematic review and meta-analysis. Translational Psychiatry, 11(1). https://doi.org/10.1038/s41398-021-01286-x

Daducci, A., Canales-Rodríguez, E. J., Zhang, H., Dyrby, T. B., Alexander, D. C., & Thiran, J.-P. (2015). Accelerated Microstructure Imaging via Convex Optimization (AMICO) from diffusion MRI data. NeuroImage, 105, 32–44. https://doi.org/10.1016/j.neuroimage.2014.10.026

Dale, A. M., Fischl, B., & Sereno, M. I. (1999). Cortical Surface-Based Analysis. NeuroImage, 9(2), 179–194. https://doi.org/10.1006/nimg.1998.0395

Dhamala, E., Jamison, K. W., Jaywant, A., Dennis, S., & Kuceyeski, A. (2021). Distinct functional and structural connections predict crystallised and fluid cognition in healthy adults. Human Brain Mapping, 42(10), 3102–3118. https://doi.org/10.1002/hbm.25420

Dubois, J., Galdi, P., Han, Y., Paul, L. K., & Adolphs, R. (2018). Resting-State Functional Brain Connectivity Best Predicts the Personality Dimension of Openness to Experience. Personality Neuroscience, 1. https://doi.org/10.1017/pen.2018.8

Elad, D., Cetin-Karayumak, S., Zhang, F., Cho, K. I. K., Lyall, A. E., Seitz-Holland, J., Ben-Ari, R., Pearlson, G. D., Tamminga, C. A., Sweeney, J. A., Clementz, B. A., Schretlen, D. J., Viher, P. V., Stegmayer, K., Walther, S., Lee, J., Crow, T. J., James, A., Voineskos, A. N., Buchanan, R. W., Szeszko, P. R., Malhotra, A. K., Keshavan, M. S., Shenton, M. E., Rathi, Y., Bouix, S., Sochen, N., Kubicki, M. R., & Pasternak, O. (2021). Improving the predictive potential of diffusion MRI in schizophrenia using normative models— Towards subject-level classification. Human Brain Mapping. https://doi.org/10.1002/hbm.25574

Elliott, M. L., Knodt, A. R., Cooke, M., Kim, M. J., Melzer, T. R., Keenan, R., Ireland, D., Ramrakha, S., Poulton, R., Caspi, A., Moffitt, T. E., & Hariri, A. R. (2019). General functional connectivity: Shared features of resting-state and task fMRI drive reliable and heritable individual differences in functional brain networks. NeuroImage, 189, 516–532. https://doi.org/10.1016/j.neuroimage.2019.01.068

Engemann, D. A., Kozynets, O., Sabbagh, D., Lemaître, G., Varoquaux, G., Liem, F., & Gramfort, A. (2020). Combining magnetoencephalography with magnetic resonance imaging enhances learning of surrogate-biomarkers. eLife, 9. https://doi.org/10.7554/elife.54055

Finn, E. S., Shen, X., Scheinost, D., Rosenberg, M. D., Huang, J., Chun, M. M., Papademetris, X., & Constable, R. T. (2015). Functional connectome fingerprinting: identifying individuals using patterns of brain connectivity. Nature Neuroscience, 18(11), 1664–1671. https://doi.org/10.1038/nn.4135

Friedman, J., Hastie, T., & Tibshirani, R. (2010). Regularization Paths for Generalized Linear Models via Coordinate Descent. Journal of statistical software, 33(1), 1–22. https://pubmed.ncbi.nlm.nih.gov/20808728 https://www.ncbi.nlm.nih.gov/pmc/articles/PMC2929880/

Gao, S., Greene, A. S., Constable, R. T., & Scheinost, D. (2019). Combining multiple connectomes improves predictive modeling of phenotypic measures. NeuroImage, 201, 116038. https://doi.org/https://doi.org/10.1016/j.neuroimage.2019.116038

Glasser, M. F., Sotiropoulos, S. N., Wilson, J. A., Coalson, T. S., Fischl, B., Andersson, J. L., Xu, J., Jbabdi, S., Webster, M., Polimeni, J. R., Van Essen, D. C., & Jenkinson, M. (2013). The minimal preprocessing pipelines for the Human Connectome Project. NeuroImage, 80, 105–124. https://doi.org/10.1016/j.neuroimage.2013.04.127

Greene, A. S., Gao, S., Scheinost, D., & Constable, R. T. (2018). Task-induced brain state manipulation improves prediction of individual traits. Nature Communications, 9(1). https://doi.org/10.1038/s41467-018-04920-3

Hagler, D. J., Hatton, S., Cornejo, M. D., Makowski, C., Fair, D. A., Dick, A. S., Sutherland, M. T., Casey, B. J., Barch, D. M., Harms, M. P., Watts, R., Bjork, J. M., Garavan, H. P., Hilmer, L., Pung, C. J., Sicat, C. S., Kuperman, J., Bartsch, H., Xue, F., Heitzeg, M. M., Laird, A. R., Trinh, T. T., Gonzalez, R., Tapert, S. F., Riedel, M. C., Squeglia, L. M., Hyde, L. W., Rosenberg, M. D., Earl, E. A., Howlett, K. D., Baker, F. C., Soules, M., Diaz, J., De Leon, O. R., Thompson, W. K., Neale, M. C., Herting, M., Sowell, E. R., Alvarez, R. P., Hawes, S. W., Sanchez, M., Bodurka, J., Breslin, F. J., Morris, A. S., Paulus, M. P., Simmons, W. K., Polimeni, J. R., Van Der Kouwe, A., Nencka, A. S., Gray, K. M., Pierpaoli, C., Matochik, J. A., Noronha, A., Aklin, W. M., Conway, K., Glantz, M., Hoffman, E., Little, R., Lopez, M., Pariyadath, V., Weiss, S. R., Wolff-Hughes, D. L., Delcarmen-Wiggins, R., Feldstein Ewing, S. W., Miranda-Dominguez, O., Nagel, B. J., Perrone, A. J., Sturgeon, D. T., Goldstone, A., Pfefferbaum, A., Pohl, K. M., Prouty, D., Uban, K., Bookheimer, S. Y., Dapretto, M., Galvan, A., Bagot, K., Giedd, J., Infante, M. A., Jacobus, J., Patrick, K., Shilling, P. D., Desikan, R., Li, Y., Sugrue, L., Banich, M. T., Friedman, N., Hewitt, J. K., Hopfer, C., Sakai, J., Tanabe, J., Cottler, L. B., Nixon, S. J., Chang, L., Cloak, C., Ernst, T., Reeves, G., Kennedy, D. N., Heeringa, S., Peltier, S., Schulenberg, J., Sripada, C., Zucker, R. A., Iacono, W. G., Luciana, M., Calabro, F. J., Clark, D. B., Lewis, D. A., Luna, B., Schirda, C., Brima, T., Foxe, J. J., Freedman, E. G., Mruzek, D. W., Mason, M. J., Huber, R., McGlade, E., Prescot, A., Renshaw, P. F., Yurgelun-Todd, D. A., Allgaier, N. A., Dumas, J. A., Ivanova, M., Potter, A., Florsheim, P., Larson, C., Lisdahl, K., Charness, M. E., Fuemmeler, B., Hettema, J. M., Maes, H. H., Steinberg, J., Anokhin, A. P., Glaser, P., Heath, A. C., Madden, P. A., Baskin-Sommers, A., Constable, R. T., Grant, S. J., Dowling, G. J., Brown, S. A., Jernigan, T. L., & Dale, A. M. (2019). Image processing and analysis methods for the Adolescent Brain Cognitive Development Study. NeuroImage, 202, 116091. https://doi.org/10.1016/j.neuroimage.2019.116091

He, T., Kong, R., Holmes, A. J., Nguyen, M., Sabuncu, M. R., Eickhoff, S. B., Bzdok, D., Feng, J., & Yeo, B. T. T. (2020). Deep neural networks and kernel regression achieve comparable accuracies for functional connectivity prediction of behavior and demographics. NeuroImage, 206, 116276. https://doi.org/https://doi.org/10.1016/j.neuroimage.2019.116276

Jiang, R., Zuo, N., Ford, J. M., Qi, S., Zhi, D., Zhuo, C., Xu, Y., Fu, Z., Bustillo, J., Turner, J. A., Calhoun, V. D., & Sui, J. (2020). Task-induced brain connectivity promotes the detection of individual differences in brain-behavior relationships. NeuroImage, 207, 116370. https://doi.org/https://doi.org/10.1016/j.neuroimage.2019.116370

Kaiser, H. F. (1958). The varimax criterion for analytic rotation in factor analysis. Psychometrika, 23(3), 187–200. https://doi.org/10.1007/BF02289233

Kebets, V., Holmes, A. J., Orban, C., Tang, S., Li, J., Sun, N., Kong, R., Poldrack, R. A., & Yeo, B. T. T. (2019). Somatosensory-Motor Dysconnectivity Spans Multiple Transdiagnostic Dimensions of Psychopathology. Biological Psychiatry, 86(10), 779–791. https://doi.org/https://doi.org/10.1016/j.biopsych.2019.06.013

Kong, R., Li, J., Orban, C., Sabuncu, M. R., Liu, H., Schaefer, A., Sun, N., Zuo, X.-N., Holmes, A. J., Eickhoff, S. B., & Yeo, B. T. T. (2019). Spatial Topography of Individual-Specific Cortical Networks Predicts Human Cognition, Personality, and Emotion. Cerebral Cortex, 29(6), 2533–2551. https://doi.org/10.1093/cercor/bhy123

Kong, R., Yang, Q., Gordon, E., Xue, A., Yan, X., Orban, C., Zuo, X.-N., Spreng, N., Ge, T., Holmes, A., Eickhoff, S., & Yeo, B. T. T. (2021). Individual-Specific Areal-Level Parcellations Improve Functional Connectivity Prediction of Behavior. Cerebral Cortex, 31(10), 4477–4500. https://doi.org/10.1093/cercor/bhab101

Lewis, G. J., Cox, S. R., Booth, T., Muñoz Maniega, S., Royle, N. A., Valdés Hernández, M., Wardlaw, J. M., Bastin, M. E., & Deary, I. J. (2016). Trait conscientiousness and the personality meta-trait stability are associated with regional white matter microstructure. Social Cognitive and Affective Neuroscience, 11(8), 1255–1261. https://doi.org/10.1093/scan/nsw037

Li, J., Kong, R., Liégeois, R., Orban, C., Tan, Y., Sun, N., Holmes, A. J., Sabuncu, M. R., Ge, T., & Yeo, B. T. T. (2019). Global signal regression strengthens association between resting-state functional connectivity and behavior. NeuroImage, 196, 126–141. https://doi.org/https://doi.org/10.1016/j.neuroimage.2019.04.016

Liégeois, R., Li, J., Kong, R., Orban, C., Van De Ville, D., Ge, T., Sabuncu, M. R., & Yeo, B. T. T. (2019). Resting brain dynamics at different timescales capture distinct aspects of human behavior. Nature Communications, 10(1), 2317. https://doi.org/10.1038/s41467-019-10317-7

Liu, X., Lai, H., Li, J., Becker, B., Zhao, Y., Cheng, B., & Wang, S. (2021). Gray matter structures associated with neuroticism: A meta-analysis of whole-brain voxel-based morphometry studies. Human Brain Mapping, 42(9), 2706–2721. https://doi.org/10.1002/hbm.25395

Llera, A., Wolfers, T., Mulders, P., & Beckmann, C. F. (2019). Inter-individual differences in human brain structure and morphology link to variation in demographics and behavior. eLife, 8. https://doi.org/10.7554/elife.44443

Lu, F., Huo, Y., Li, M., Chen, H., Liu, F., Wang, Y., Long, Z., Duan, X., Zhang, J., Zeng, L., & Chen, H. (2014). Relationship between Personality and Gray Matter Volume in Healthy Young Adults: A Voxel-Based Morphometric Study. PLoS ONE, 9(2), e88763. https://doi.org/10.1371/journal.pone.0088763

Lui, S., Zhou, X. J., Sweeney, J. A., & Gong, Q. (2016). Psychoradiology: The Frontier of Neuroimaging in Psychiatry. Radiology, 281(2), 357–372. https://doi.org/10.1148/radiol.2016152149

Mansour, S., Tian, Y., Yeo, B. T. T., Cropley, V., & Zalesky, A. (2021). High-resolution connectomic fingerprints: Mapping neural identity and behavior. NeuroImage, 229, 117695. https://doi.org/10.1016/j.neuroimage.2020.117695

Meng, X., Jiang, R., Lin, D., Bustillo, J., Jones, T., Chen, J., Yu, Q., Du, Y., Zhang, Y., Jiang, T., Sui, J., & Calhoun, V. D. (2017). Predicting individualized clinical measures by a generalized prediction framework and multimodal fusion of MRI data. NeuroImage, 145(Pt B), 218–229. https://doi.org/10.1016/j.neuroimage.2016.05.026

Mill, R. D., Winfield, E. C., Cole, M. W., & Ray, S. (2021). Structural MRI and functional connectivity features predict current clinical status and persistence behavior in prescription opioid users. NeuroImage. Clinical, 30, 102663–102663. https://doi.org/10.1016/j.nicl.2021.102663

Nadeau, C., & Benigo, Y. (2003). Inference for Generalization Error. Machine Learning, 52(3), 239–281. https://doi.org/10.1023/a:1024068626366

Peter, L.-J., Schindler, S., Sander, C., Schmidt, S., Muehlan, H., McLaren, T., Tomczyk, S., Speerforck, S., & Schomerus, G. (2021). Continuum beliefs and mental illness stigma: a systematic review and meta-analysis of correlation and intervention studies. Psychological Medicine, 51(5), 716–726. https://doi.org/10.1017/s0033291721000854

Rasero, J., Sentis, A. I., Yeh, F.-C., & Verstynen, T. (2021). Integrating across neuroimaging modalities boosts prediction accuracy of cognitive ability. PLOS Computational Biology, 17(3), e1008347. https://doi.org/10.1371/journal.pcbi.1008347

Recht, B., Roelofs, R., Schmidt, L., & Shankar, V. (2019, 2019-06-12). Do ImageNet Classifiers Generalize to ImageNet? Proceedings of the 36th International Conference on Machine Learning, Proceedings of Machine Learning Research. https://proceedings.mlr.press/v97/recht19a.html

Rosenberg, M. D., Casey, B. J., & Holmes, A. J. (2018). Prediction complements explanation in understanding the developing brain. Nature Communications, 9(1). https://doi.org/10.1038/s41467-018-02887-9

Rosenberg, M. D., Finn, E. S., Scheinost, D., Papademetris, X., Shen, X., Constable, R. T., & Chun, M. M. (2016). A neuromarker of sustained attention from whole-brain functional connectivity. Nature Neuroscience, 19(1), 165–171. https://doi.org/10.1038/nn.4179

Sabuncu, M. R., & Konukoglu, E. (2015). Clinical Prediction from Structural Brain MRI Scans: A Large-Scale Empirical Study. Neuroinformatics, 13(1), 31–46. https://doi.org/10.1007/s12021-014-9238-1

Schaefer, A., Kong, R., Gordon, E. M., Laumann, T. O., Zuo, X.-N., Holmes, A. J., Eickhoff, S. B., & Yeo, B. T. T. (2018). Local-Global Parcellation of the Human Cerebral Cortex from Intrinsic Functional Connectivity MRI. Cerebral Cortex, 28(9), 3095–3114. https://doi.org/10.1093/cercor/bhx179

Smith, S. M., Jenkinson, M., Johansen-Berg, H., Rueckert, D., Nichols, T. E., Mackay, C. E., Watkins, K. E., Ciccarelli, O., Cader, M. Z., Matthews, P. M., & Behrens, T. E. J. (2006). Tract-based spatial statistics: Voxelwise analysis of multi-subject diffusion data. NeuroImage, 31(4), 1487–1505. https://doi.org/10.1016/j.neuroimage.2006.02.024

Smith, S. M., Nichols, T. E., Vidaurre, D., Winkler, A. M., Behrens, T. E. J., Glasser, M. F., Ugurbil, K., Barch, D. M., Van Essen, D. C., & Miller, K. L. (2015). A positive-negative mode of population covariation links brain connectivity, demographics and behavior. Nature Neuroscience, 18(11), 1565–1567. https://doi.org/10.1038/nn.4125

Sripada, C., Rutherford, S., Angstadt, M., Thompson, W. K., Luciana, M., Weigard, A., Hyde, L. H., & Heitzeg, M. (2020). Prediction of neurocognition in youth from resting state fMRI. Molecular Psychiatry, 25(12), 3413–3421. https://doi.org/10.1038/s41380-019-0481-6

Sui, J., Jiang, R., Bustillo, J., & Calhoun, V. (2020). Neuroimaging-based Individualized Prediction of Cognition and Behavior for Mental Disorders and Health: Methods and Promises. Biological Psychiatry, 88(11), 818–828. https://doi.org/https://doi.org/10.1016/j.biopsych.2020.02.016

Thompson, W. H., Wright, J., Bissett, P. G., & Poldrack, R. A. (2020). Dataset decay and the problem of sequential analyses on open datasets. eLife, 9. https://doi.org/10.7554/elife.53498

Tournier, J. D., Smith, R., Raffelt, D., Tabbara, R., Dhollander, T., Pietsch, M., Christiaens, D., Jeurissen, B., Yeh, C.-H., & Connelly, A. (2019). MRtrix3: A fast, flexible and open software framework for medical image processing and visualisation. NeuroImage, 202, 116137. https://doi.org/10.1016/j.neuroimage.2019.116137

Uher, J. (2015). Developing “Personality” Taxonomies: Metatheoretical and Methodological Rationales Underlying Selection Approaches, Methods of Data Generation and Reduction Principles. Integrative Psychological and Behavioral Science, 49(4), 531–589. https://doi.org/10.1007/s12124-014-9280-4

Van Essen, D. C., Smith, S. M., Barch, D. M., Behrens, T. E. J., Yacoub, E., & Ugurbil, K. (2013). The WU-Minn Human Connectome Project: An overview. NeuroImage, 80, 62–79. https://doi.org/https://doi.org/10.1016/j.neuroimage.2013.05.041

Xia, C. H., Ma, Z., Ciric, R., Gu, S., Betzel, R. F., Kaczkurkin, A. N., Calkins, M. E., Cook, P. A., García de la Garza, A., Vandekar, S. N., Cui, Z., Moore, T. M., Roalf, D. R., Ruparel, K., Wolf, D. H., Davatzikos, C., Gur, R. C., Gur, R. E., Shinohara, R. T., Bassett, D. S., & Satterthwaite, T. D. (2018). Linked dimensions of psychopathology and connectivity in functional brain networks. Nature Communications, 9(1), 3003. https://doi.org/10.1038/s41467-018-05317-y

Xiao, Y., Lin, Y., Ma, J., Qian, J., Ke, Z., Li, L., Yi, Y., Zhang, J., & Dai, Z. (2021). Predicting visual working memory with multimodal magnetic resonance imaging. Human Brain Mapping, 42(5), 1446–1462. https://doi.org/10.1002/hbm.25305

Zatorre, R. J., Fields, R. D., & Johansen-Berg, H. (2012). Plasticity in gray and white: neuroimaging changes in brain structure during learning. Nature Neuroscience, 15(4), 528–536. https://doi.org/10.1038/nn.3045

