## supplemental_material for "Comparison of individualized behavioral predictions across anatomical, diffusion and functional connectivity MRI"

Table S1. Behavioral measures for HCP

|  | **Description** | **HCP field** |
| --- | --- | --- |
| 1 | Visual Episodic Memory | PicSeq_Unadj |
| 2 | Cognitive Flexibility (DCCS) | CardSort_Unadj |
| 3 | Inhibition (Flanker Task) | Flanker_Unadj |
| 4 | Fluid Intelligence (PMAT) | PMAT24_A_CR |
| 5 | Vocabulary (Pronunciation) | ReadEng_Unadj |
| 6 | Vocabulary (Picture Matching) | PicVocab_Unadj |
| 7 | Processing Speed | ProcSpeed_Unadj |
| 8 | Delay Discounting | DDic_AUC_40K |
| 9 | Spatial Orientation | VSPLOT_TC |
| 10 | Sustained Attention – Sens. | SCPT_SEN |
| 11 | Sustained Attention – Spec. | SCPT_SPEC |
| 12 | Verbal Episodic Memory | IWRD_TOT |
| 13 | Working Memory (List Sorting) | ListSort_Unadj |
| 14 | Cognitive Status (MMSE) | MMSE_Score |
| 15 | Sleep Quality (PSQI) | PSQI_Score |
| 16 | Walking Endurance | Endurance_Unadj |
| 17 | Walking Speed | GaitSpeed_Unadj |
| 18 | Manual Dexterity | Dexterity_Unadj |
| 19 | Grip Strength | Strength_Unadj |
| 20 | Odor Identification | Odor_Unadj |
| 21 | Pain Interference Survey | PainInterf_Tscore |
| 22 | Taste Intensity | Taste_Unadj |
| 23 | Contrast Sensitivity | Mars_Final |
| 24 | Emotional Face Matching | Emotion_Task_Face_Acc |
| 25 | Arithmetic | Language_Task_Math_Avg_Difficulty_Level |
| 26 | Story Comprehension | Language_Task_Story_Avg_Difficulty_Level |
| 27 | Relational Processing | Relational_Task_Acc |
| 28 | Social Cognition – Random | Social_Task_Perc_Random |
| 29 | Social Cognition – Interaction | Social_Task_Perc_TOM |
| 30 | Working Memory (N-back) | WM_Task_Acc |
| 31 | Agreeableness (NEO) | NEOFAC_A |
| 32 | Openness (NEO) | NEOFAC_O |
| 33 | Conscientiousness (NEO) | NEOFAC_C |
| 34 | Neuroticism (NEO) | NEOFAC_N |
| 35 | Extraversion (NEO) | NEOFAC_E |
| 36 | Emot. Recog. – Total | ER40_CR |
| 37 | Emot. Recog. – Angry | ER40ANG |
| 38 | Emot. Recog. – Fear | ER40FEAR |
| 39 | Emot. Recog. – Happy | ER40HAP |
| 40 | Emot. Recog. - Neutral | ER40NOE |
| 41 | Emot. Recog. – Sad | ER40SAD |
| 42 | Anger – Affect | AngAffect_Unadj |
| 43 | Anger – Hostility | AngHostil_Unadj |
| 44 | Anger – Aggression | AngAggr_Unadj |
| 45 | Fear – Affect | FearAffect_Unadj |
| 46 | Fear – Somatic Arousal | FearSomat_Unadj |
| 47 | Sadness | Sadness_Unadj |
| 48 | Life Satisfaction | LifeSatisf_Unadj |
| 49 | Meaning & Purpose | MeanPurp_Unadj |
| 50 | Positive Affect | PosAffect_Unadj |
| 51 | Friendship | Friendship_Unadj |
| 52 | Loneliness | Loneliness_Unadj |
| 53 | Perceived Hostility | PercHostil_Unadj |
| 54 | Perceived Rejection | PercReject_Unadj |
| 55 | Emotional Support | EmotSupp_Unadj |
| 56 | Instrument Support | InstruSupp_Unadj |
| 57 | Perceived Stress | PercStress_Unadj |
| 58 | Self-Efficacy | SelfEff_Unadj |

Table S2. Behavioral measures for ABCD

|  | **Description** | **ABCD field** | **ABCD file** |
| --- | --- | --- | --- |
| 1 | Anxious Depressed | cbcl_scr_syn_anxdep_r | abcd_cbcls01.txt |
| 2 | Withdrawn Depressed | cbcl_scr_syn_withdep_r | abcd_cbcls01.txt |
| 3 | Somatic Complaints | cbcl_scr_syn_somatic_r | abcd_cbcls01.txt |
| 4 | Social Problems | cbcl_scr_syn_social_r | abcd_cbcls01.txt |
| 5 | Thought Problems | cbcl_scr_syn_thought_r | abcd_cbcls01.txt |
| 6 | Attention Problems | cbcl_scr_syn_attention_r | abcd_cbcls01.txt |
| 7 | Rule-breaking Behavior | cbcl_scr_syn_rulebreak_r | abcd_cbcls01.txt |
| 8 | Aggressive Behavior | cbcl_scr_syn_aggressive_r | abcd_cbcls01.txt |
| 9 | Vocabulary | nihtbx_picvocab_uncorrected | abcd_tbss01.txt |
| 10 | Attention | nihtbx_flanker_uncorrected | abcd_tbss01.txt |
| 11 | Working Memory | nihtbx_list_uncorrected | abcd_tbss01.txt |
| 12 | Executive Function | nihtbx_cardsort_uncorrected | abcd_tbss01.txt |
| 13 | Processing Speed | nihtbx_pattern_uncorrected | abcd_tbss01.txt |
| 14 | Episodic Memory | nihtbx_picture_uncorrected | abcd_tbss01.txt |
| 15 | Reading | nihtbx_reading_uncorrected | abcd_tbss01.txt |
| 16 | Fluid Cognition | nihtbx_fluidcomp_uncorrected | abcd_tbss01.txt |
| 17 | Crystallized Cognition | nihtbx_cryst_uncorrected | abcd_tbss01.txt |
| 18 | Overall Cognition | nihtbx_totalcomp_uncorrected | abcd_tbss01.txt |
| 19 | Negative Urgency | upps_y_ss_negative_urgency | abcd_mhy02.txt |
| 20 | Lack of Planning | upps_y_ss_lack_of_planning | abcd_mhy02.txt |
| 21 | Sensation Seeking | upps_y_ss_sensation_seeking | abcd_mhy02.txt |
| 22 | Positive Urgency | upps_y_ss_positive_urgency | abcd_mhy02.txt |
| 23 | Lack Perseverance | upps_y_lack_of_perseverance | abcd_mhy02.txt |
| 24 | Behavioral Inhibition | bis_y_ss_bis_sum | abcd_mhy02.txt |
| 25 | Reward Responsiveness | bis_y_ss_bas_rr | abcd_mhy02.txt |
| 26 | Drive | bis_y_ss_bas_drive | abcd_mhy02.txt |
| 27 | Fun Seeking | bis_y_ss_bas_fs | abcd_mhy02.txt |
| 28 | Total Psychosis Symptoms | pps_y_ss_number | abcd_mhy02.txt |
| 29 | Psychosis Severity | pps_y_ss_severity_score | abcd_mhy02.txt |
| 30 | Mania | pgbi_p_ss_score | abcd_mhp02.txt |
| 31 | Short Delay Recall | pea_ravlt_sd_trial_vi_tc | abcd_ps01.txt |
| 32 | Long Delay Recall | pea_ravlt_ld_trial_vii_tc | abcd_ps01.txt |
| 33 | Fluid Intelligence | pea_wiscv_trs | abcd_ps01.txt |
| 34 | Visuospatial Accuracy | lmt_scr_perc_correct | lmtp201.txt |
| 35 | Visuospatial Reaction Time | lmt_scr_rt_correct | lmtp201.txt |
| 36 | Visuospatial Efficiency | lmt_scr_efficiency | lmtp201.txt |

Table S3. Top 10 loadings (absolute) for HCP behavioral factors

|  | **Cognition** | | **Dissatisfaction** | | **Emotion** | |
| --- | --- | --- | --- | --- | --- | --- |
| 1 | Working Memory (N-back) | 0.3024 | Sadness | 0.2740 | Emot. Recog. - Total | 0.4951 |
| 2 | Vocabulary (Pronunciation) | 0.2964 | Perceived Stress | 0.2738 | Emot Recog - Fear | 0.3384 |
| 3 | Story Comprehension | 0.2900 | Loneliness | 0.2691 | Emot. Recog - Sad | 0.3293 |
| 4 | Vocabulary (Picture Matching) | 0.2803 | Neuroticism (NEO) | 0.2599 | Grip Strength | 0.3067 |
| 5 | Fluid Intelligence (PMAT) | 0.2730 | Fear - Affect | 0.2423 | Emot. Recog. - Anger | 0.2720 |
| 6 | Relational Processing | 0.2627 | Anger - Affect | 0.2363 | Agreeableness (NEO) | 0.2485 |
| 7 | Spatial Orientation | 0.2459 | Perceived Rejection | 0.2353 | Anger - Aggression | -0.2348 |
| 8 | Working Memory (List Sorting) | 0.2325 | Positive Affect | -0.2315 | Manual Dexterity | 0.1636 |
| 9 | Walking Endurance | 0.2258 | Life Satisfaction | -0.2255 | Perceived Hostility | -0.1629 |
| 10 | Cognitive Flexibility (DCCS) | 0.2201 | Emotional Support | -0.2226 | Verbal Episodic Memory | 0.1596 |

Table S4. Top 10 loadings (absolute) for ABCD behavioral factors

|  | **Cognition** | | **Personality** | | **Mental health** | |
| --- | --- | --- | --- | --- | --- | --- |
| 1 | Overall Cognition | 0.3760 | Fun seeking | 0.3883 | Aggressive Behavior | 0.3652 |
| 2 | Fluid Cognition | 0.3442 | Total Psychosis Symptoms | 0.3455 | Thought Problems | 0.3584 |
| 3 | Crystalized Cognition | 0.2975 | Reward Responsiveness | 0.3450 | Social Problems | 0.3568 |
| 4 | Reading | 0.2619 | Drive | 0.3415 | Anxious Depressed | 0.3408 |
| 5 | Vocabulary | 0.2603 | Psychosis Severity | 0.3365 | Attention Problems | 0.3347 |
| 6 | Working Memory | 0.2585 | Positive Urgency | 0.3246 | Withdrawn Depressed | 0.3219 |
| 7 | Executive Function | 0.2468 | Negative Urgency | 0.3178 | Mania | 0.3079 |
| 8 | Attention | 0.2282 | Behavioral inhibition | 0.2798 | Rule-breaking behavior | 0.3072 |
| 9 | Long Delay Recall | 0.2243 | Sensation Seeking | 0.2299 | Somatic Complaints | 0.2611 |
| 10 | Short Delay Recall | 0.2229 | Visuospatial Reaction Time | -0.0947 | Lack perseverance | 0.0923 |

Table S5. Site clusters for ABCD

| **ABCD Site** | **Make** | **Model** | **N** | **Site-cluster** |
| --- | --- | --- | --- | --- |
| 4 | GE | Discovery MR750 | 89 | A |
| 10 | GE | Discovery MR750 | 102 | A |
| 8 | GE | Discovery MR750 | 49 | B |
| 13 | GE | Discovery MR750 | 70 | B |
| 18 | GE | Discovery MR750 | 63 | B |
| 22 | GE | Discovery MR750 | 1 | B |
| 3 | Siemens | Prisma | 141 | C |
| 11 | Siemens | Prisma | 67 | C |
| 16 | Siemens | Prisma | 320 | D |
| 14 | Siemens | Prisma/Prisma fit | 163 | E |
| 7 | Siemens | Prisma fit | 62 | F |
| 20 | Siemens | Prisma/Prisma fit | 95 | F |
| 5 | Siemens | Prisma fit | 67 | G |
| 21 | Siemens | Prisma fit/Prisma | 81 | G |
| 2 | Siemens | Prisma fit | 123 | H |
| 15 | Siemens | Prisma fit | 29 | H |
| 6 | Siemens | Prisma fit | 141 | I |
| 9 | Siemens | Prisma fit | 62 | J |
| 12 | Siemens | Prisma fit | 98 | J |

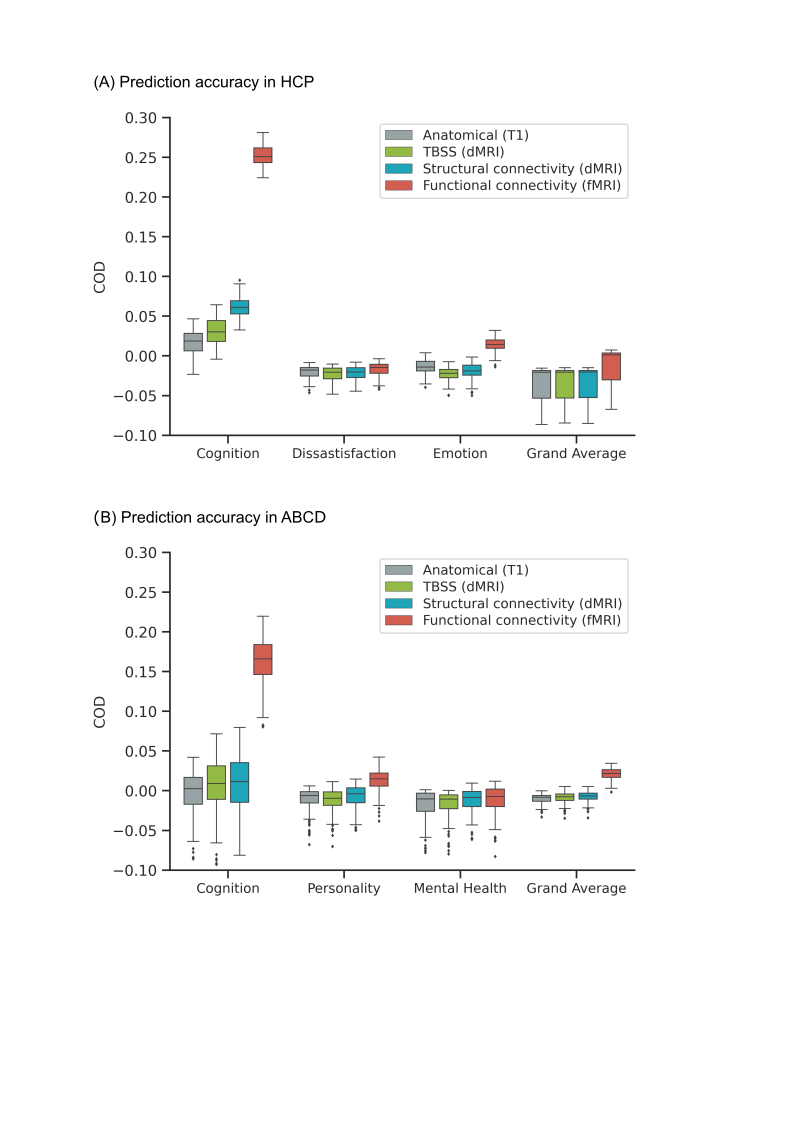

**Figure S1.** Functional connectivity (FC) outperforms other modalities for kernel ridge regression (KRR). Figure is the same as Figure 1 except that COD is shown instead of Pearson’s correlation. (A) Prediction performance (COD) of KRR averaged across single-feature-type predictive models within each modality (anatomical, TBSS, structural connectivity, functional connectivity) in the HCP dataset. Results are shown for the three behavioral components and “grand average” obtained by averaging prediction performance across 58 behavioral measures. Each boxplot shows the distribution of performance over 60 repetitions of the nested cross-validation procedure. (B) Prediction performance (COD) of KRR averaged across single-feature-type predictive models within each modality (anatomical, TBSS, structural connectivity, functional connectivity) in the ABCD dataset. Results are shown for the three behavioral components and “grand average” obtained by averaging prediction performance across 36 behavioral measures. Each boxplot shows the distribution of performance over 120 repetitions of the nested cross-validation procedure.

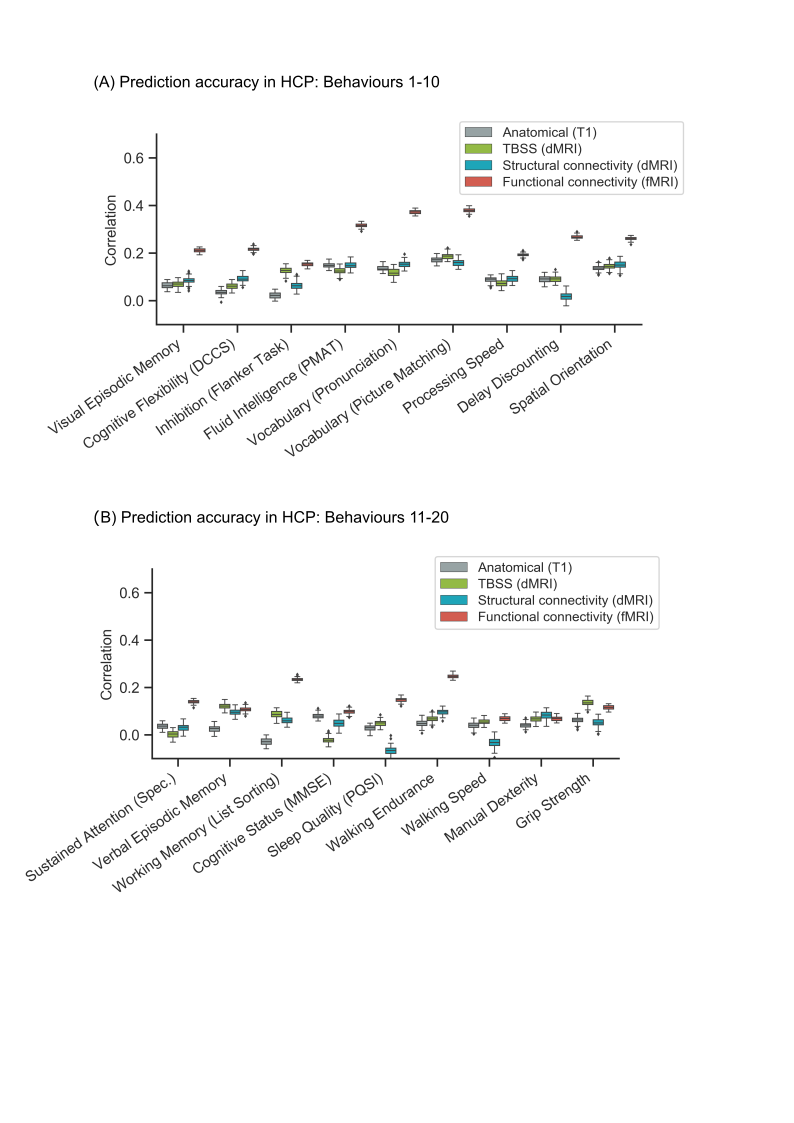

**Figure S2.** Prediction performance (Pearson’s correlation) of kernel ridge regression (KRR) for individual behavioural measures in the HCP dataset. (A) Prediction performance (Pearson’s correlation) of KRR averaged across single-feature-type predictive models within each modality (anatomical, TBSS, structural connectivity, functional connectivity) in the HCP dataset. Results are shown for behaviors 1-10 of Table S1. Each boxplot shows the distribution of performance over 60 repetitions of the nested cross-validation procedure. (B) Prediction performance (Pearson’s correlation) of KRR averaged across single-feature-type predictive models within each modality (anatomical, TBSS, structural connectivity, functional connectivity) in the HCP dataset. Results are shown for behaviors 11-21 of Table S1. Each boxplot shows the distribution of performance over 60 repetitions of the nested cross-validation procedure.

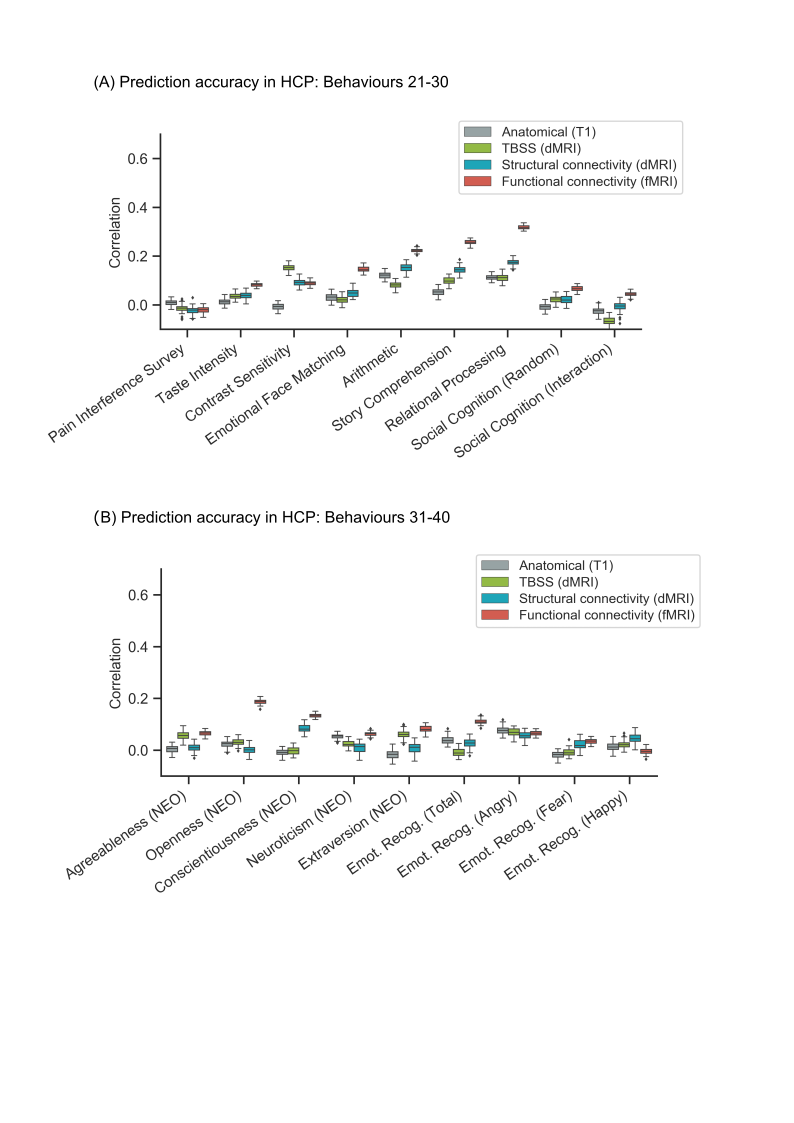

**Figure S3.** Prediction performance (Pearson’s correlation) of kernel ridge regression (KRR) for individual behavioural measures in the HCP dataset. (A) Prediction performance (Pearson’s correlation) of KRR averaged across single-feature-type predictive models within each modality (anatomical, TBSS, structural connectivity, functional connectivity) in the HCP dataset. Results are shown for behaviors 21-30 of Table S1. Each boxplot shows the distribution of performance over 60 repetitions of the nested cross-validation procedure. (B) Prediction performance (Pearson’s correlation) of KRR averaged across single-feature-type predictive models within each modality (anatomical, TBSS, structural connectivity, functional connectivity) in the HCP dataset. Results are shown for behaviors 31-40 of Table S1. Each boxplot shows the distribution of performance over 60 repetitions of the nested cross-validation procedure.

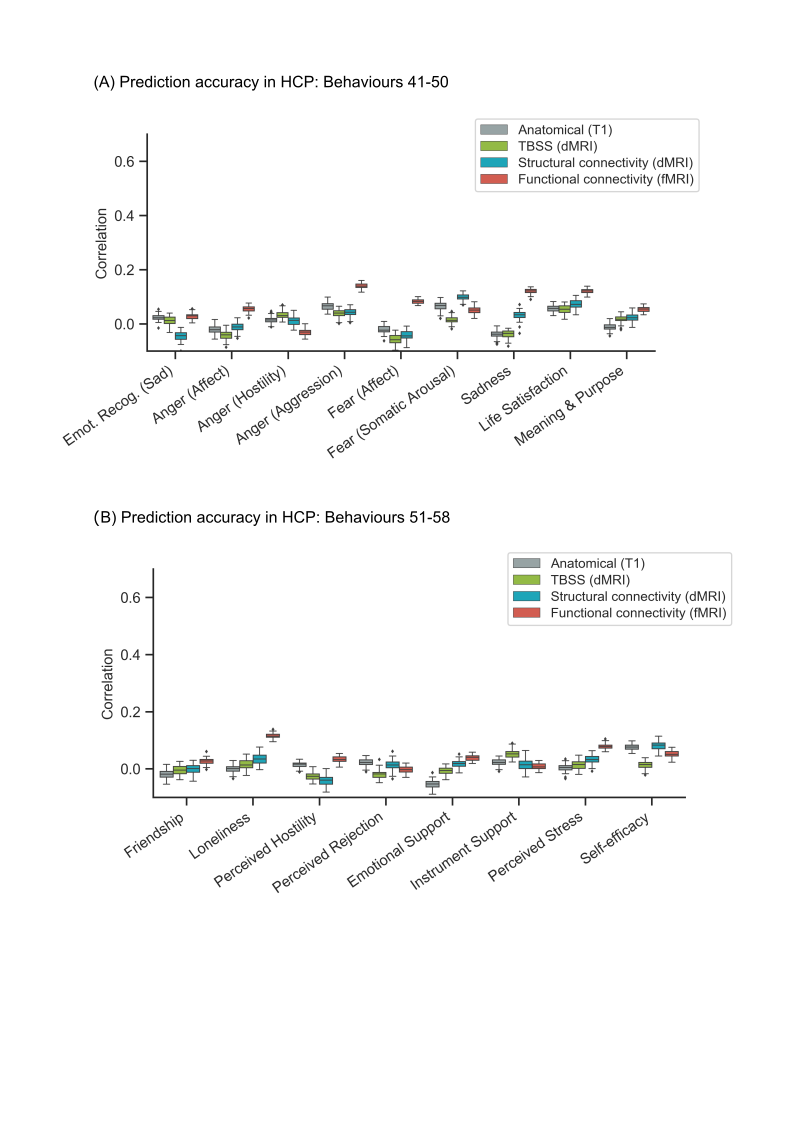

**Figure S4.** (Prediction performance (Pearson’s correlation) of kernel ridge regression (KRR) for individual behavioural measures in the HCP dataset. (A) Prediction performance (Pearson’s correlation) of KRR averaged across single-feature-type predictive models within each modality (anatomical, TBSS, structural connectivity, functional connectivity) in the HCP dataset. Results are shown for behaviors 41-50 of Table S1. Each boxplot shows the distribution of performance over 60 repetitions of the nested cross-validation procedure. (B) Prediction performance (Pearson’s correlation) of KRR averaged across single-feature-type predictive models within each modality (anatomical, TBSS, structural connectivity, functional connectivity) in the HCP dataset. Results are shown for behaviors 51-58 of Table S1. Each boxplot shows the distribution of performance over 60 repetitions of the nested cross-validation procedure.

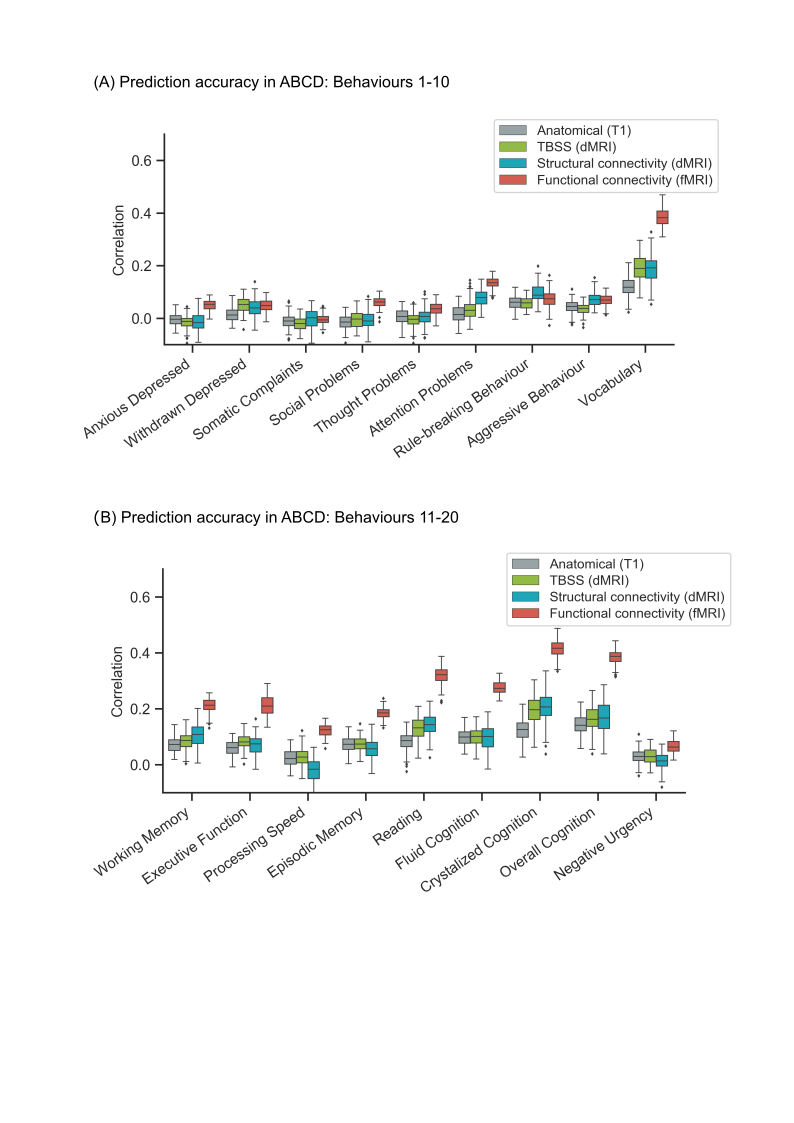

**Figure S5.** Prediction performance (Pearson’s correlation) of kernel ridge regression (KRR) for individual behavioural measures in the ABCD dataset. (A) Prediction performance (Pearson’s correlation) of KRR averaged across single-feature-type predictive models within each modality (anatomical, TBSS, structural connectivity, functional connectivity) in the ABCD dataset. Results are shown for behaviors 1-10 of Table S2. Each boxplot shows the distribution of performance over 120 repetitions of the nested cross-validation procedure. (B) Prediction performance (Pearson’s correlation) of KRR averaged across single-feature-type predictive models within each modality (anatomical, TBSS, structural connectivity, functional connectivity) in the ABCD dataset. Results are shown for behaviors 11-20 of Table S2. Each boxplot shows the distribution of performance over 120 repetitions of the nested cross-validation procedure.

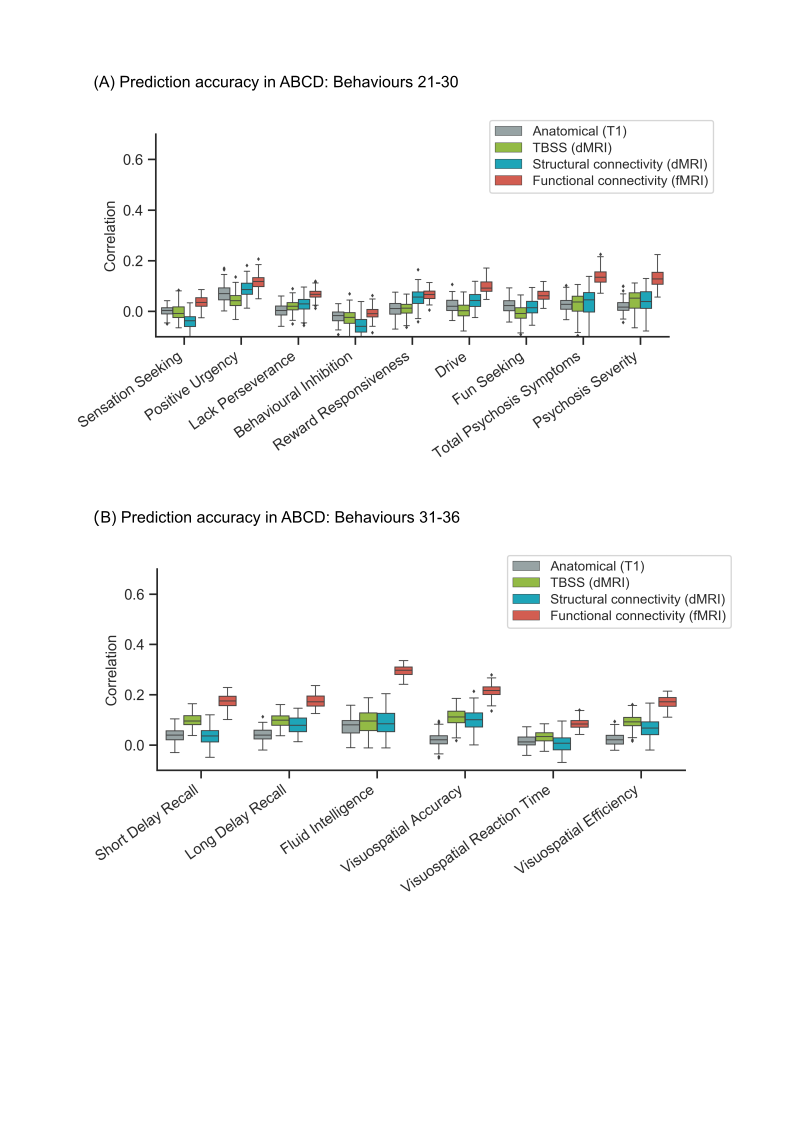

**Figure S6.** Prediction performance (Pearson’s correlation) of kernel ridge regression (KRR) for individual behavioural measures in the ABCD dataset. (A) Prediction performance (Pearson’s correlation) of KRR averaged across single-feature-type predictive models within each modality (anatomical, TBSS, structural connectivity, functional connectivity) in the ABCD dataset. Results are shown for behaviors 21-30 of Table S2. Each boxplot shows the distribution of performance over 120 repetitions of the nested cross-validation procedure. (B) Prediction performance (Pearson’s correlation) of KRR averaged across single-feature-type predictive models within each modality (anatomical, TBSS, structural connectivity, functional connectivity) in the ABCD dataset. Results are shown for behaviors 31-36 of Table S2. Each boxplot shows the distribution of performance over 120 repetitions of the nested cross-validation procedure.

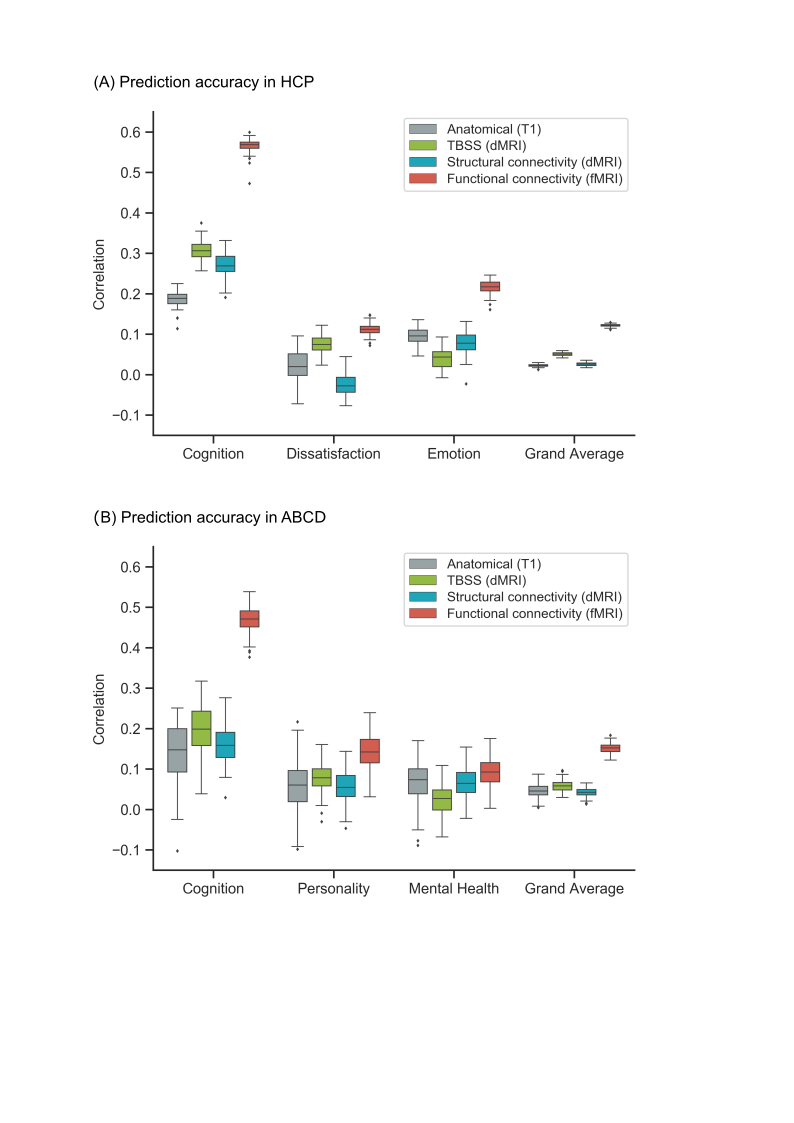

**Figure S7.** Functional connectivity (FC) outperforms other modalities for linear ridge regression (LRR). Figure is the same as Figure 4 except that LRR was utilized instead of kernel ridge regression. (A) Prediction performance (Pearson’s correlation) of LRR for the best performing feature-type within each modality in the HCP dataset. For the cognition component, the best features were cortical area, TBSS OD, SC FA and language FC. For the dissatisfaction component, the best features were cortical thickness, TBSS FA, SC stream count and working memory FC. For the emotion component, the best features were cortical volume, TBSS AD, SC FA and social cognition FC. For the grand average, the best features were cortical volume, TBSS AD, SC AD and language FC. (B) Prediction performance (Pearson’s correlation) of LRR for the best performing feature-type within each modality in the ABCD dataset. For the cognition component, the best features were cortical area, TBSS OD, SC ICVF and N-back FC. For the personality component, the best features were cortical volume, TBSS OD, SC MD and MID FC. For the mental health component, the best features were cortical area, TBSS OD, SC MD and resting FC. For the grand average, the best features were cortical area, TBSS OD, SC ICVF and N-back FC.

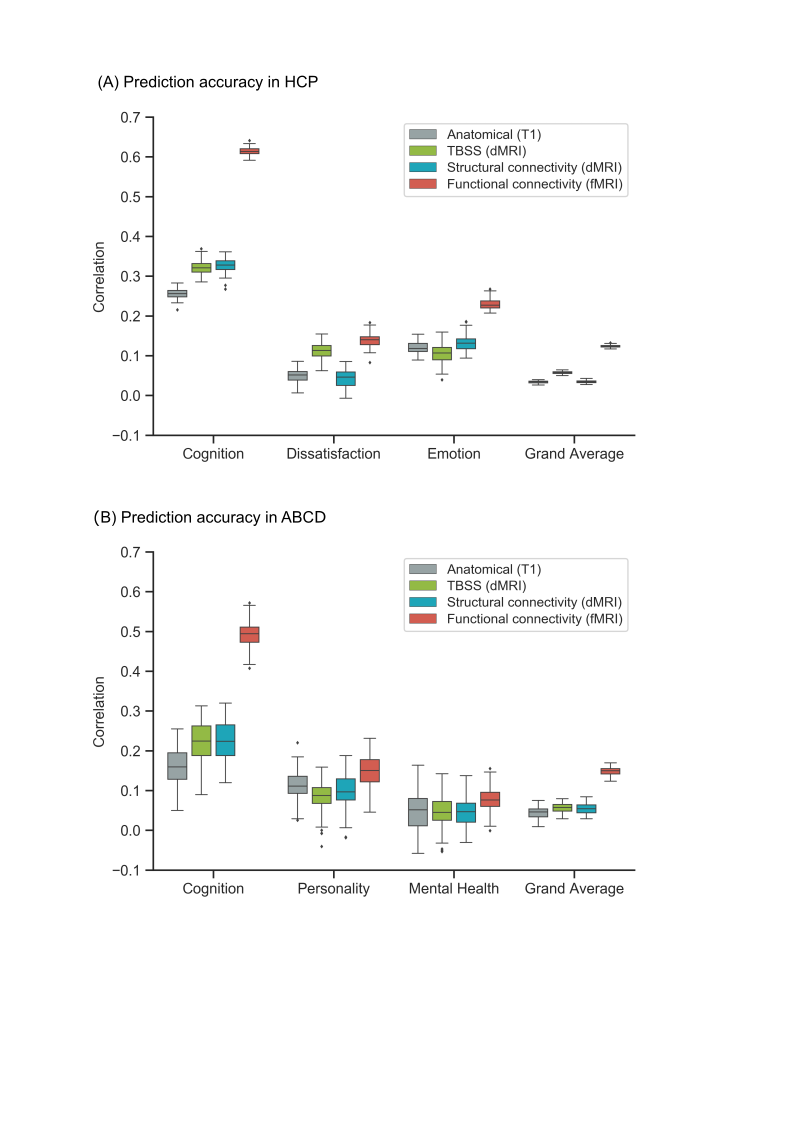

**Fig S8.** Functional connectivity (FC) outperforms other modalities for elastic net. Figure is the same as Figure 4 except that elastic net was utilized instead of kernel ridge regression. (A) Prediction performance (Pearson’s correlation) of elastic net for the best performing feature-type within each modality in the HCP dataset. For the cognition component, the best features were cortical area, TBSS FA, SC FA and language FC. For the dissatisfaction component, the best features were cortical thickness, TBSS AD, SC stream length and working memory FC. For the emotion component, the best features were cortical volume, TBSS OD, SC FA and language FC. For the grand average, the best features were cortical thickness, TBSS FA, SC AD and language FC. (B) Prediction performance (Pearson’s correlation) of elastic net for the best performing feature-type within each modality in the ABCD dataset. For the cognition component, the best features were cortical thickness, TBSS OD, SC ICVF and N-back FC. For the personality component, the best features were cortical thickness, TBSS OD, SC OD and MID FC. For the mental health component, the best features were cortical area, TBSS OD, SC OD and SST FC. For the grand average, the best features were cortical area, TBSS OD, SC ICVF and N-back FC.

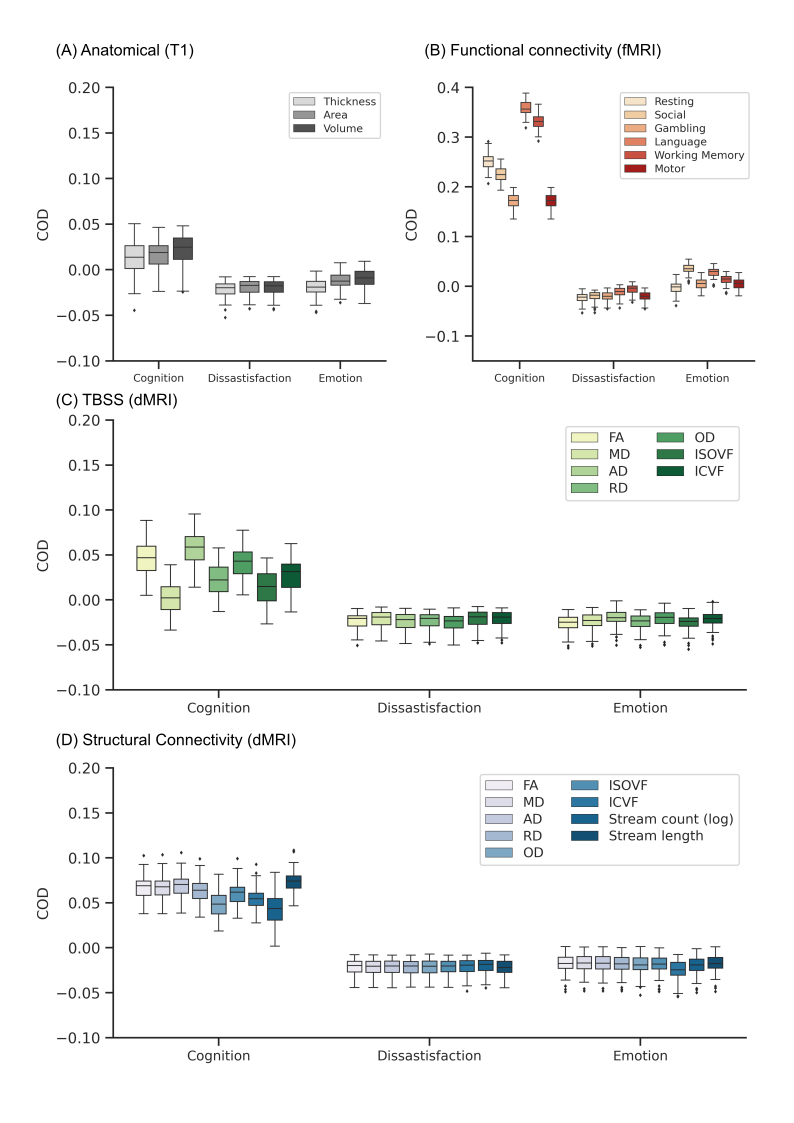

**Figure S9**. Prediction performance (COD) of kernel ridge regression (KRR) for each single-feature-type in the HCP dataset. Figure is the same as Figure 5, except that COD was shown instead of Pearson’s correlation. Results are shown separately for (A) anatomical features, (B) FC, (C) TBSS and (D) structural connectivity.

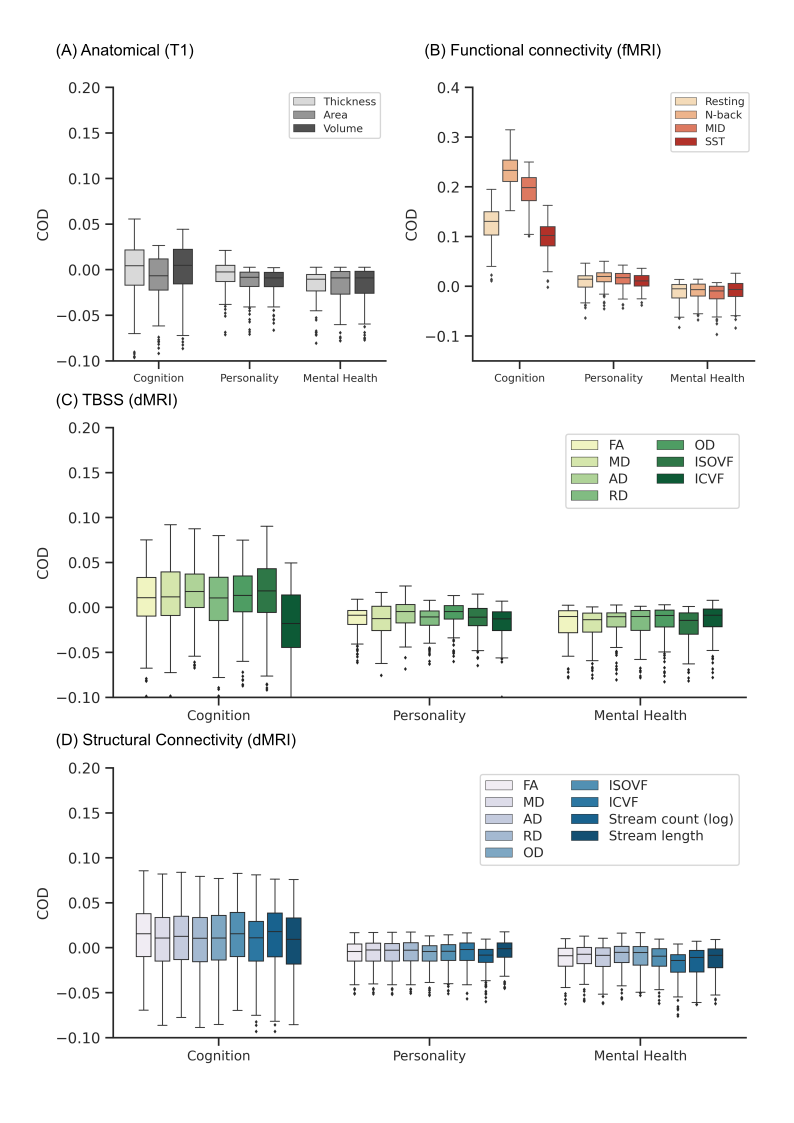

**Figure S10**. Prediction performance (COD) of kernel ridge regression (KRR) for each single-feature-type in the ABCD dataset. Figure is the same as Figure 6, except that COD was shown instead of Pearson’s correlation. Results are shown separately for (A) anatomical features, (B) FC, (C) TBSS and (D) structural connectivity.

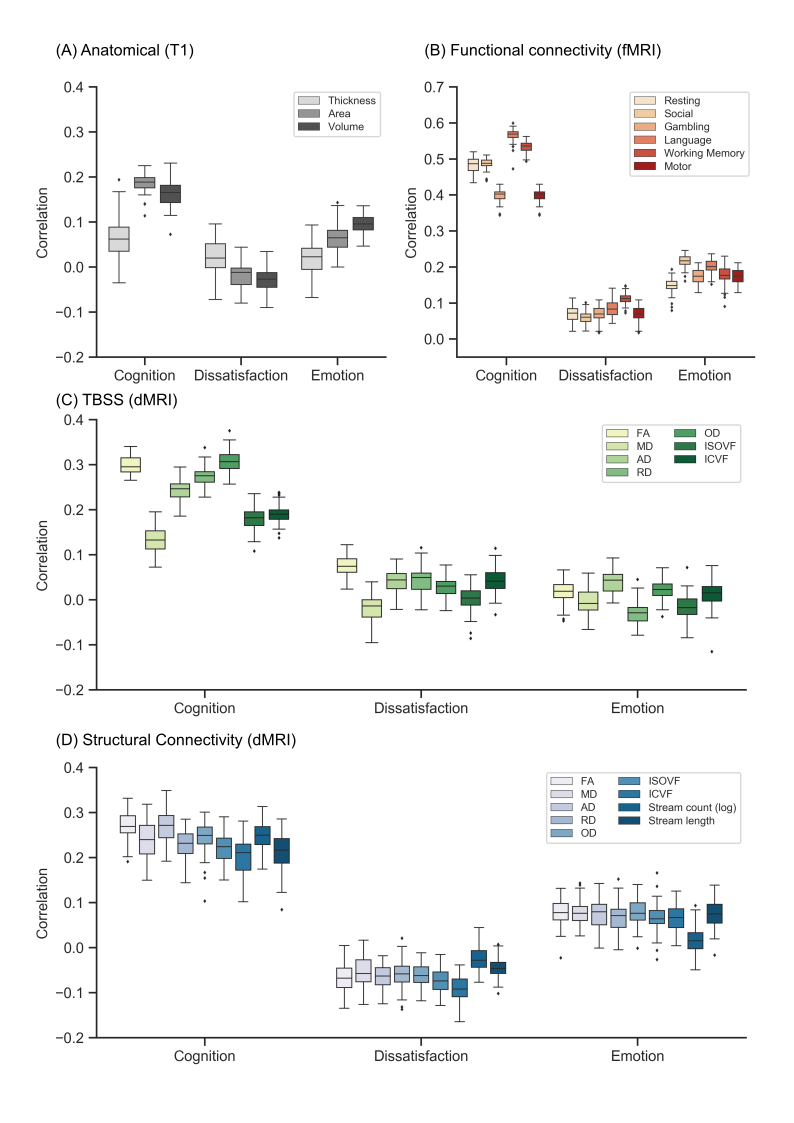

**Figure S11**. Prediction performance (Pearson’s correlation) of linear ridge regression (LRR) for each single-feature-type in the HCP dataset. Figure is the same as Figure 5 except that LRR was utilized instead of kernel ridge regression. Results are shown separately for (A) anatomical features, (B) FC, (C) TBSS and (D) structural connectivity.

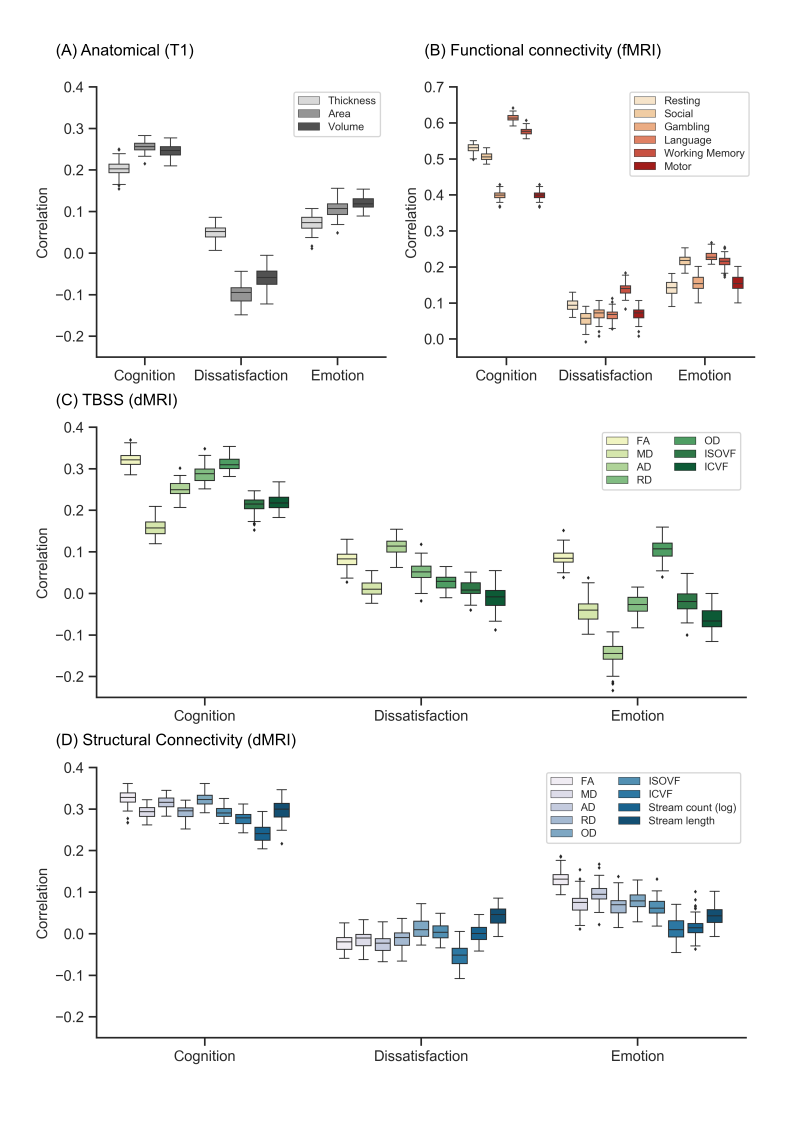

**Figure S12**. Prediction performance (Pearson’s correlation) of elastic net for each single-feature-type in the HCP dataset. Figure is the same as Figure 5 except that elastic net was utilized instead of kernel ridge regression. Results are shown separately for (A) anatomical features, (B) FC, (C) TBSS and (D) structural connectivity.

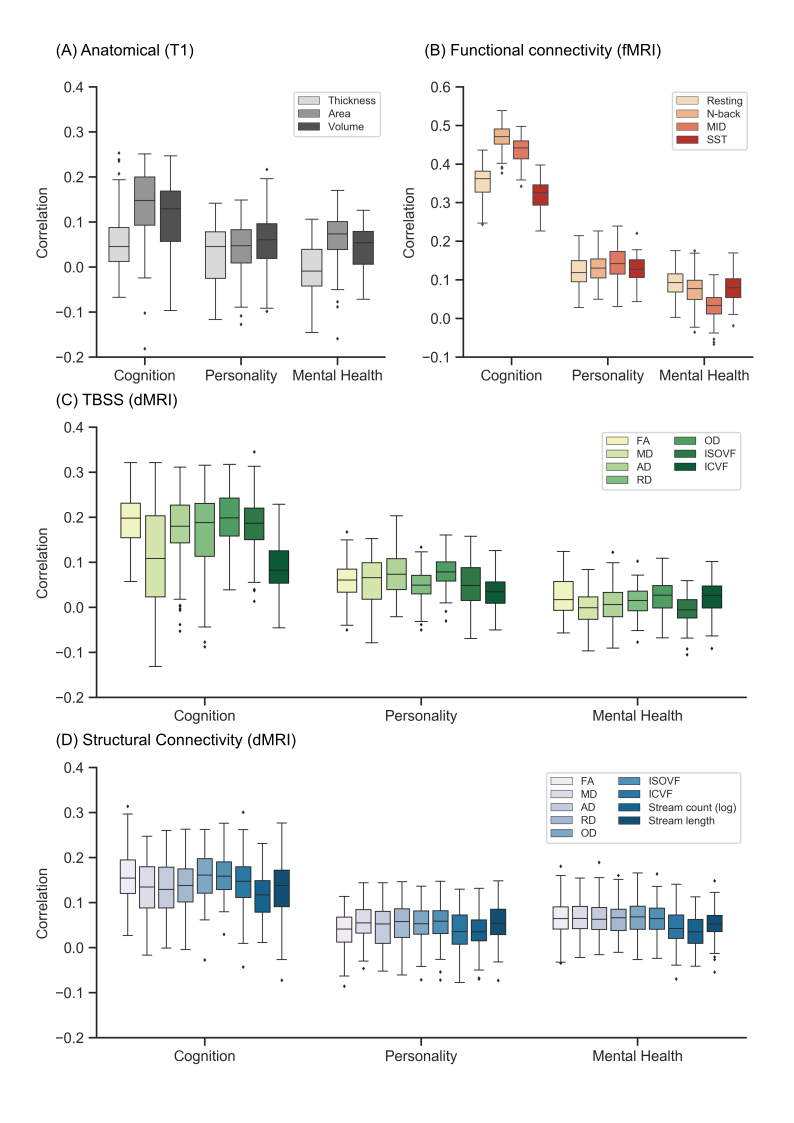

**Figure S13**. Prediction performance (Pearson’s correlation) of linear ridge regression (LRR) for each single-feature-type in the ABCD dataset. Figure is the same as Figure 6 except that LRR was utilized instead of kernel ridge regression. Results are shown separately for (A) anatomical features, (B) FC, (C) TBSS and (D) structural connectivity.

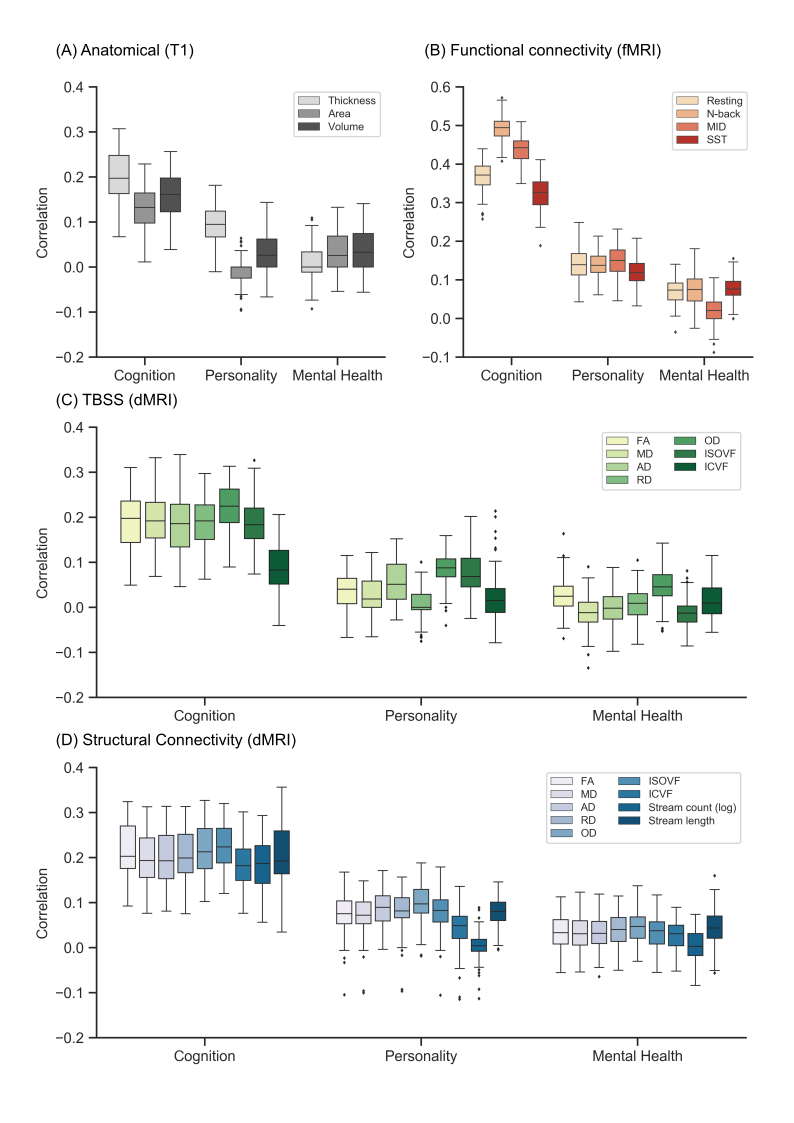

**Figure S14**. Prediction performance (Pearson’s correlation) of elastic net for each single-feature-type in the ABCD dataset. Figure is the same as Figure 6 except that elastic net was utilized instead of kernel ridge regression. Results are shown separately for (A) anatomical features, (B) FC, (C) TBSS and (D) structural connectivity.

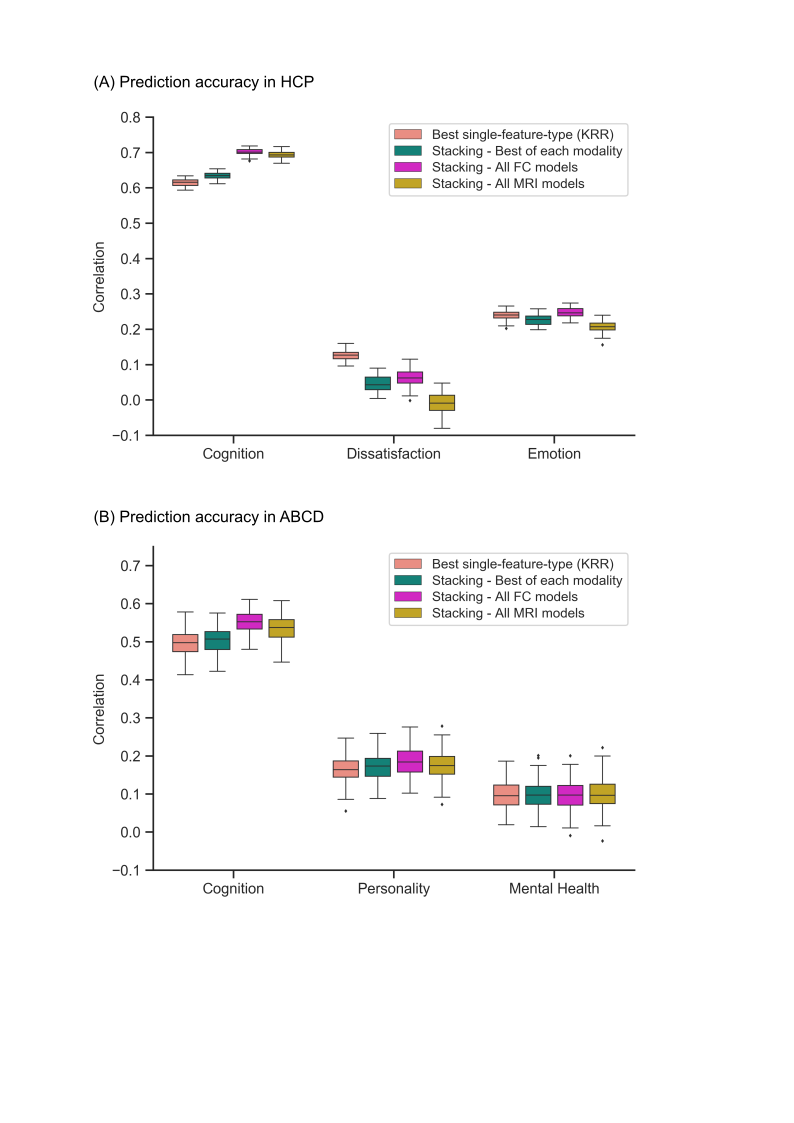

**Figure S15.** Combining resting and task FC was as good as combining across all modalities, or combining the best single-feature-type models of each modality. Figure is the same as Figure 7, except that the results of multi-KRR are replaced with the results of stacking the best single-feature-type models (which can be found in Figure 4). (A) Prediction performance (Pearson’s correlation) from combining various MRI features and modalities in the HCP dataset. We considered stacking the best single-feature-type model of each modality, all FC models, and all single-feature-type models across all modalities. For comparison, the best single-feature-type from KRR is shown. Each boxplot shows the distribution over 60 repetitions of the nested cross-validation procedure. (B) Prediction performance (Pearson’s correlation) from combining various MRI features and modalities in the ABCD dataset. We considered stacking the best single-feature-type model of each modality, all FC models, and all single-feature-type models across all modalities. For comparison, the best single-feature-type from KRR is shown. Each boxplot shows the distribution over 120 repetitions of the nested cross-validation procedure.

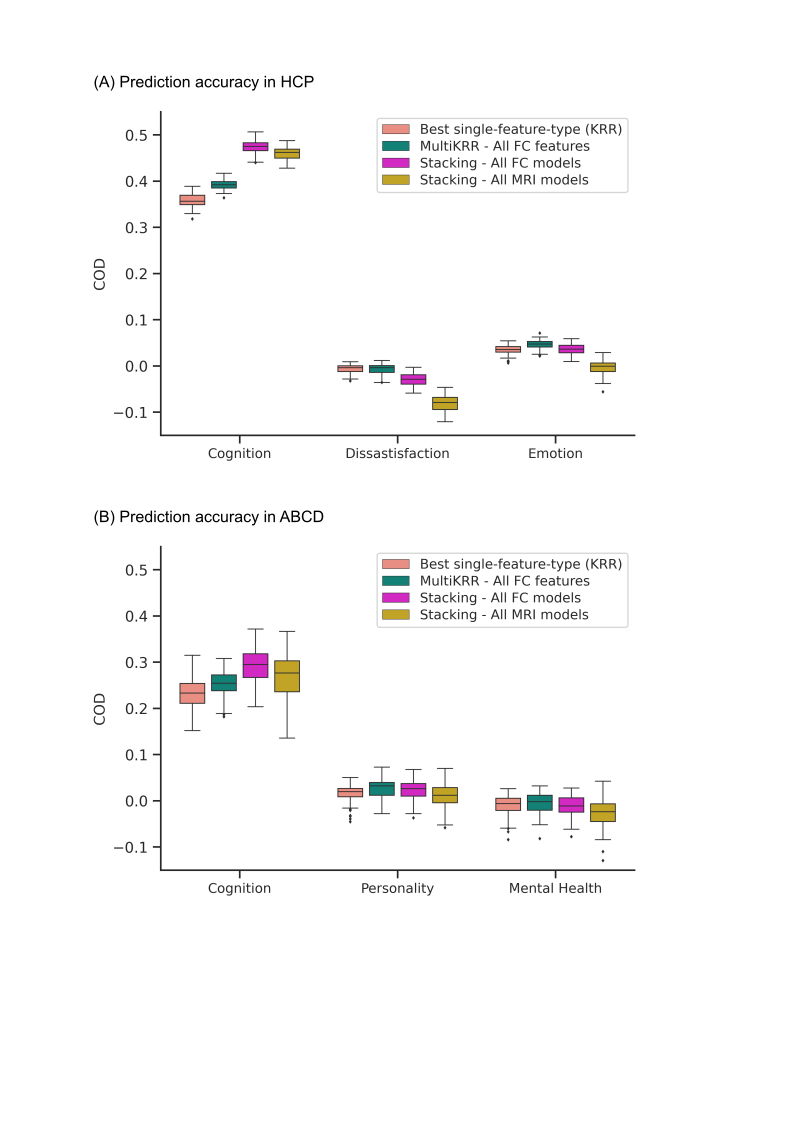

**Figure S16.** Combining resting and task FC was as good as combining across all modalities. Figure is the same as Figure 7, except that COD was shown instead of Pearson’s correlation. (A) Prediction performance (COD) from combining various MRI features and modalities in the HCP dataset. We considered multi-KRR of all FC features, stacking of all FC models and stacking of all single-feature-type models across all modalities. For comparison, the best single-feature-type from KRR is shown. Each boxplot shows the distribution over 60 repetitions of the nested cross-validation procedure. (B) Prediction performance (COD) from combining various MRI features and modalities in the ABCD dataset. We considered multi-KRR of all FC features, stacking of all FC models and stacking of all single-feature-type models across all modalities. For comparison, the best single-feature-type from KRR is shown. Each boxplot shows the distribution over 120 repetitions of the nested cross-validation procedure.
